# Mechanical fidelity of brain tissue phantoms: A comparative experimental characterization of equilibrium and dissipative behavior

**DOI:** 10.64898/2026.10.01.755978

**Authors:** Elisa Baldoin, Manuel P. Kainz, Nicola Petrone, Piero G. Pavan, Gerhard A. Holzapfel, Michele Terzano

**Affiliations:** Department of Industrial Engineering, University of Padova, Padua, Italy; Institute of Biomechanics, Graz University of Technology, Graz, Austria; Fondazione Istituto di Ricerca Pediatrica Città Della Speranza, Padua, Italy; Department of Structural Engineering, Norwegian University of Science and Technology, Trondheim, Norway

**Keywords:** brain tissue, tissue phantom, mechanical characterization, inelastic material behavior, head model

## Abstract

Despite lacking a dedicated structural function, brain tissue is exposed to severe mechanical loading during accidents and trauma, with potentially permanent or lethal consequences. Therefore, characterizing its mechanical behavior assumes particular importance for designing protective equipment, planning surgeries, and training neurosurgeons, tasks that often rely on physical brain replicas made from phantom materials. Although the mechanics of brain tissue has been extensively studied, the selection of suitable phantom materials still relies on simplified metrics that fail to capture the highly nonlinear and rate-dependent behavior of the brain. In this work, we characterize the mechanical response at low-to-medium strain rates of common synthetic brain phantom materials, including Sylgard 527, Ecoflex Gel, and a composite hydrogel, and establish rigorous comparisons with porcine brain tissue as a benchmark. To this end, we employ a multistep stress relaxation protocol under both uniaxial extension and compression across two different strain rates, introduce thermodynamic metrics, and perform comparisons based on rigorous statistical analysis. In particular, we isolate the thermodynamic equilibrium stress-strain curves, representing the baseline material’s response, from rate-dependent dissipative inelastic effects, relatable to viscoelastic and poroelastic mechanisms. Our results provide a thorough comparison, revealing critical limitations in existing simulant materials and highlighting significant space for developing improved brain surrogates. The comprehensive dataset collected can also support the calibration of constitutive models, enhancing the predictive accuracy of numerical simulations involving brain tissue and its surrogates. More broadly, this study highlights the need to move beyond simplistic, scalar-metric-based comparisons of mechanical responses across materials.

## 1. Introduction

The mechanical characterization of human brain tissue is a crucial step in the production of physical head phantoms featuring a phantom material that mimics the behavior of the brain [1–3]. Such replicas provide valuable tools for neurosurgical education and training, surgical planning, and protective equipment testing [4–7]. Traditional approaches in neurosurgery involve using human cadavers, which are associated with logistics issues, high costs, and ethical concerns. On the other hand, platforms based on digital simulations and virtual reality lack realistic haptic feedback and are often incompatible with surgical instruments. Anthropomorphic phantoms have therefore emerged as a promising alternative, enabling realistic anatomical geometries and tactile feedback through suitable material selection [4, 5]. In this context, identifying materials capable of replicating the *mechanical* behavior of brain tissue is essential. In the following, we simply refer to these materials as brain phantoms or simulants, but keep in mind that only mechanical similarity is addressed.

In the field of traumatic brain injury (TBI) biomechanics, the protection effectiveness of helmets has recently been evaluated through experimental tests on headforms such as the Hybrid III and EN960, which provide input kinematics for a human head finite element (FE) model to assess strain-derived injury metrics [8]. Within this framework, the limited biofidelity of rigid headforms has led to the development of physical replicas that feature brain simulants [7, 9–12]. In parallel, human head FE models are validated using outdated and rare cadaveric data, which are subjected to potential biases related to hydration, age and health conditions. To mitigate these shortcomings, the use of biofidelic instrumented head replicas, coupled with their corresponding FE models, could improve consistency between experimental measurements and numerical predictions [13]. Notably, the success of this methodology ultimately hinges on the accurate characterization and constitutive modeling of both brain tissue and simulant materials.

The brain is a highly heterogeneous organ, composed of gray matter, which consists mainly of neuronal cell bodies, and white matter, which contains axonal fibers, astrocytes, and microglial cells. In addition, the fluid stored in cells and the interstitial fluid constitute a substantial fraction of the volumetric content of the brain [14]. Macroscopically, the brain exhibits a highly nonlinear and strongly rate-dependent mechanical response [15], which is influenced by both fluid-related mechanisms associated with interstitial fluid flow [16–18] and intrinsic viscoelasticity of the solid tissue matrix [19]. As a result, the observed mechanical behavior arises from the combined contribution of multiple constitutive mechanisms, involving inelastic effects associated with viscoelastic and poroelastic dissipative phenomena.

Despite the large number of experimental and numerical studies published over the past decades, the mechanical characterization of brain tissue remains challenging [20]. Numerous experimental investigations have addressed the response of brain tissue under compression [21–31], tension [32–36], or shear [17, 37–42]. However, experimental data reported in the literature often show significant variability, with the inferred mechanical properties differing by orders of magnitude across different studies [2, 43]. Such discrepancies may be due to differences in specimen preparation and testing protocols, among other factors. This poses a significant challenge for the development of constitutive models, whose predictive capability strongly depends on the accuracy of the underlying experimental data.

Furthermore, the theoretical framework for modeling the mechanical behavior of the brain remains a matter of discussion. Evidence of the rate-dependence of the stress-strain curve of brain tissue is overwhelming [20]; however, several computational studies still treat the brain as a purely hyperelastic material [17, 44–51]. This can be partly explained by the lack of established experimental procedures capable of isolating the equilibrium response of the material from dissipative contributions. In turn, this affects the ability to properly calibrate and validate rate-dependent constitutive models. In principle, the equilibrium response of a material can be approximated by assuming quasi-static loading rates; in practice, this is often infeasible [49, 52]. As an alternative, some studies have proposed averaging the loading and unloading stress–strain curves obtained from cyclic tests [17, 49], although this method lacks rigorous mechanical foundations and may lead to inaccurate model predictions [53]. Generally, rate-dependent material behavior is characterized by means of single- and multi-step stress relaxation experiments. In single-step relaxation tests, several specimens are required to determine the equilibrium response at different strain levels. This limitation can be addressed through multi-step relaxation tests, also known as stepwise relaxation tests. This procedure involves applying a series of increasing strain levels to a single specimen, each followed by a holding phase to allow stress relaxation. Although stepwise stress relaxation protocols are commonly used in the mechanical characterization of polymers, rubber-like materials, and hydrogels [54–69], they are not widely adopted in biomechanics [52, 70, 71].

Studies addressing the comparative mechanical characterization of synthetic materials used for brain phantoms have employed different experimental methods, including quasi-static and cyclic loading, stress-relaxation tests, and dynamic mechanical analysis [2, 31, 72–74]. Forte et al. [2] compared Phytagel (PHY), gelatin gels, Polyvinyl Alcohol (PVA), Sylgard 184, Sylgard 527, and a PVA–PHY composite hydrogel (CH) with porcine brain tissue under unconfined compression in air, identifying CH as the most suitable candidate. Further indentation-relaxation and frequency-sweep tests supported its feasibility as a brain surrogate [2]. Navarro-Lozoya et al. [31] assessed silicone gels, gelatin, gelatin-agarose hydrogels, and emulsions against porcine brain tissue under unconfined compression and stress relaxation, with a gelatin–agarose hydrogel and an emulsion showing the closest compressive response, although none fully reproduced brain tissue relaxation. Singh et al. [72] investigated bovine and agarose gelatins and Sylgard 527 alongside porcine brain tissue under quasi-static and sinusoidal compression, showing better agreement for 3% and 5% bovine gelatin under quasi-static loading and for agarose gels and Sylgard 527 under sinusoidal loading. Distler et al. [73] tested alginate–gelatin hydrogels against porcine brain tissue under cyclic and stress-relaxation loading in compression, tension, and shear, showing their ability to reproduce characteristic nonlinear, conditioning, and compression–tension asymmetric behavior. Kainz et al. [74] compared the mechanical response of the CH introduced by Forte et al. [2] to porcine brain tissue using a cyclic indentation-relaxation protocol with a flat indenter, and observed close agreement in terms of total reaction force over time. Across these studies, the following metrics have been identified for comparison: apparent elastic modulus, peak stress, complex, storage, and loss moduli, and normalized stress decay. Nonetheless, these metrics do not explicitly isolate the inelastic mechanisms governing the material behavior, thereby preventing a consistent comparison of equilibrium and dissipative responses between brain tissue and simulant materials.

This study introduces an experimental methodology to characterize the mechanical response of brain tissue, and employs it to perform a consistent comparison between native tissue and selected simulant materials for brain phantoms. The mechanical response of porcine brain tissue is established as a benchmark, using a multistep stress relaxation protocol designed to separate the equilibrium response from the inelastic effects associated with rate-dependent dissipative phenomena. In a purely mechanical setting, the equilibrium response of a material is represented by an *equilibrium relation*, that is, a stress-strain curve describing the material behavior at steady state [75]. Physically, it describes the long-term mechanical response of brain tissue, which is relevant to modeling slow processes, such as tumor-induced tissue deformation. Rate-dependent dissipative phenomena, by contrast, characterize the transient response of the material and correspond to viscous effects such as stress relaxation and creep. In practice, these play an important role in multiple scenarios, including intra-operative brain shift, surgical tool insertion, haptic-feedback-based neurosurgical simulation, and impact scenarios for TBI applications. In addition, separating equilibrium and rate-dependent dissipative behavior is beneficial for calibrating constitutive models, where rate-dependent contributions are superposed onto a base, often hyperelastic, constitutive function.

The same experimental protocol is applied to three polymeric materials already proposed as simulant materials for brain phantoms: Sylgard 527, Ecoflex Gel, and the CH introduced by Forte et al. [2] and prepared following the modified protocol presented by Kainz et al. [74]. The inelastic dissipative behavior is quantified by means of macroscopic metrics defined in line with with Bogoni et al. [52], including the stress work associated with the equilibrium response and the accumulated rate-dependent dissipation during both loading and unloading. These measures are combined with more conventional metrics, such as the normalized stress decay, to provide a comprehensive comparison between brain tissue and simulant materials. The experimental investigation is carried out under both uniaxial extension and compression, at two different loading rates. This allows the influence of loading mode and rate on both the equilibrium and inelastic responses to be thoroughly evaluated and compared across materials.

## 2. Materials and methods

This section describes the theoretical framework underpinning the proposed mechanical characterization (Sect. 2.1), the experimental methods, including preparation of biological and synthetic specimens, description of the experimental setup, definition of the experimental protocols, and data analysis (Sect. 2.2), and the quantitative metrics employed for comparison across different materials (Sect. 2.3).

### 2.1 Theoretical considerations

The mechanical characterization of a material is performed under controlled conditions, which define the experimental protocol, and assuming that each specimen tested is representative of the material’s behavior under those conditions and at the length scale of interest. The constitutive theory of materials provides the framework for interpreting the experimental results and for formulating constitutive equations that are expected to describe the material response beyond the specific test conditions. In practice, little is known about the explicit form of the constitutive relations before testing, which may depend on the history of deformation, temperature, and additional internal variables [75]. Therefore, attempting a mechanical experiment without first formulating physically motivated assumptions on the materials under investigation appears infeasible.

To avoid unnecessary complications, all materials considered in the present study are assumed to be *local* and to undergo isothermal processes. Accordingly, the constitutive response at a material point ***X*** and time *t* is solely determined by the local state and its history [76]. The material history prior to the beginning of each experiment is neglected. Furthermore, we state that the constitutive behavior does not explicitly depend on ***X***, i.e., that each single specimen is *homogeneous*. This assumption is particularly amenable, since it allows, at least theoretically, to produce spatially homogeneous stress and deformation fields through suitably prescribed boundary conditions. In homogeneous solids, for instance, the constitutive response can be directly inferred from global force and displacement measurements, although care must be applied to ensure that homogeneity is actually achieved in experimental practice [70]. However, some of the materials considered in the present study are more appropriately described as biphasic fluid-saturated materials rather than as standard homogeneous solids. In such materials, the achievement of homogeneous fields is generally not possible using standard boundary conditions of mechanical testing, such as prescribed force or platen displacement. Moreover, the dissipative behaviour of a biphasic material cannot, in general, be assessed from deformation measurements alone. These aspects and their implications for the interpretation of the mechanical experiments are discussed below.

#### 2.1.1. Unconfined compression and tension

A specimen of brain tissue at the macroscale can be viewed as a mixture of cellular and extracellular solid constituents and interstitial fluid. When immersed in a prescribed fluid bath, the specimen exchanges fluid with its environment, eventually reaching a new thermodynamic equilibrium. The resulting swollen configuration is here adopted as the *reference* configuration for the mechanical experiment. This is a cylinder of reference height *H*_0_ and radius *R*_0_, in which material points are parametrized by cylindrical coordinates 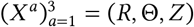 with respect to an orthonormal basis (***e***_*r*_, ***e***_*θ*_, ***e***_*z*_). The origin is placed at the center of the bottom surface of the cylinder, and ***e***_*z*_ points upwards along its axis. Mechanical loading is applied by prescribing a time-dependent axial displacement of the surface *Z* = *H*_0_ through the controlled motion of a rigid, impermeable platen whose area is much larger than the cross-sectional area of the specimen. The surface *Z* = 0 rests upon an analogous platen, while the lateral surface *R* = *R*_0_ is free to expand or contract and remains in contact with the bathing fluid. A load cell measures the reaction force at the top platen in the axial direction. This setup corresponds to the classical realization of an *unconfined* compression or tension experiment [77], see Fig. 1.

**Figure 1.**
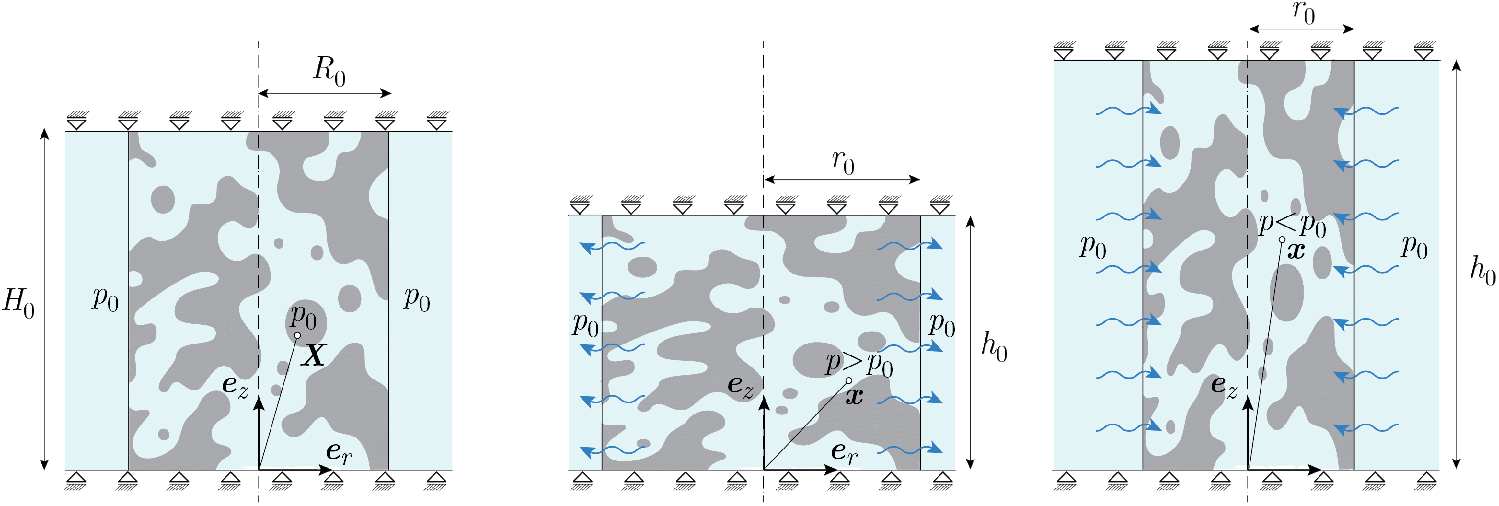
Sketch of an unconfined compression and tension experiment, showing the radial-axial plane defined by the basis vectors (***e***_*r*_, ***e***_*z*_). A cylinder of reference height *H*_0_ and radius *R*_0_, initially in equilibrium with a fluid at pressure *p*_0_, is subject to a prescribed deformation along ***e***_*z*_. In the current configuration, of height *h*_0_ and radius *r*_0_, a pressure gradient develops in the radial direction, driving fluid inwards or outwards, until a new equilibrated state is established.

We assume an axisymmetric kinematic ansatz in which cross-sections normal to the cylinder axis undergo spatially homogeneous axial deformation. Under this assumption, the deformation field is independent of the axial coordinate. Experimentally, this is achievable only when the contact with the platens is frictionless [77]; in practice, we are neglecting end effects, a common assumption in mechanics which we adopt without further verification. The motion *χ* (***X***, *t*) can therefore be written as [78]^1^

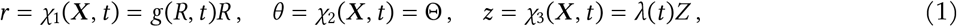

where 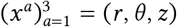 are the coordinates in the *current* configuration, *λ*(*t*) is the imposed axial stretch, and *g*(*R, t*) = *r*(*R, t*)/*R* is a radial deformation function.

The deformation gradient associated with the motion of Eq. (1) takes the following diagonal representation

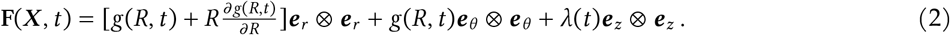

Assuming isotropy, we expect a diagonal structure of the Cauchy stress tensor **T**(***x***, *t*) in the cylindrical coordinate system, namely,

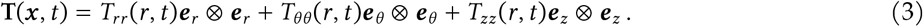

The experimentally accessible kinematic quantity is the stretch *λ*(*t*), which can be directly computed from the prescribed axial displacement *u*_*z*_(*t*):

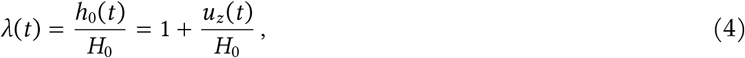

where *h*_0_(*t*) is the current height. A constant platen velocity is enforced via a prescribed stretch rate 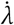, i.e., 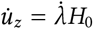.

The axial reaction force *f*_*z*_(*t*), recorded by the load cell at *h*_0_(*t*), is the resultant of the axial component of the stress tensor over the upper surface. Accordingly, we can write

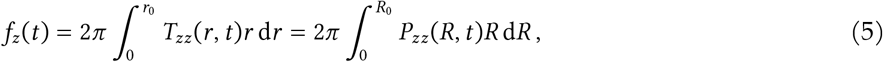

where *r*_0_(*t*) = *g*(*R*_0_, *t*)*R*_0_ is the current specimen radius and *P*_*zz*_(*R, t*) is the axial component of the nominal stress tensor **P** = *J* **TF**^−⊤^. From the measured force we can compute the nominal stress metric

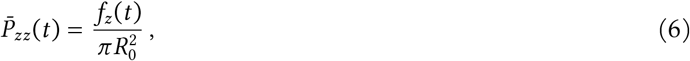

which represents an ‘area average’ of the axial component of the nominal stress tensor.

At this point, we should already recognize several limitations of the experimental characterization. First, the stress fields in the sample may depend on the radial coordinate and cannot be inferred from the measured axial force alone. Their determination requires the solution of the initial boundary value problem (IBVP) associated with unconfined uniaxial deformation. The role of spatial inhomogeneity, as well as particular conditions under which a homogeneous uniaxial solution is recovered, are discussed in Appendix A. Second, the definition of the reference configuration assumed in the experiments requires careful consideration. Tissue specimens are harvested post-mortem, dissected from whole brains, and mounted on the testing apparatus. Although care is taken to maintain hydration throughout these operations, the specimens require time to equilibrate after mounting. Residual stresses and osmotic effects might develop throughout this process, implying that the reference configuration is not necessarily stress-free. As described in Sect. 2.2, the fluid medium was selected to minimize tissue swelling. In addition, the reaction force during the equilibration period was recorded by the load cell of the testing apparatus. Stabilization of this force provides an experimental indication that the mounted specimen has approached a stationary state. In principle, the subsequent experiment could then be interpreted as an incremental loading process about a pre-stressed equilibrium configuration. In practice, a single reaction force is insufficient to determine the complete initial stress field without additional assumptions or measurements. The treatment of this initial stress state is beyond the scope of the current study.

#### 2.1.2. Equilibrium response and dissipative behavior

To interpret the mechanical behavior of a material, it is beneficial to isolate its equilibrium response from dissipative contributions. Following the framework presented by Bogoni et al. [52], the experimental measurements are intended to access the equilibrium relation of the material, defined as ‘the set of strain processes and equilibrium stresses, attainable by means of relaxation processes’ [58]. These concepts motivate the experimental protocols described in Sect. 2.2.2 and the metrics introduced in Sect. 2.3. Importantly, these metrics enable a comparative assessment of the materials without imposing a predefined constitutive model on the experimental data.

To make further progress, it is useful to briefly revisit the thermodynamics of biphasic materials, with particular reference to experimentally accessible quantities. Let us start from the local dissipation inequality, expressed per unit current volume^2^:

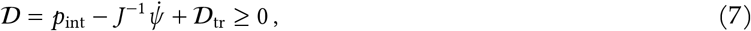

where *D* is the dissipated power, *ψ* is the strain-energy function of the solid, and *D*_tr_ accounts for transport-related dissipation. In the context of fluid-saturated solids, *ψ* is understood as an energy per unit reference volume of the solid. The stress power *p*_int_ is the contraction between an effective stress tensor and its work-conjugate rate of deformation [79].

With respect to inequality (7), a state of thermodynamic equilibrium entails both 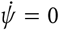 and *p*_int_ = 0 (since the rate of deformation is null during relaxation). In addition, the transport related dissipation *D*_tr_ is also null (since the fluid flow ceases). The condition *D* = 0 associated with such a state is local to the current time; in other terms, this state can still be path-dependent (for instance, due to material’s damage), resulting in an equilibrium hysteresis when loading and unloading is performed. Crucially, as observed by Bogoni et al. [70], this response can also be influenced by the strain rate applied during the loading phases.

Experimentally, not all terms appearing in inequality (7) are directly accessible. In particular, we observe that: (i) the stress power depends on the local stress and deformation fields; (ii) the strain-energy function remains unknown unless a constitutive equation is specified; and (iii) the transport-related dissipation cannot be determined without knowledge of the fluid flux. Considering the first limitation, the experimental setup allows to compute the volume-averaged nominal stress power as

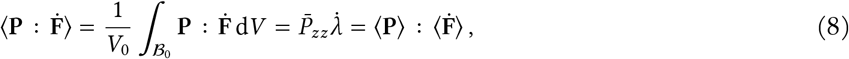

where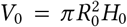 is the volume of the solid in its reference configuration *B*_0_. This is a consequence of the Hill–Mandel identity and it holds for inhomogeneous fields [80]. However, it is related to the total nominal stress and not to the effective stress power *p*_int_. The volume-averaged effective stress power therefore differs from Eq. (8) by a term related to the interstitial pressure (see Appendix B).

Regarding the second limitation, we follow the approach introduced by Bogoni et al. [52] and define an accumulated dissipation with respect to the equilibrium curve, such that the time-integrated contribution of the strain-energy function cancels out. The volume-averaged accumulated dissipation relative to the equilibrium curve then reads

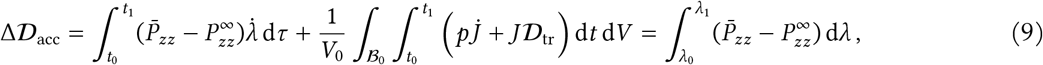

where 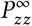 is the equilibrium nominal stress at given stretch and *p* is the interstitial fluid pressure. The integration is on an interval [*t*_0_, *t*_1_] on which *λ*(*t*) is assumed injective, such that the change of variables 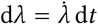 is allowed.

The last equivalence is a key result obtained under the assumptions of the unconfined uniaxial deformation, with additional restrictions; see Appendix B for its full derivation. In the remainder of this paper, the subscripts referring to the axial component are dropped, having clarified that this is the only force component accessible experimentally.

### 2.2. Mechanical characterization

#### 2.2.1. Sample preparation

Porcine brain tissue was obtained from a local slaughterhouse in the form of hemispheres and tested within 12 h upon acquisition. To preserve tissue integrity and hydration, the hemispheres were immersed in phosphatebuffered saline (PBS) solution and refrigerated at 3 ^°^C until required for testing. Each specimen was prepared immediately before the corresponding experiment. The cerebellum and medulla oblongata were first removed, after which a coronal section was extracted from each hemisphere using PBS-wetted blades (Fig. 2a–b). Individual specimens were then obtained from the corona radiata at the interface with the cerebral cortex using a stainless-steel biopsy punch (Stiegelmeyer GmbH & Co. KG) with an inner diameter of 8 mm (Fig. 2b, dashed circle). As shown in Fig. 2c, this sampling strategy produced specimens composed predominantly of white matter, with a smaller contribution of gray matter. Owing to the extremely soft nature of brain tissue, the samples were susceptible to deformation under their own weight. Therefore, the diameter of each specimen was measured immediately after extraction using a caliper to account for potential deviations from the nominal punch diameter. The specimen thickness was instead inferred optically using a video extensometer, as described by [81]. Across all samples (*n* = 10 per combination of loading mode and strain rate), the diameter and thickness were 7.7 ± 0.4 mm and 5.1 ± 0.3 mm, respectively (mean ± standard deviation).

**Figure 2.**
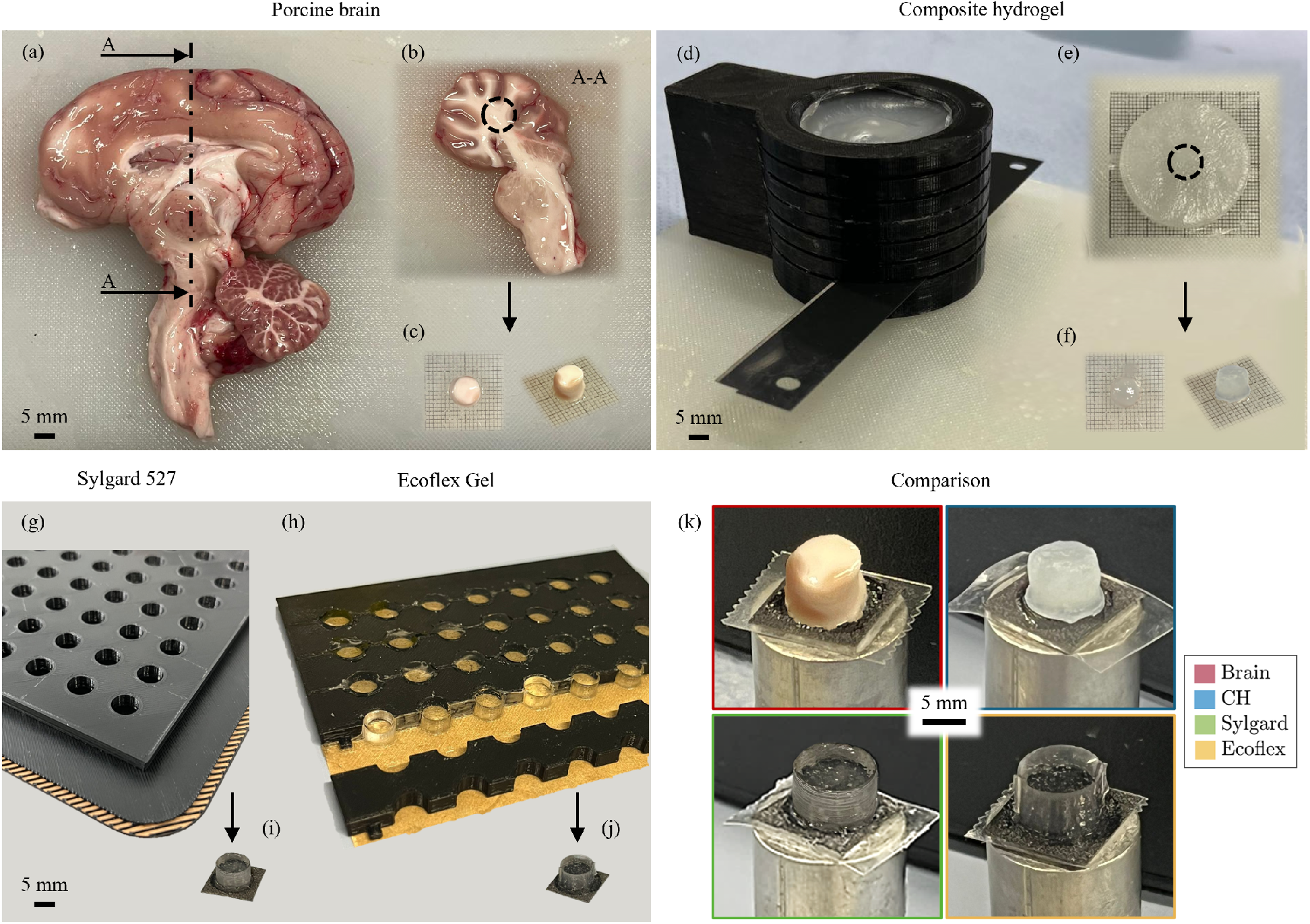
Sample preparation for porcine brain tissue (a–c), composite hydrogel (CH) (d–f), Sylgard and Ecoflex (g–j), with a detailed view of the mounted samples prior to testing (k). In detail: (a) Brain hemisphere; (b) brain coronal section with identification of the sampling region; (c) brain samples; (d) slicing device used for CH; (e) CH section with identification of the sampling region; (f) CH samples; (g–h) molds used for Sylgard and Ecoflex, respectively; (i–j) Sylgard and Ecoflex samples, respectively; (k) brain (red), CH (blue), Sylgard (green), and Ecoflex (yellow) samples prior to testing. This color scheme is consistently adopted when comparing the materials in the results section.

As previously mentioned, three simulant materials were also tested: a composite hydrogel (described in Kainz et al. [74] and hereafter referred to as ‘CH’), a silicone dielectric two-part gel (known by its commercial name Sylgard 527 and hereafter referred to as ‘Sylgard’), and a platinum silicone rubber (known by its commercial name Ecoflex Gel and hereafter referred to as ‘Ecoflex’). The specimens were prepared using 3D-printed polylactic acid (PLA) tools. Their diameter and thickness were measured using a caliper. For CH, a cylindrical batch of polymerized material was placed inside a hollow cylindrical mold with evenly spaced slits (Fig. 2d). This mold was specifically designed to drive blades wetted with deionized water (DIW) and produce slices with a nominal thickness of 5 mm. Individual specimens were subsequently extracted from these slices using the same biopsy punch for the brain tissue (Fig. 2e, dashed circle). Across all CH samples (Fig. 2f; *n* = 10 per combination of loading mode and strain rate), the diameter and thickness were 7.6 ± 0.4 mm and 4.8 ± 0.3 mm, respectively.

The adhesive nature of Sylgard and Ecoflex precluded the slicing procedure used for CH. Therefore, custom molds were designed to cast specimens directly with a nominal diameter of 8 mm and a thickness of 5 mm. For Sylgard, a mold comprising through-holes and a temporary base (raft) was used to prevent material leakage during curing (Fig. 2g). After curing, removal of the base left the cavities open at both ends; consequently, the mold could not be reused for subsequent casting. In contrast, Ecoflex was cast using a reusable mold consisting of an outer housing and multiple removable inner inserts arranged to form an array of cylindrical cavities. The base was lined with baking paper, which served as a temporary bottom surface during casting. After curing, the inserts were disassembled by sliding them off the base (Fig. 2h), allowing straightforward specimen removal and reuse of the mold components. For both Sylgard and Ecoflex, the materials were prepared by mixing the two components at a 1A:1B ratio and poured into the mold cavities. Sylgard was then cured at room temperature for seven days, whereas Ecoflex required approximately two hours. Because of their adhesive features, the cured specimens were glued directly to the sandpaper used during testing with cyanoacrylate glue (Fig. 2i–j). The diameter was 8 mm for all Sylgard and Ecoflex samples (*n* = 5 and *n* = 10, respectively, per combination of loading mode and strain rate), while the thickness was 5 mm for Ecoflex and 4.83 ± 0.08 mm for Sylgard.

#### 2.2.2. Experimental setup

The tests were conducted using a triaxial testing device previously introduced in Sommer et al. [82]. Although the present experiments only involved uniaxial compression and tension, this apparatus was selected because its load cell has a full-scale capacity of ±2 N, making it suitable for the characterization of ultra-soft materials such as brain tissue. Before testing, each specimen was mounted onto a sandpaper-covered platen using cyanoacrylate adhesive. Figure 2k shows representative specimens of the four materials mounted on the upper adapter before placement into the testing device. At the start of each test, the upper adapter holding the specimen was lowered towards a lower adapter, also coated with sandpaper and cyanoacrylate adhesive. The approach ended when the load cell recorded a compressive force of 5 mN. The setup at the beginning and end of this approach procedure for a brain specimen is shown in Fig. S1a–b of the Supplementary Material. Following contact with the lower adapter, the specimen was held between the adapters to allow the adhesive to cure before the loading protocol was applied. A holding time of 2 min was found to be sufficient for brain tissue and CH, whereas 20 min was required for Sylgard and Ecoflex.

All specimens were tested while submerged in a bath to maintain hydration and provide consistent testing conditions. PBS was used for brain tissue, whereas DIW was used for the simulant materials. Different media were used for brain tissue and CH to avoid osmotic effects that could otherwise alter their mechanical response. For the hydrophobic polymeric materials, Sylgard and Ecoflex, the surrounding acqueous medium was not expected to affect their mechanical behavior. These materials were therefore tested in DIW, consistent with the conditions used for CH. Once the adhesive had cured, the appropriate fluid was poured into the glass basin until the specimen was fully submerged (Fig. S1c, Supplementary Material). Testing in air tests was avoided to maintain comparable boundary conditions across all materials and to minimize potential differences associated with buoyancy effects.

#### 2.2.3. Multi-step stress relaxation protocols

Based on the considerations developed in Sect. 2.1.2, we identify multi-step stress relaxation experiments, incorporating both loading and unloading paths at various strain rates, as the most suitable experimental framework to characterize the equilibrium response as well as the dissipative behavior of the material. Specifically, two multi-step stress relaxation protocols were defined: one in compression and one in tension. Both protocols were first calibrated using brain tissue through preliminary tests aimed at minimizing the overall testing duration while ensuring that stress relaxation occurred within the prescribed holding periods. The duration of each holding phase was established according to the quantitative and qualitative relaxation criteria described in Appendix C. Once the protocols had been established using brain tissue as the reference material, they were applied unchanged to the simulant materials.

Both protocols were displacement-controlled and consisted of five incremental stretch levels, followed by holding phases. In compression, the axial stretch was progressively reduced to *λ*_min_ = 0.85, whereas in tension it was progressively increased to *λ*_max_ = 1.15. Before the loading sequence, an initial holding period of 10 min was imposed at *λ* = 1 to allow for mechanical stabilization of the specimen after immersion in the fluid bath. Loading and unloading were discretized into equally spaced stretch increments of Δ*λ* = 0.025, except for the first increment, which was Δ*λ*_start_ = 0.05. The prescribed deformation was applied through displacement-controlled ramps at two constant stretch rates, 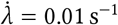 and 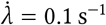, hereafter referred to as the ‘low’ and ‘high’ stretch rates, respectively. For compression, the holding times varied among stretch levels during both loading and unloading, as summarized in Table 1. In tension, a constant holding time of 700 s was sufficient to satisfy the relaxation criterion at all stretch levels during both loading and unloading. The final compression and tension protocols are illustrated in Fig. 3a–b.

**Table 1.**
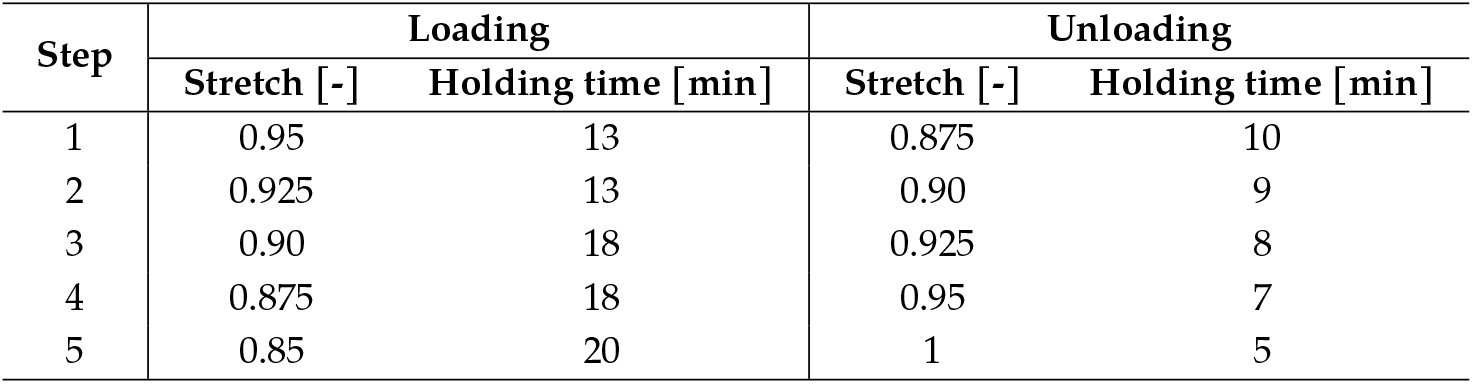
Holding times adopted at each stretch level during loading and unloading steps in the final multi-step stress relaxation protocol in compression.

**Figure 3.**
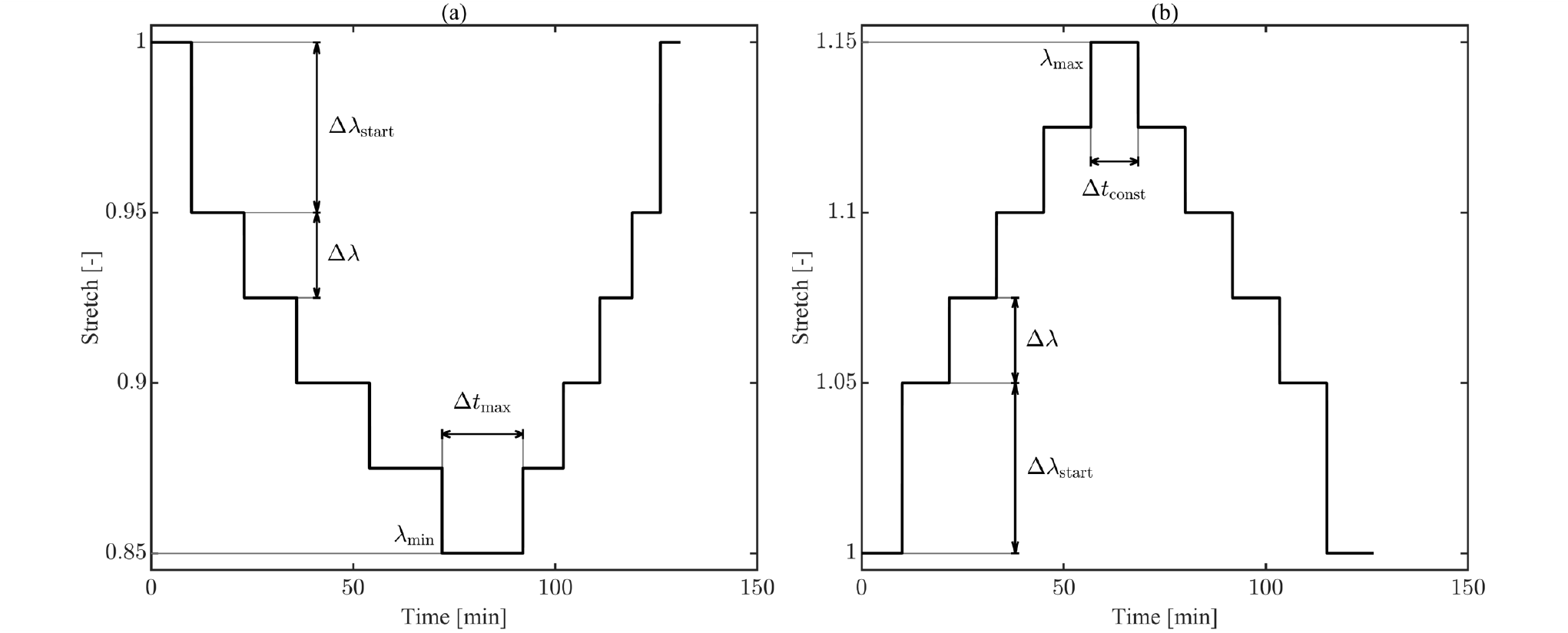
Final multi-step stress relaxation protocols adopted for compression (a) and tension (b). For compression, variable holding times were assigned to loading and unloading steps (the maximum duration is indicated as Δ*t*_max_). For tension, a constant holding time (Δ*t*_const_ = 700 s) was set for all steps.

Representative specimen configurations during compression are shown in Fig. 4 for each material. The images show the specimens in their initial configurations, *λ* = 1, and at the maximum applied deformation, corresponding to *λ*_min_ in compression and *λ*_max_ in tension.

**Figure 4.**
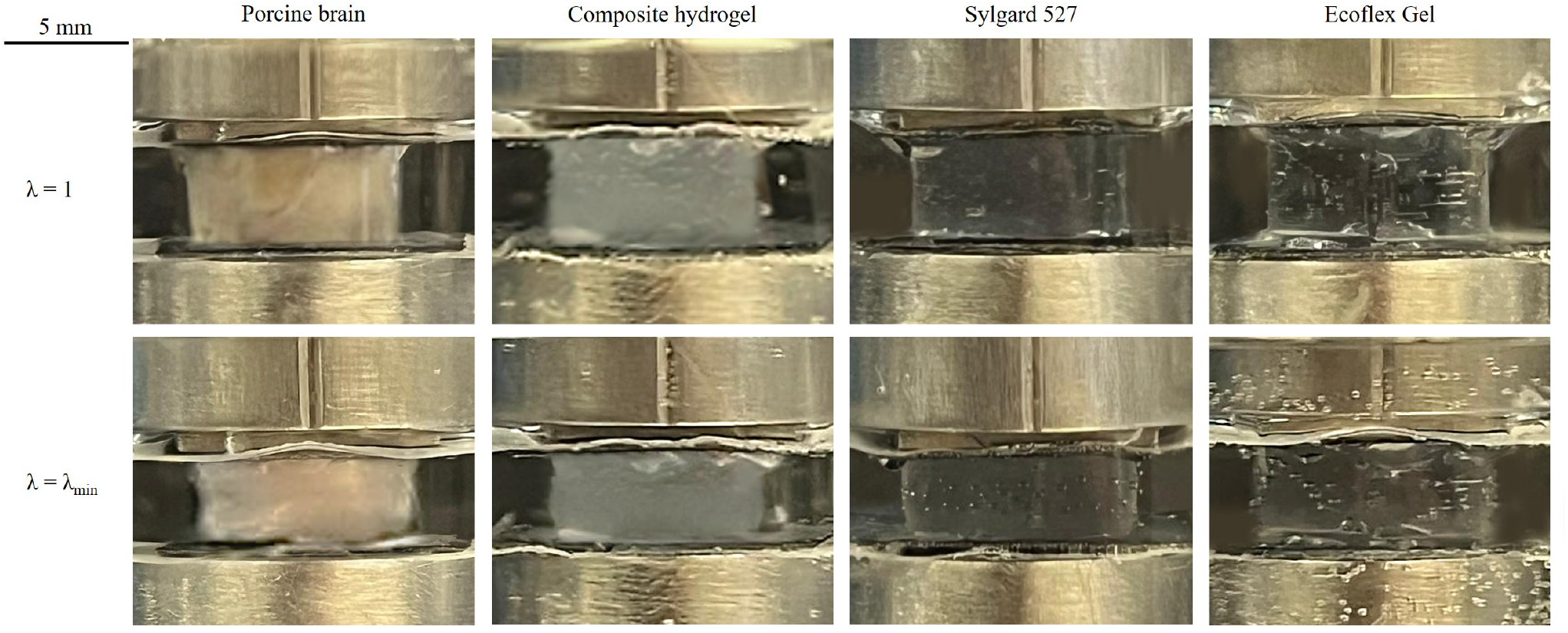
Representative configuration of the four tested materials during compression tests. The first row refers to the undeformed configuration (*λ* = 1), while the second one refers to the configuration under maximum compression (*λ* = *λ*_min_ = 0.85). The corresponding configurations during tension tests are provided in Fig. S2 of the Supplementary Material available online.

#### 2.2.4. Data analysis: acquisition and processing

The acquired signals comprised time *t*, displacement *u*(*t*), and force *f* (*t*). Data were recorded at a sampling frequency of 20 Hz during the initial holding period after specimen submersion, 100 Hz during each loading or unloading ramp and during the first 60 s of relaxation following each stretch increment, and 5 Hz for the remainder of each holding phase. This variable sampling strategy was adopted to accurately capture the transient response immediately following the onset of stress relaxation while limiting the volume of data acquired during the later stages of the holding periods. The raw signals were processed in Matlab 2023b (The MathWorks Inc.). Specifically, the data from all runs were interpolated onto a common synthetic time vector defined separately for each stretch rate and loading condition. This procedure ensured consistent temporal discretization across all runs performed under a given testing condition. Given the sampling frequencies ranging from 5 to 100 Hz, the influence of interpolation on the reconstructed signals was considered negligible. The processed signals were subsequently used to extract equilibrium and inelastic metrics, enabling a quantitative comparison between each simulant material and brain tissue, as described in Sect. 2.3.

Owing to the limited sample size for each loading-rate condition and the difficulty of reliably assessing normality, non-parametric statistical tests were used throughout the study. Statistical analyses were performed in Matlab using a significance level of *α* = 0.05. For each material, the influence of the effect of stretch rate on the equilibrium, energy-based, and normalized relaxation metrics was assessed using the Mann–Whitney U (MWU) test. Difference in accumulated dissipation between the loading and unloading phases within the same testing condition were evaluated using a one-tailed Wilcoxon signed-rank (WSR) test. Finally, comparisons between brain tissue and simulant materials were performed using the MWU test for each quantitative metric.

### 2.3. Quantitative metrics for simulant fidelity

The multi-step stress relaxation protocols adopted in this study enable the equilibrium and dissipative contributions to the mechanical response to be assessed separately. On this basis, a set of quantitative metrics was defined to compare the mechanical behavior of simulant materials with that of brain tissue. The selected metrics capture complementary features of the response, including equilibrium stresses at prescribed deformation levels, as well as dissipative effects. The proposed framework allows the fidelity of each simulant relative to brain tissue to be evaluated according to the requirements of a specific application. For illustrative purposes, the metrics introduced in this section are shown using representative data collected from brain tissue in compression in Fig. 5. This representations provides an idealized representation of the mechanical response of the material and is intended solely to demonstrate the definition, extraction, and interpretation of the proposed metrics. The same framework was applied analogously to the tensile tests.

**Figure 5.**
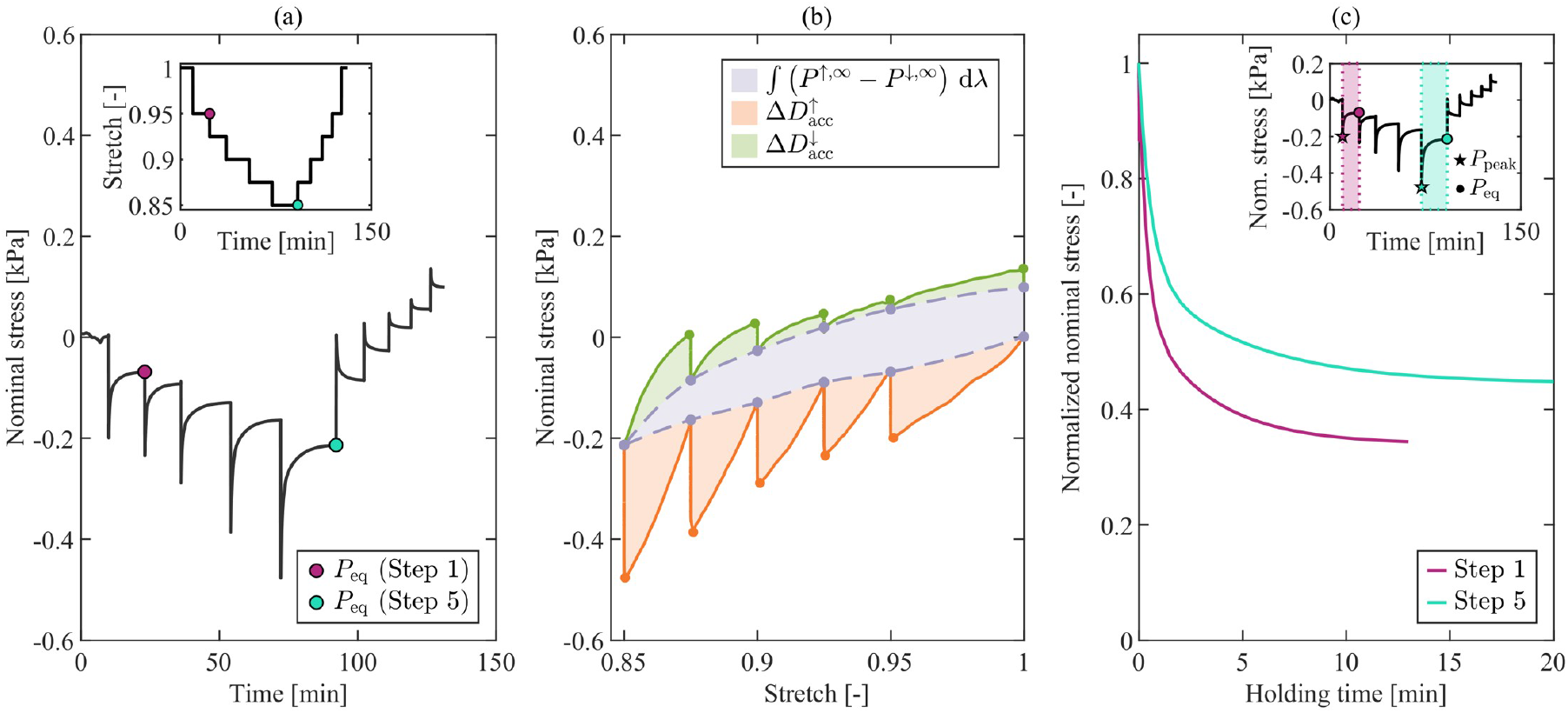
Quantitative metrics used to assess simulant fidelity, illustrated for a compression test on brain tissue at low rate. (a) Evolution of the nominal stress over time and stretch-controlled protocol, with equilibrium points at the end of the first (magenta) and fifth (aquamarine) loading steps; (b) accumulated dissipation during loading (orange area) and unloading (green area), and stress work associated with the equilibrium response (gray area); (c) stress decay during the first (magenta) and fifth (aquamarine) loading steps. The corresponding steps are highlighted in the inset, which shows the nominal stress–time response. Star and circular markers denote the peak nominal stresses *P*_peak_ and equilibrium nominal stresses *P*_eq_, respectively, for the corresponding loading steps.

#### 2.3.1. Equilibrium response

Equilibrium stresses *P*_eq_ were identified at the end of each holding phase, corresponding to the attainment of a relaxed state at the prescribed stretch level. Two representative equilibrium points were selected to compare the response of each simulant with that of brain tissue:

i. the equilibrium stress at the end of the first loading step, corresponding to the lowest applied deformation level: *λ* = 0.95 in compression (magenta point in Fig. 5a) and *λ* = 1.05 in tension;
ii. the equilibrium stress at the end of the fifth loading step, corresponding to the highest applied deformation level: *λ*_min_ = 0.85 in compression (aquamarine point in Fig. 5a) and *λ*_max_ = 1.15 in tension.

Together, these quantities provide a concise yet informative description of the equilibrium response, enabling a comparison between materials independently of the dissipative contributions.

#### 2.3.2. Dissipative effects

To quantify dissipative effects, three energy-based metrics were chosen. The definition of these metrics is based on the framework proposed by Bogoni et al. [52] and adapted to the current experiment in an averaged sense, as explained in Sect. 2.1.2. In addition to the considerations and limitations already mentioned, we note that the calculation of an equilibrium stress–stretch curve is needed. Since equilibrium stresses were available only at the discrete stretch levels imposed during the multi-step protocol, the equilibrium curve was constructed by interpolating these points. Therefore, the calculated metrics also depend on the selected interpolation method.

Following from Eq. (9), the accumulated dissipation during loading and unloading can be expressed as

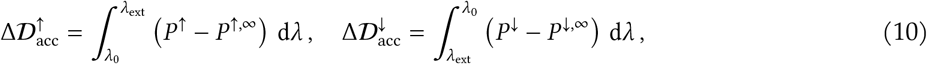

whereas the stress work associated with the equilibrium response is given by

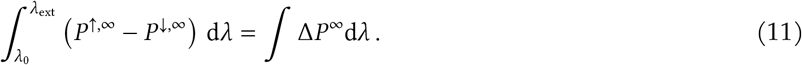

Here, *λ*_0_ = 1 denotes the stretch in the undeformed configuration, whereas *λ*_ext_ is the extreme stretch reached during loading, namely *λ*_max_ in tension and *λ*_min_ in compression. The quantities *P* ^↑^ and *P* ^↓^ denote the nominal stresses measured during loading and unloading, respectively; *P* ^↑,∞^ and *P* ^↓,∞^ denote the equilibrium stresses associated with each stretch level.

In the present study, the nominal stresses at the end of each holding phase (gray dots in Fig. 5b) were connected by piecewise linear interpolation to construct the loading and unloading equilibrium stress-stretch curves (dashed lines in Fig. 5b). The accumulated dissipation was then calculated as the area between the experimentally measured nominal stress–stretch curve and the corresponding equilibrium curve. This quantity was evaluated separately during loading (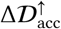, orange area in Fig. 5b) and unloading (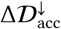, green area in Fig. 5b). The stress work associated with the equilibrium response, ∫ Δ*P*^∞^d*λ*, was calculated as the area enclosed between the loading and unloading equilibrium stress–stretch curves (gray area in Fig. 5b).

Together, these three quantities provide measures of the rate-dependent and path-dependent components of the mechanical response and may be particularly useful to support the selection of simulant materials for applications in which rate-dependent effects are important.

#### 2.3.3. Normalized stress relaxation

Although energy-based metrics offer a quantitative evaluation of the overall magnitude of inelastic effects, they do not fully capture the temporal evolution of stress relaxation. Therefore, the material response during each holding period of the loading phase was further characterized using normalized stress relaxation. Specifically, the nominal stress evolution over time was normalized by the peak stress reached at the end of the corresponding loading ramp:

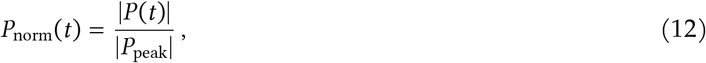

where *P* (*t*) is the nominal stress at time *t* and *P*_peak_ is the nominal stress at the beginning of the holding phase. Accordingly, *P*_norm_(*t*) decreases from unity at the beginning of the holding phase towards the normalized equilibrium stress |*P*_eq_| / |*P*_peak_| at the end of relaxation. As for the equilibrium response in Sect. 2.3.1, normalized stress relaxation was evaluated for two representative loading steps to compare each simulant material with brain tissue (lowest applied deformation, magenta curve in Fig. 5c, and highest applied deformation, aquamarine curve).

This metric describes the temporal stress decay throughout the selected holding phases, highlighted in magenta and aquamarine in the inset of Fig. 5c. Normalization with respect to the peak stress allows a direct comparison independently of the absolute stress level, thus highlighting the stress decay over time. To enable a quantitative comparison between each simulant and brain tissue, *P*_norm_ was evaluated at was evaluated at 1, 10, 30, 60, and 300 s after the onset of the holding phase, as well as at the end of the corresponding holding period. These time points were selected to capture both the initial stress decay following the loading ramp and the subsequent evolution towards equilibrium. Accurate characterization of stress relaxation is particularly relevant for applications involving time-dependent mechanical interactions, such as haptic-integrated neurosurgical simulators, where reproducing the temporal decay of the mechanical response is required to provide a biofidelic force feedback.

## 3. Results and discussion

This section presents and discusses the main experimental results, together with the statistical analyses used to compare the simulant materials with brain tissue according to the metrics introduced in Sect. 2.3. Results are reported separately for compression and tension. For each loading mode, the response of brain tissue is presented first as the reference benchmark, followed by the responses of the simulant materials. Data are reported as median values, with variability described by the interquartile range (IQR) and shown using error bars or shaded areas, as appropriate. For readability, only selected results are presented in this section. The remaining data are provided in the Supplementary Material available online, and specific supplementary plots are referenced where relevant.

### 3.1. Compression

#### 3.1.1. Brain tissue response

Figure 6a shows the median nominal stress as a function of time, together with the corresponding IQR, at high and low stretch rates (shown in red and blue, respectively). At both stretch rates, the nominal stress becomes positive during the unloading steps, indicating the development of tensile forces within the specimen. A comparison of the equilibrium stresses at identical stretch levels during loading and unloading shows that they do not coincide, indicating the presence of *equilibrium hysteresis*. This behavior is observed at both stretch rates and is further highlighted in Fig. 6b, which presents the reconstructed equilibrium stress–stretch relation obtained by interpolating the equilibrium points extracted at the end of each holding phase. At the same time, Fig. 6b suggests that the equilibrium relation is not affected by the stretch rate. This observation is supported by the statistical analysis: the MWU test performed at each equilibrium point did not reveal significant differences between the high and low stretch rates. Examples of this statistical comparison are reported in Fig. 6c for the equilibrium stresses attained at the end of the first and fifth loading steps.

**Figure 6.**
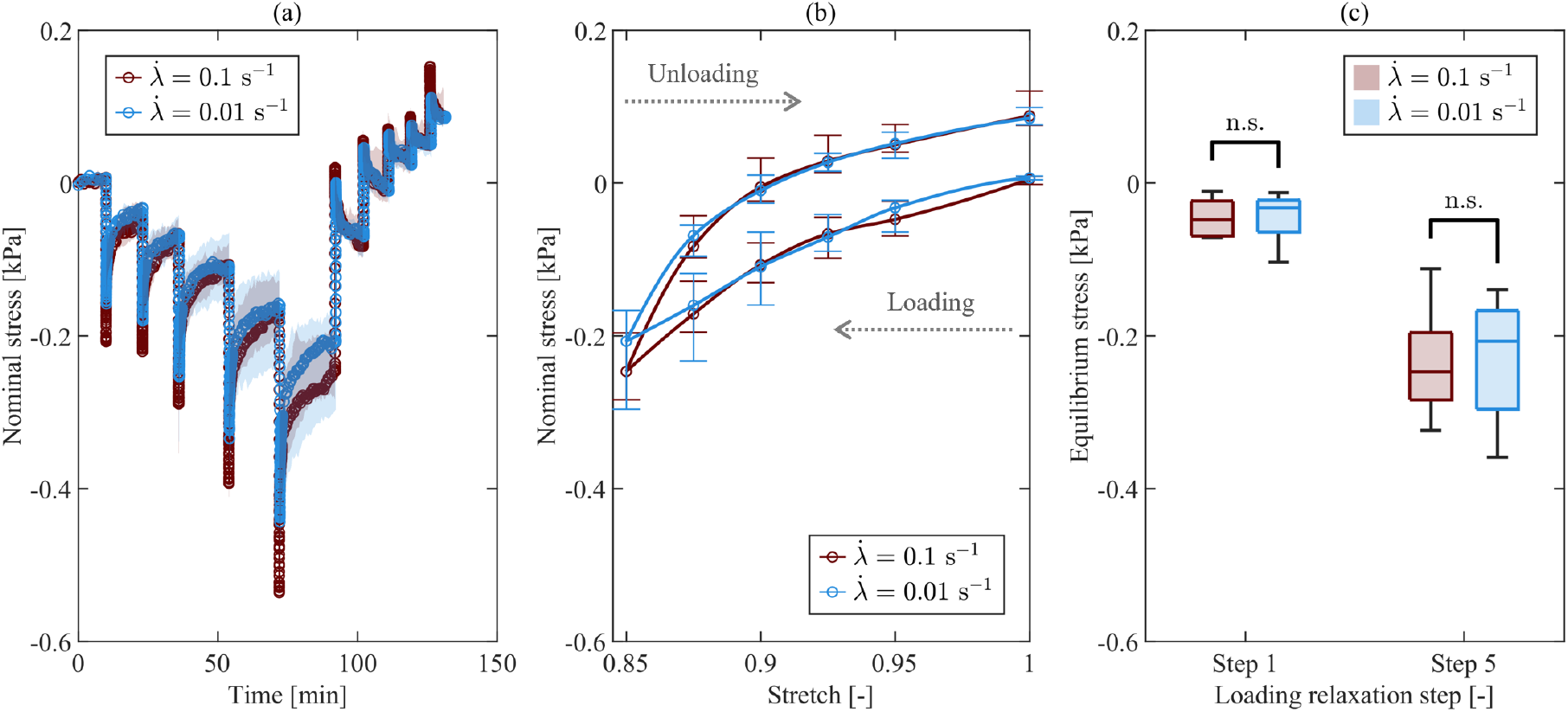
Brain tissue equilibrium response in compression at high (red) and low (blue) stretch rates. (a) Median nominal stress–time with IQR (shaded areas); (b) equilibrium stress–stretch relation with IQR (bars); (c) comparison of equilibrium stress at two representative loading steps (Step 1: *λ* = 0.95; Step 5: *λ* = 0.85) between stretch rates, showing no significant differences.

The inelastic response of brain tissue in compression is analyzed in Fig. 7. Figure 7a shows the median nominal stress as a function of stretch at high and low stretch rates (shown in red and blue, respectively), highlighting the highly hysteretic and dissipative behavior during loading and unloading. The gray, orange, and green areas provide a graphical representation of the inelastic quantitative metrics introduced in Sect. 2.3.2, respectively: the stress work associated with the equilibrium response, the accumulated dissipation during loading, and during unloading. This visualization allows the contributions to the overall dissipative response to be qualitatively compared between stretch rates. Their quantitative assessment is reported in Fig. 7b. According to the MWU test, the accumulated dissipation during loading 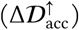 and unloading 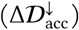 was significantly affected by stretch rate (p = 0.0140 and p = 0.0036, respectively). In contrast, no statistically significant evidence was found with respect to the equilibrium stress work (∫ Δ*P*^∞^d*λ*).

**Figure 7.**
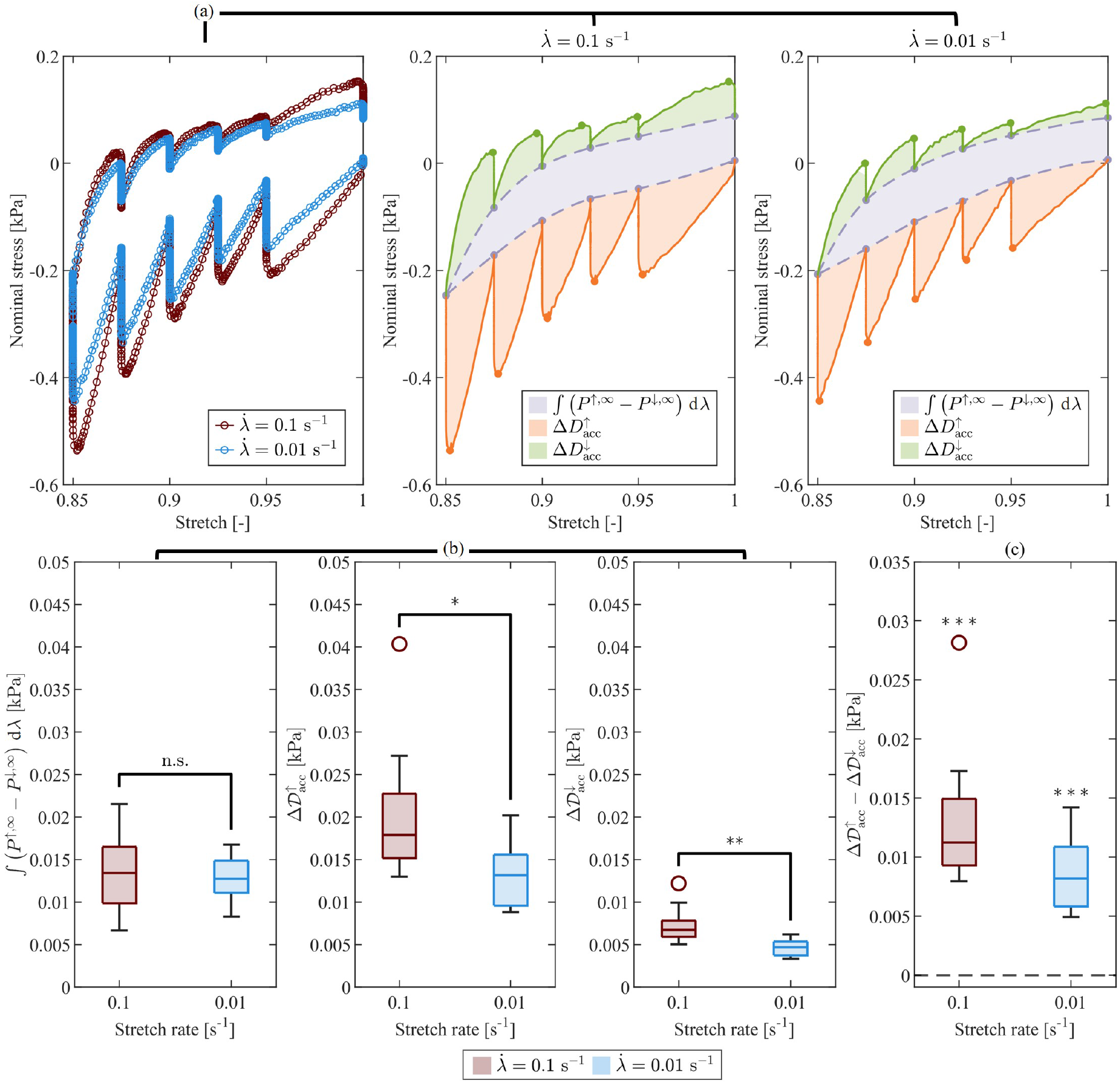
Brain tissue inelastic effects in compression at high (red) and low (blue) stretch rates. (a) Median nominal stress–stretch response with shaded areas representing the stress work associated with the equilibrium relation (gray), and the accumulated dissipation during loading (orange) and unloading (green), at the two stretch rates. The gray dots indicate the equilibrium stresses attained at the end of each relaxation step, while the orange and green dots show the peak stresses reached at the end of each loading and unloading ramp, respectively; (b) comparison of the energy-based metrics between stretch rates (MWU test), showing rate-dependent accumulated dissipation (*p<0.05; **p<0.01) but no evidence of significant rate dependence in the equilibrium stress work; (c) comparison between loading and unloading accumulated dissipation (one-tailed WSR test), indicating significantly higher dissipation during loading at both stretch rates (***p<0.001).

A one-tailed WSR test further indicates that the difference in accumulated dissipation between loading and unloading, 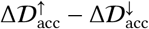, was significantly greater than zero at both stretch rates (p = 0.9766·10^−3^ in both cases, Fig. 7c). This result suggests that stress relaxation is large during loading, which can also be qualitatively appreciated from the stress–stretch curves in Fig. 7a at both stretch rates.

Further insights into the relaxation of brain tissue in compression is provided by Fig. 8, which analyzes the normalized nominal stress evolution during the holding phases (see Sect. 2.3.3 for its definition). Figure 8a reports the normalized nominal stress during the holding phases associated with all loading steps (Steps 1–5) at the high and low stretch rates. At both rates, a dependence of the normalized stress on the stretch level is observed. In particular, lower normalized stress values were observed at lower deformation levels, indicating a greater relative stress relaxation.

**Figure 8.**
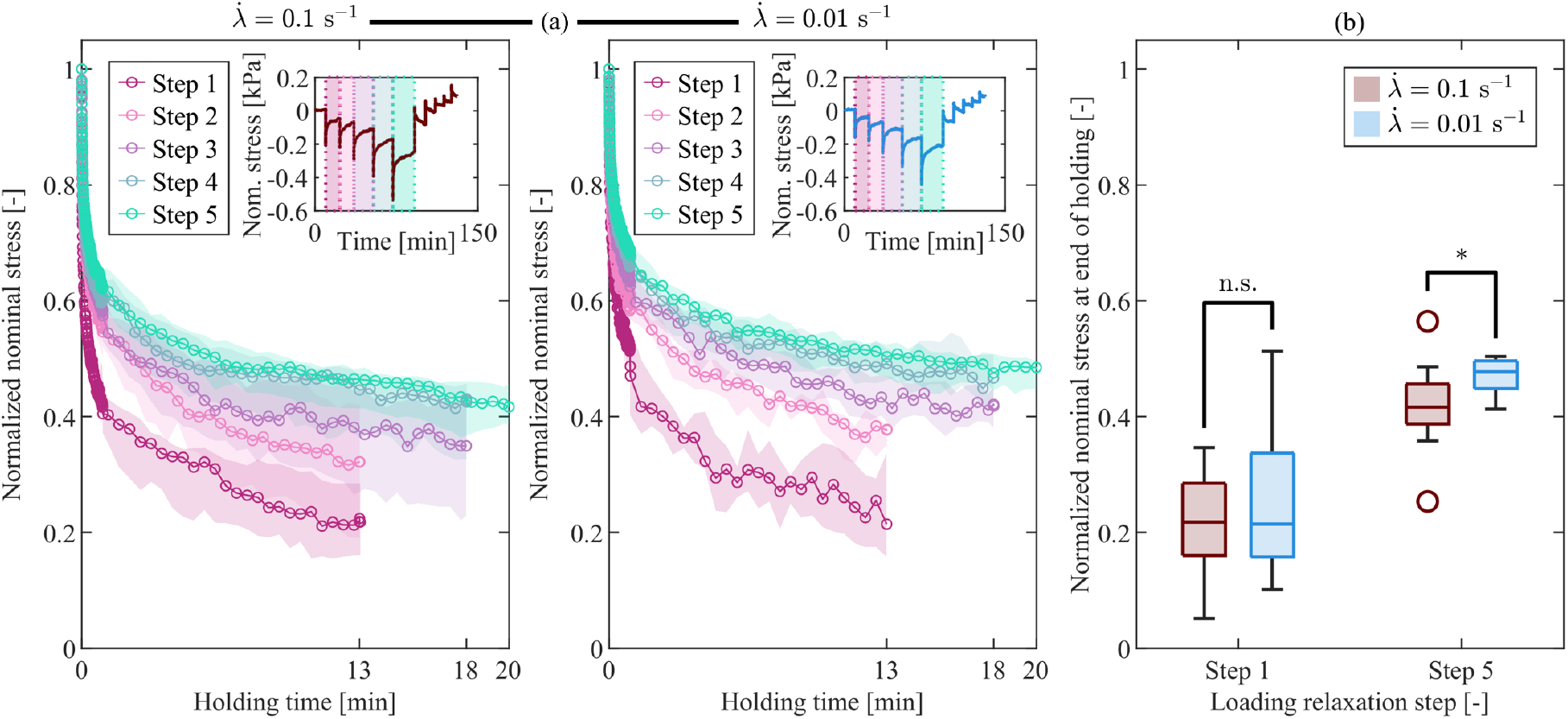
Brain tissue normalized stress relaxation response in compression at high and low stretch rates. (a) Median normalized nominal stress during the holding phases for all loading steps and both stretch rates. Shaded regions indicate the interquartile range (IQR). The insets show the corresponding median nominal stress–time curves and identify the individual loading steps; (b) comparison of normalized stress between the high (red) and low (blue) stretch rates for two representative loading steps (Step 1: *λ* = 0.95; Step 5: *λ* = 0.85). No significant difference was observed for Step 1, whereas a significant difference was found for Step 5 (*p<0.05).

The MWU test performed on the normalized stress at the end of the holding phases did not identify significant differences between stretch rates at the lower deformation levels (Steps 1–3). In contrast, significantly lower normalized stress values were observed at the high stretch rate for Step 4 (*p* = 0.0376) and Step 5 (*p* = 0.0452). Representative results for Steps 1 and 5 are reported in Fig. 8b. These findings indicate that the effect of stretch rate on relaxation becomes relevant only at higher deformation.

#### 3.1.2. Comparison with simulant materials

Porcine brain tissue and simulant materials were compared to assess the mechanical fidelity of each material in compression. The individual responses of the simulant materials, analyzed analogously to brain tissue, are reported in Sect. S2 of the Supplementary Material available online. That section also presents the comparison across all materials at the low stretch rate, whereas those at the high stretch rate are presented below.

Comparison of the equilibrium response in compression across materials revealed significant differences in stress magnitude at both stretch rates. Figure 9a illustrates the nominal stress–time responses at the higher stretch rate. Ecoflex exhibited markedly higher stresses than brain tissue, CH, and Sylgard at equivalent stretch levels. Therefore, the equilibrium stress–stretch relation is complemented by an inset focused on the stress range attained by brain tissue, CH, and Sylgard (Fig. 9b), allowing a clearer comparison among these softer materials. For the equilibrium stresses at the representative loading Steps 1 and 5, Fig. 9c show statistically significant differences between brain tissue and each simulant material, with p-values below 0.001 according to the MWU test. The equilibrium response of CH and Sylgard did not exhibit rate-dependence at any deformation level (MWU test, p > 0.05; Supplementary Material, Figs. S3–S4). This behavior is consistent with that observed for brain tissue. In contrast, Ecoflex exhibited a statistically significant dependence on stretch rate during loading at the higher deformation levels, corresponding to Steps 3–5 (Supplementary Material, Fig. S5).

**Figure 9.**
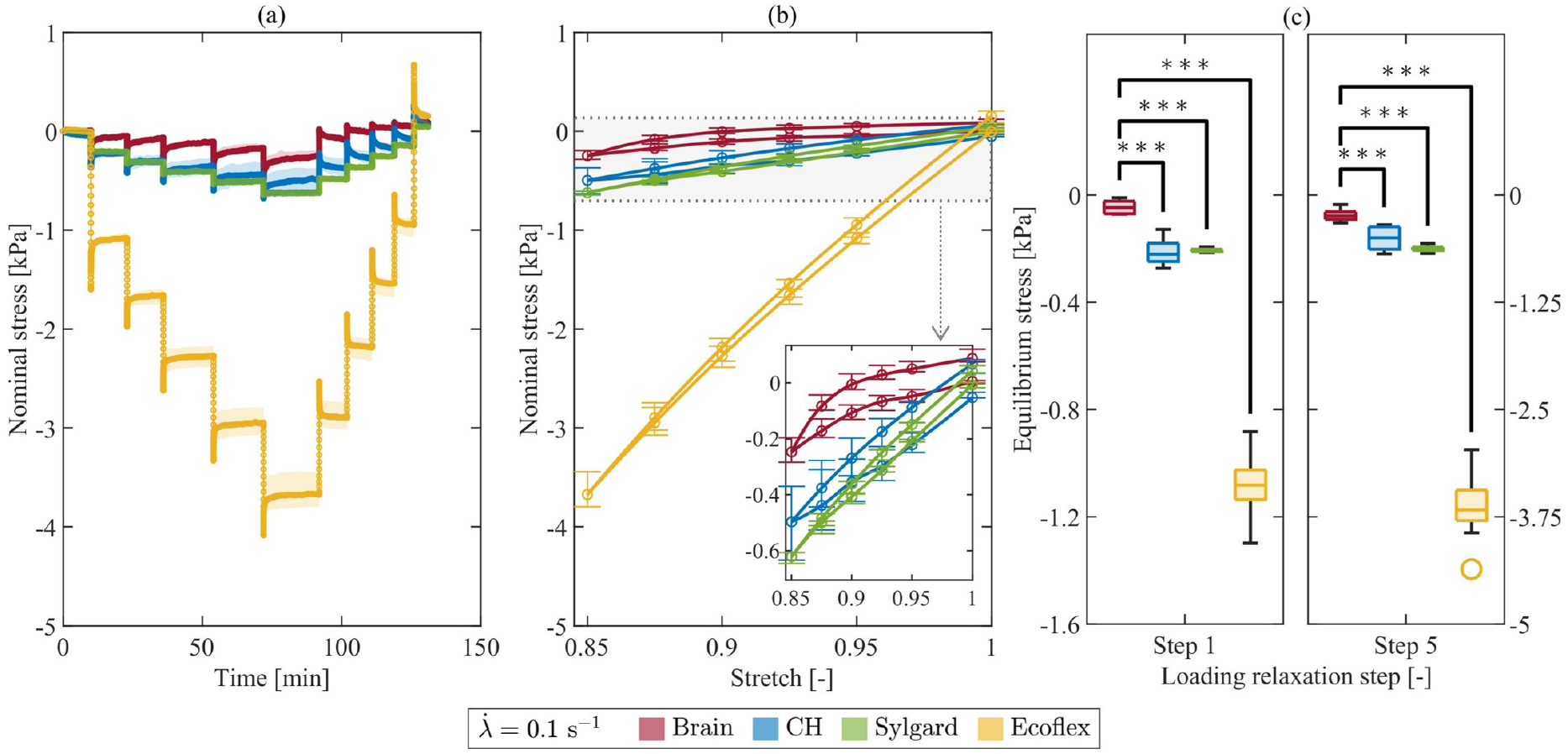
Comparison of the equilibrium response in compression across all materials at the high stretch rate. (a) Median nominal stress–time with IQR (shaded areas); (b) equilibrium stress–stretch relation with IQR (bars). The inset enlarges the stress range relevant to brain tissue, CH, and Sylgard to facilitate comparison; (c) comparison of equilibrium stress at two representative loading steps (Step 1: *λ* = 0.95; Step 5: *λ* = 0.85) between brain tissue and each simulant material (***p<0.001). Corresponding results at the low stretch rate are provided in Fig. S6 of the Supplementary Material.

The comparison was subsequently extended to the inelastic response. As observed for brain tissue, the equilibrium stress work did not exhibit statistically significant rate dependence for any of the simulant materials (p > 0.05; Supplementary Material, Figs. S7b–S9b). In contrast, the accumulated dissipation during both loading and unloading was significantly greater at the high stretch rate for all materials (CH: p = 0.0073 in loading, p = 0.0173 in unloading; Sylgard: p = 0.0079 in loading and unloading; Ecoflex: p = 1.8267 · 10^−4^ in loading and unloading). This trend is consistent with the rate-dependent dissipative response observed in brain tissue.

Pairwise comparisons between each simulant material and brain tissue at the high stretch rate are illustrated in Fig. 10. Figure 10a shows the nominal stress–stretch curves for all materials. To improve visual clarity, the areas enclosed by the loading and unloading curves are not highlighted. The corresponding quantitative comparisons based on the energy-related metrics are reported in Fig. 10b.

**Figure 10.**
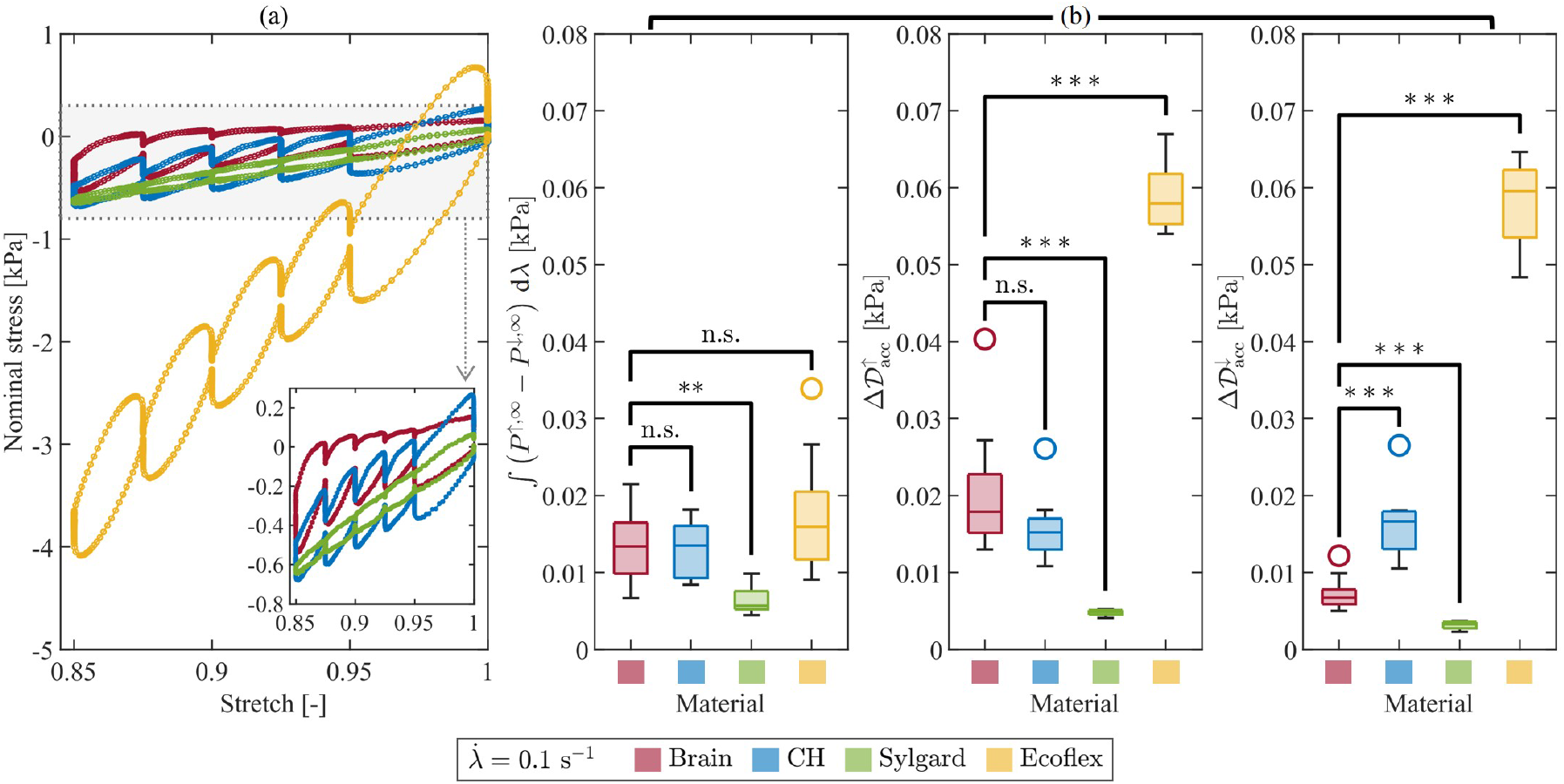
Comparison of inelastic effects in compression across all materials at the high stretch rate. (a) Median nominal stress–stretch response. The inset enlarges the stress range relevant to brain tissue, CH, and Sylgard to facilitate comparison; (b) comparison of the energy-based metrics between brain tissue and each simulant material (**p<0.01; ***p<0.001). Corresponding results at the low stretch rate are provided in Fig. S10 of the Supplementary Material.

Among the simulant materials, CH exhibited the closest overall agreement with brain tissue. In particular, no statistically significant differences with brain tissue were observed for the equilibrium stress work and the accumulated dissipation during loading. However, the dissipation during unloading was significantly greater for CH than for brain tissue (high rate: p = 0.2461 · 10^−3^; low rate: p = 0.1827 · 10^−3^; Supplementary Material, Fig. S10b). This results suggest that CH does not reproduce the asymmetric inelastic behavior observed in brain tissue. Consistently, the one-tailed WSR test did not provide evidence that dissipation during loading was significantly greater than that during unloading at either stretch rate (Supplementary Material, Fig. S7c). This is in disagreement with brain tissue (Fig. 7c).

Sylgard exhibited significantly lower equilibrium stress work than brain tissue at both stretch rates (high rate: p = 0.0080; low rate: p = 0.0027). It also exhibited lower accumulated dissipation during both loading and unloading at both stretch rates (p = 0.6660 · 10^−3^). This systematic reduction in the energy-based metrics suggests that Sylgard under-represents the inelastic contribution to the compressive mechanical response of brain tissue. Nevertheless, Sylgard reproduced the loading–unloading asymmetry in dissipation observed for brain tissue. Specifically, the WSR test indicated that dissipation was significantly greater during loading than during unloading at both stretch rates (p = 0.0313; Supplementary Material, Fig. S8c).

Ecoflex generally exhibited energy-based metrics that differed significantly from those of brain tissue, indicating a tendency to over-represent the energetic contributions associated with the mechanical response. The accumulated dissipation during both loading and unloading was greater than that of brain tissue at both stretch rates (p = 0.1827 · 10^−3^). A similar trend was observed for the equilibrium stress work at the low stretch rate (p = 0.4586 · 10^−3^; Supplementary Material, Fig. S10b); however, at the high stretch rate, no statistically significant difference in equilibrium stress work was identified. The one-tailed WSR test confirmed significantly greater dissipation during loading than during unloading, as observed for brain tissue, only at the low stretch rate (p = 0.0020; Supplementary Material, Fig. S9c).

The comparison was extended to the normalized nominal stress response. Figure 11a shows the normalized stress relaxation in compression during the holding phase at the high rate across all materials at two representative loading steps (Step 1: *λ* = 0.95; Step 5: *λ* = 0.85).

**Figure 11.**
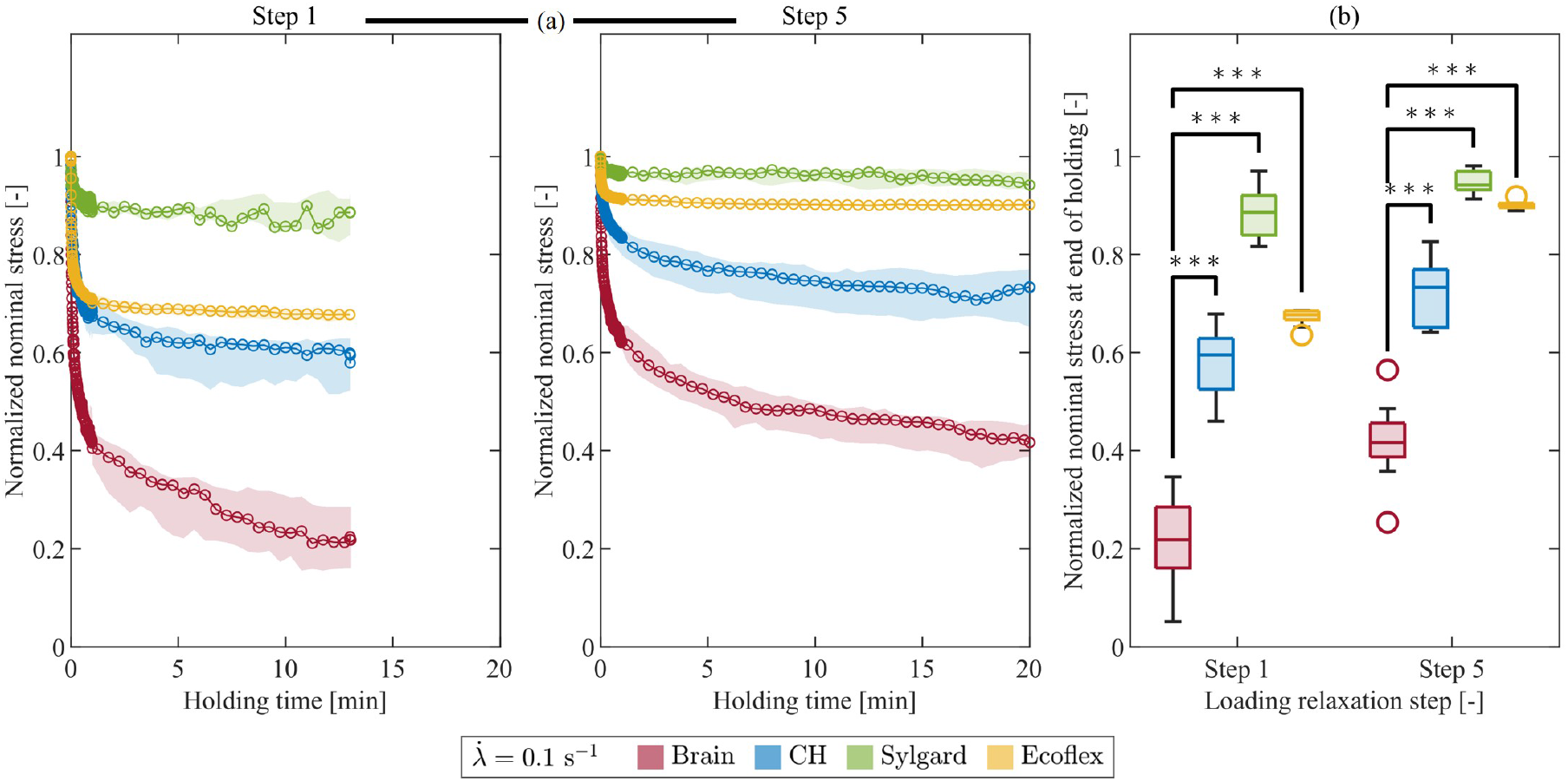
Comparison of normalized stress relaxation in compression across all materials at the high stretch rate for two representative loading steps (Step 1: *λ* = 0.95; Step 5: *λ* = 0.85). (a) Median normalized nominal stress during the holding phase with IQR (shaded areas); (b) comparison of normalized nominal stress at the end of the holding phase between brain tissue and each simulant material (***p<0.001). Corresponding results at the low stretch rate are provided in Fig. S14 of the Supplementary Material.

The analysis of stress decay suggests distinct relaxation behaviors among the simulant materials and brain tissue. To quantify these differences, the normalized nominal stress was evaluated at six time points during the holding phases of the selected loading steps, as explained in Sect. 2.3.3. Brain tissue was characterized by rapid stress decay, particularly within the first minute of the holding phase. At the high stretch rate, normalized stress decreased from 0.81 to 0.22 during Step 1, and from 0.89 to 0.42 during Step 5. A similar trend was observed at the low stretch rate, with normalized stress values decreasing from 0.90 to 0.21 (Step 1) and from 0.93 to 0.48 (Step 5) (Fig. S14, Supplementary Material).

In contrast, all synthetic materials retained a greater fraction of their peak stress throughout the holding phase. Sylgard exhibited the most limited relaxation, with normalized stress remaining above 0.89 under all tested conditions. CH showed more pronounced relaxation than Sylgard, reaching a normalized stress of 0.59 in the condition associated with the greatest relaxation (Step 1 at high stretch rate). Nevertheless, its relaxation remains significantly slower than that of brain tissue. Ecoflex displayed an intermediate response at the lower deformation level (Step 1), with normalized stress of approximately 0.68 and 0.79 for the high and low stretch rates, respectively. On the other hand, at the higher deformation level (Step 5), its response approached that of Sylgard, with normalized stress remaining above 0.90 at both stretch rates.

Statistical analysis confirmed that all investigated simulant materials exhibit slower relaxation behavior compared to brain tissue. Statistically significant differences are observed at all selected time points, from the early stages of relaxation (1–10 s) to the end of the holding phase, for both stretch rates and both loading steps (p < 0.001). The comparisons at the end of the holding phase are shown in Fig. 11b. A detailed quantitative comparison of normalized stress values at all selected time points is provided in the Supplementary Material (Tables S1 and S2 for the high and low stretch rates, respectively).

Finally, the simulant materials exhibited rate-dependence trends in the normalized stress relaxation that differed from those of brain tissue (Supplementary Material, Figs. S11–S13). While statistically significant differences between stretch rates were observed in the brain only at higher deformation levels (Fig. 8b), CH and Sylgard showed a statistically significant dependence on the stretch rate only at the lowest deformation level (Step 1), whereas Ecoflex displayed a pronounced rate dependence at all deformation levels. Further details are provided in the Supplementary Material.

#### 3.1.3. Summary

In general, the main findings from the compression experiments can be summarized as follows:

- *Stress magnitude and equilibrium response*. None of the simulant materials fully reproduced the equilibrium stress response of brain tissue at the stretch levels considered. However, CH and Sylgard provided a closer approximation than Ecoflex, exhibiting more comparable stress magnitudes throughout the protocol and no statistically significant rate dependence of the equilibrium stress–stretch curve, consistent with brain tissue.
- *Inelastic effects*. None of the simulant materials fully reproduced the inelastic response of brain tissue,. Nevertheless, CH showed the closest overall agreement in the magnitude of the energy-based metrics, whereas Sylgard and Ecoflex displayed larger discrepancies. All materials reproduced the qualitative rate dependence observed in brain tissue, namely greater dissipation at the high stretch rate. However, only Sylgard reproduced the loading-unloading asymmetry in dissipation observed in brain tissue, with greater dissipation during loading than during unloading.
- *Normalized stress relaxation*. All simulant materials exhibited significantly slower normalized stress relaxation than brain tissue, although CH provided the closest approximation. None of the materials reproduced the rate-dependence pattern observed in brain tissue, for which statistically significant differences in normalized stress at the end of the holding phase emerged between stretch rates only at the higher deformation levels.

### 3.2. Tension

#### 3.2.1. Brain tissue response

Figure 12a shows the median nominal stress as a function of time, together with the corresponding IQR, at high and low stretch rates (shown in red and blue, respectively). In contrast to the compression response, a reversal in the sign of the nominal stress during unloading occurs only near the end of the protocol, as the specimen returns towards the undeformed configuration. The equilibrium stress values attained at identical stretch levels during loading and unloading do not fully overlap, indicating the presence of equilibrium hysteresis also in tension (Fig. 12b), albeit smaller than in compression. More specifically, WSR tests revealed statistically significant differences between loading and unloading equilibrium stresses at the high stretch rate, except at the lowest deformation level (*λ* = 1.05). In contrast, significant differences at the low stretch rate emerged only at *λ* = 1.125. As in compression, the equilibrium stress–stretch relation did not exhibit significant rate dependence according to the MWU test. This is shown in Fig. 12c dor the representative equilibrium stresses obtained at the first and fifth loading steps.

**Figure 12.**
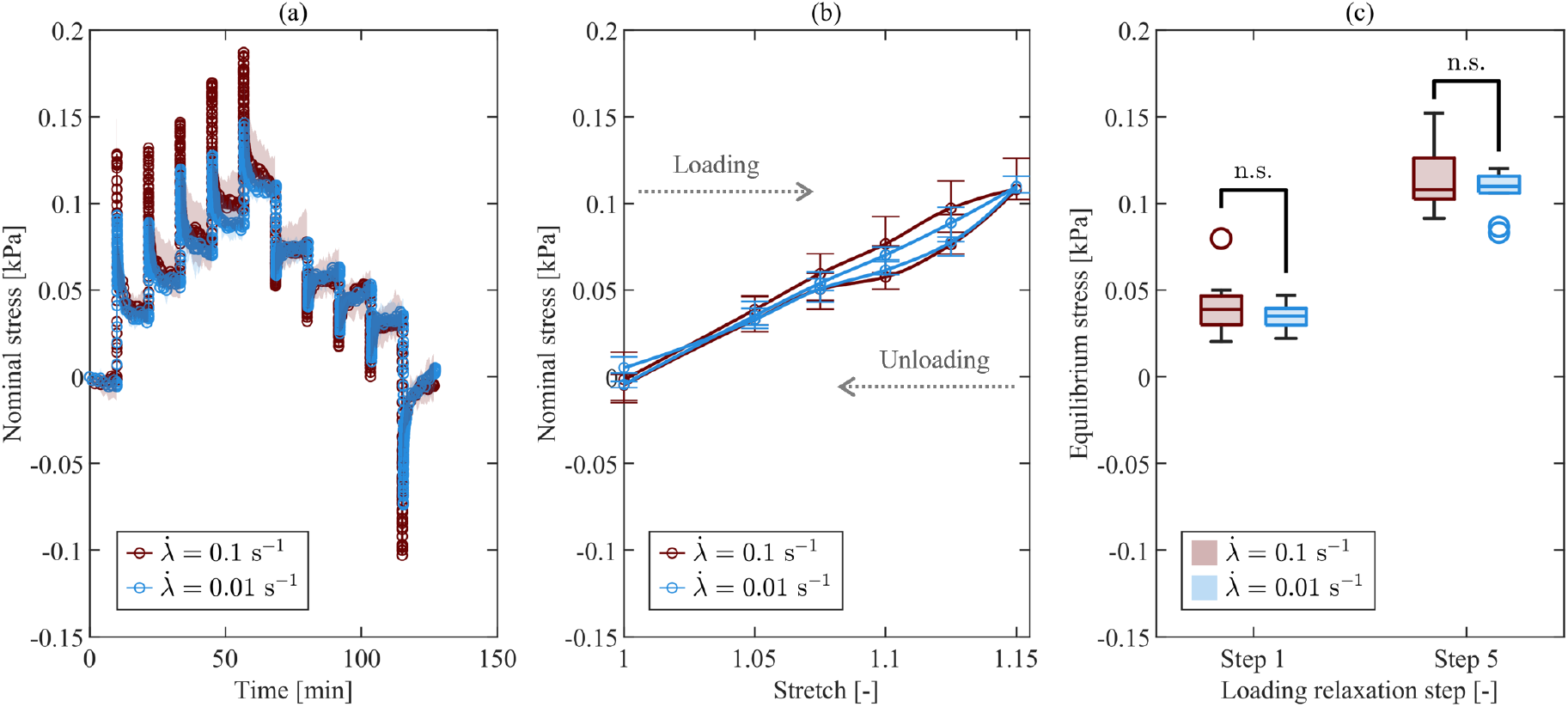
Brain tissue equilibrium response in tension at high (red) and low (blue) stretch rates. (a) Median nominal stress–time with IQR (shaded areas); (b) equilibrium stress–stretch relation with IQR (bars); (c) comparison of equilibrium stresses at two representative loading steps (Step 1: *λ* = 1.05; Step 5: *λ* = 1.15) between stretch rates, showing no significant differences.

The inelastic response of brain tissue in tension is analyzed in Fig. 13. Figure 13a shows the median nominal stress as a function of stretch at high and low stretch rates (shown in red and blue, respectively), highlighting the hysteretic and dissipative behavior during loading and unloading. The quantitative comparison of energy-based metrics between stretch rates is shown in Fig. 13b. According to the MWU test, the accumulated dissipation during loading 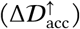 and unloading 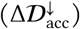 was significantly higher at the high stretch rate (p = 0.0013 and p = 1.8267 · 10^−4^, respectively). In contrast, no statistically significant rate dependence was found for the equilibrium stress work (∫ Δ*P*^∞^d*λ*). This behavior is consistent with that observed in compression.

**Figure 13.**
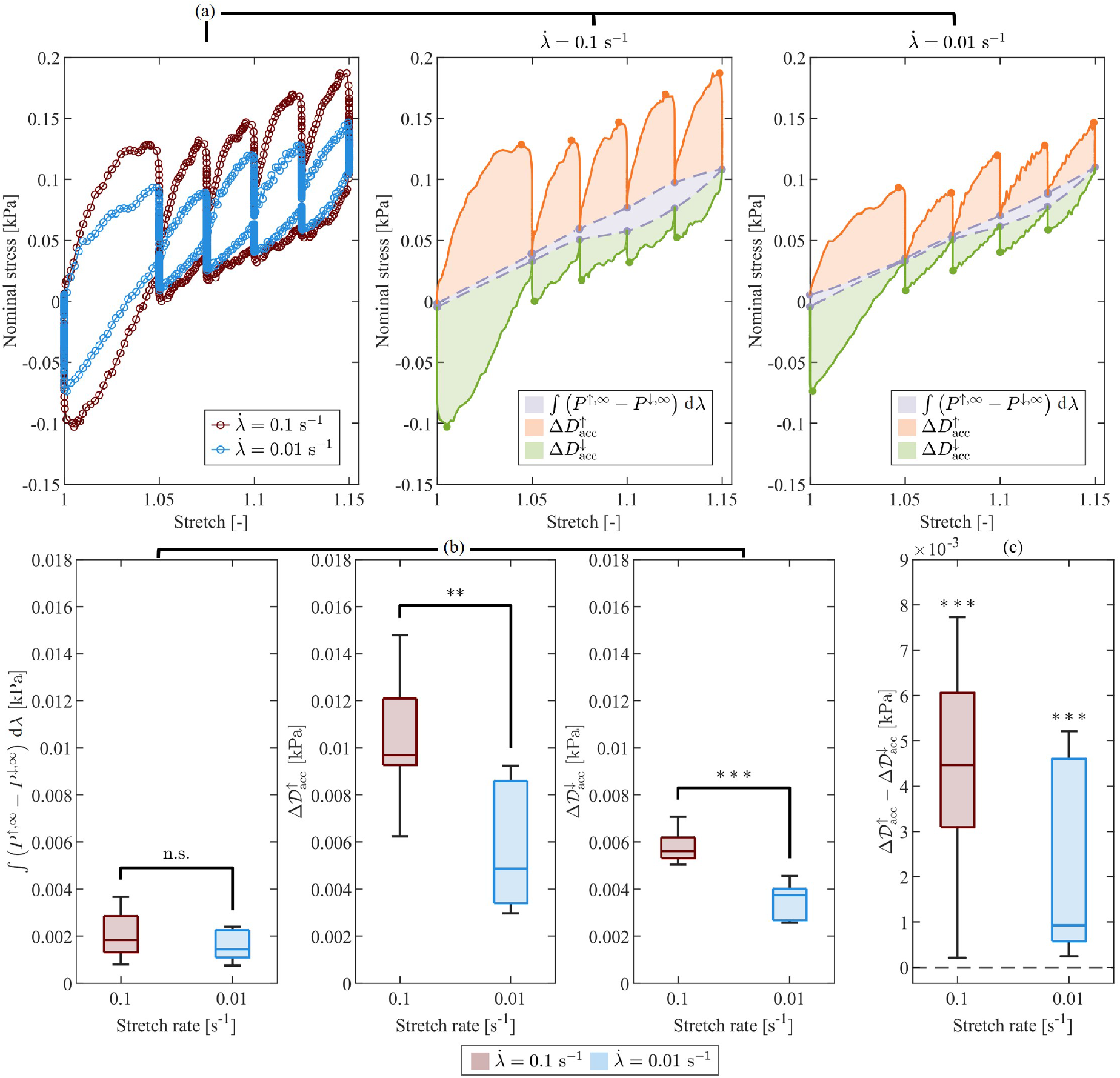
Brain tissue inelastic effects in tension at high (red) and low (blue) stretch rates. (a) Median nominal stress–stretch response with shaded areas representing the stress work associated with the equilibrium relation (gray), and the accumulated dissipation during loading (orange) and unloading (green), at the two stretch rates. The gray dots indicate the equilibrium stresses attained at the end of each relaxation step, while the orange and green dots show the peak stresses reached at the end of each loading and unloading ramp, respectively; (b) comparison of the energy-based metrics between stretch rates (MWU test), showing rate-dependent accumulated dissipation (**p<0.01; ***p<0.001) but no evidence of significant rate dependence in the equilibrium stress work; (c) comparison between loading and unloading accumulated dissipation (one-tailed WSR test), indicating significantly higher dissipation during loading at both stretch rates (***p<0.001).

A one-tailed WSR test further indicates that the accumulated dissipation during loading is significantly greater than during unloading at both high and low stretch rates (p = 9.7656 · 10^−4^; Fig. 13c). This finding is also consistent with the behavior previously observed in compression.

The relaxation behavior in tension was further analyzed in normalized form in Fig. 14. At both stretch rates, the initial loading steps exhibited lower normalized stress values throughout the holding phase (Fig. 14a), indicating a more pronounced stress relaxation at lower deformation levels. MWU tests comparing the normalized stress at the end of each holding phase between high and low stretch rates, identified statistically significant differences at all deformation levels during loading (Step 1: p = 0.0312; Step 2: p = 0.0091; Step 3: p = 0.0028; Step 4: p = 0.0046; Step 5: p = 7.6854 · 10^−4^). Representative comparisons for Steps 1 and 5 are shown in Fig. 14b. These findings indicate that the applied stretch rate affects the normalized stress relaxation response in tension across all deformation levels investigated. This differs from the compression response, for which significant rate-dependent difference emerged only at the higher stretch levels. Overall, the results suggest that the influence of stretch rate on normalized stress relaxation depends on the loading mode.

**Figure 14.**
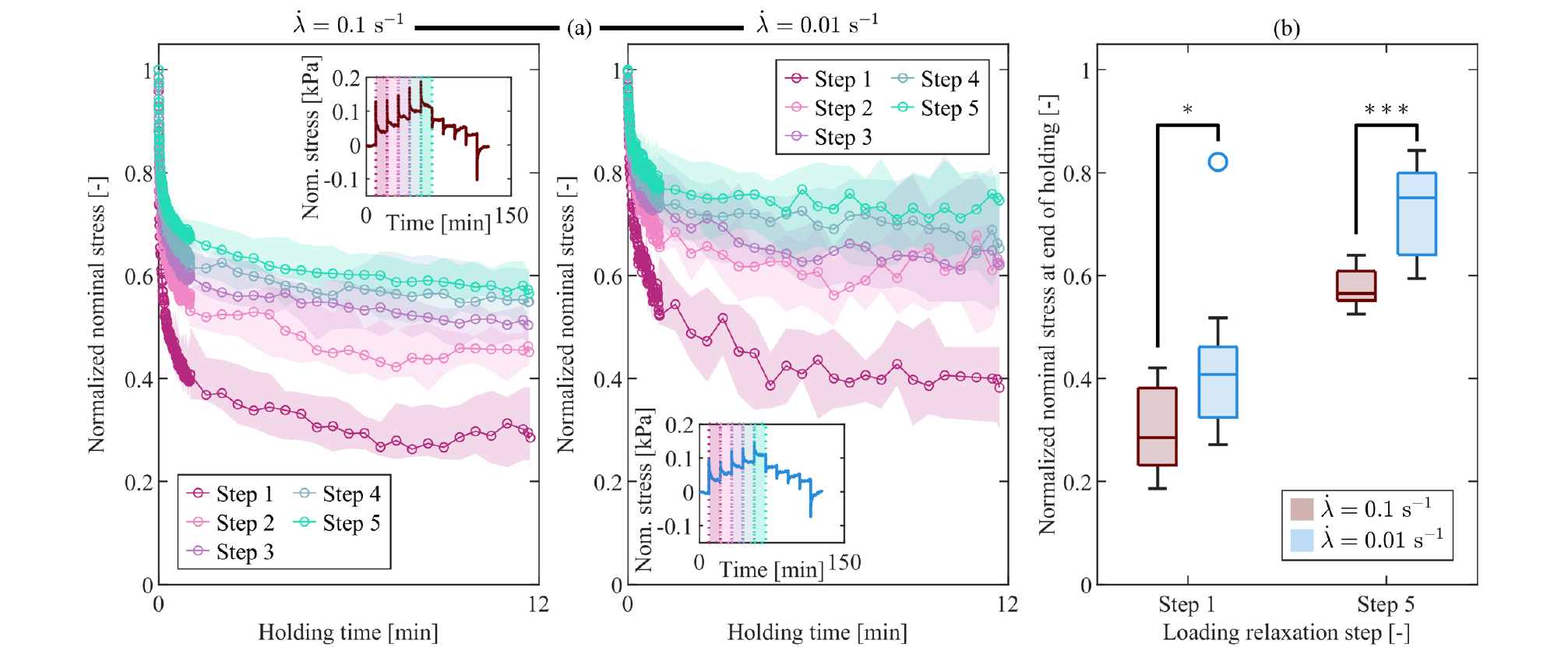
Brain tissue normalized stress relaxation response in tension at high and low stretch rates. (a) Median normalized nominal stress during the holding phases for all loading steps and both stretch rates. Shaded regions indicate the interquartile range (IQR). The insets show the corresponding median nominal stress–time curves and identify the individual loading steps; (b) comparison of normalized stress between the high (red) and low (blue) stretch rates for two representative loading steps (Step 1: *λ* = 1.05; Step 5: *λ* = 1.15). Significant differences were observed in both loading steps (*p<0.05; ***p<0.001).

#### 3.2.2. Comparison with simulant materials

Porcine brain tissue and simulant materials were compared to further assess their mechanical fidelity in tension. The individual responses of the simulant materials, analyzed analogously to brain tissue, are reported in Sect. S3 of the Supplementary Material available online. That section also presents the comparison across all materials at the low stretch rate, whereas those at the high stretch rate are presented below.

Similarly to the compression experiment, Ecoflex exhibited markedly higher stress values than brain tissue, CH, and Sylgard at equivalent stretch levels (Fig. 15a). At the selected loading Steps 1 and 5, statistically significant differences in equilibrium stress were observed between brain tissue and each simulant material (Fig. 15c). According to the MWU test, all comparisons are statistically significant at p < 0.001, except for CH at Step 1 at the high stretch rate, for which p < 0.01.

**Figure 15.**
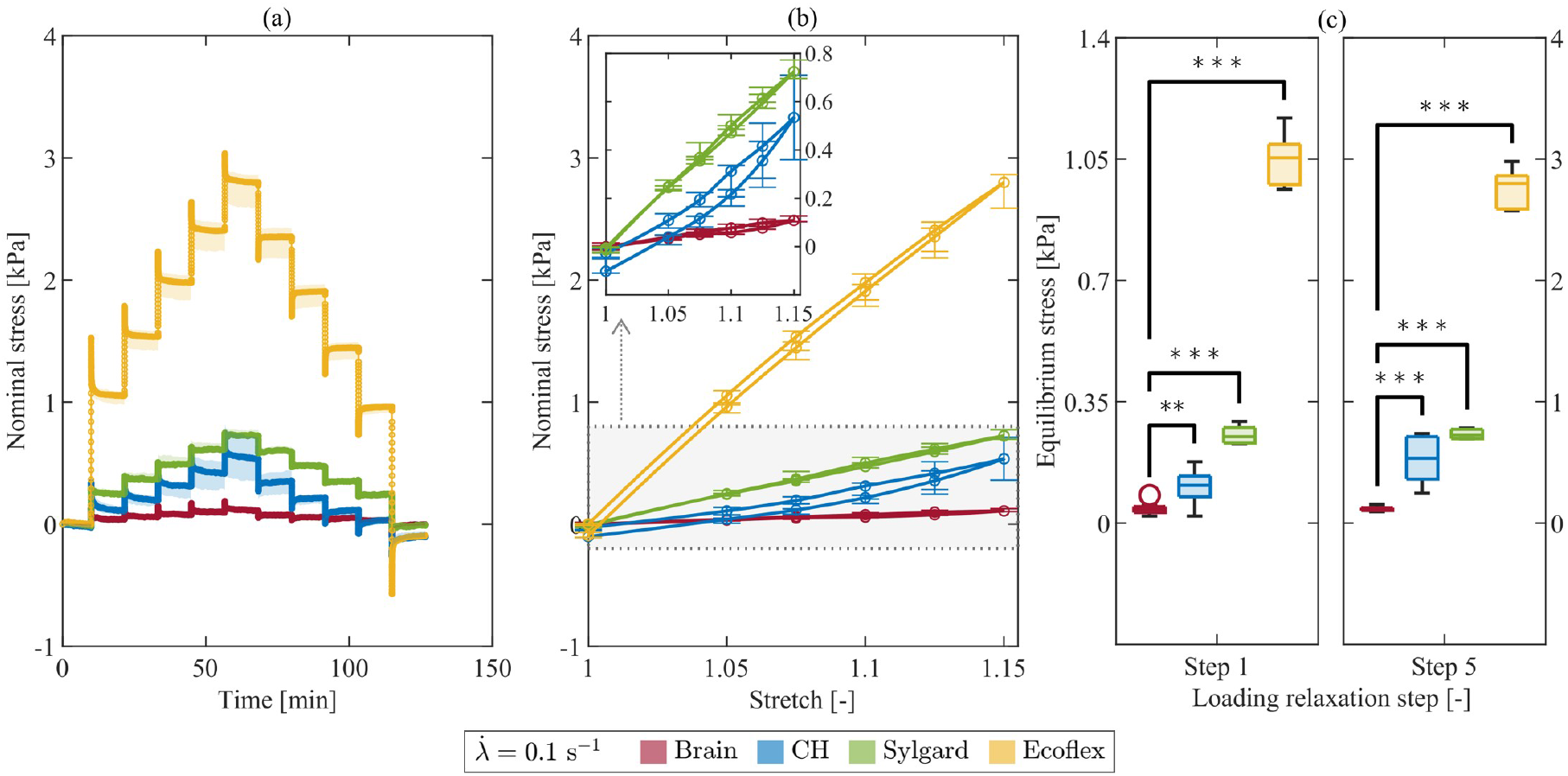
Comparison of equilibrium response in tension across all materials at the high stretch rate. (a) Median nominal stress–time with IQR (shaded areas); (b) equilibrium stress–stretch relation with IQR (bars). The inset enlarges the stress range relevant to brain tissue, CH, and Sylgard to facilitate comparison; (c) comparison of equilibrium stress at two representative loading steps (Step 1: *λ* = 1.05; Step 5: *λ* = 1.15) between brain tissue and each simulant material (**p<0.01; ***p<0.001). Corresponding results at the low stretch rate are provided in Fig. S18 of the Supplementary Material.

The equilibrium response did not exhibit statistically significant rate dependence for any of the simulant materials (MWU test, *p* > 0.05). This behavior is reported in the Supplementary Material (Figs. S15–S17), and is consistent with the response observed for brain tissue.

The inelastic behavior of the simulant materials was compared with that of brain tissue in tension. Figure 16a shows the nominal stress–stretch responses at the high stretch rate, whereas the corresponding energy-based metrics are quantitatively compared in Fig. 16b.

**Figure 16.**
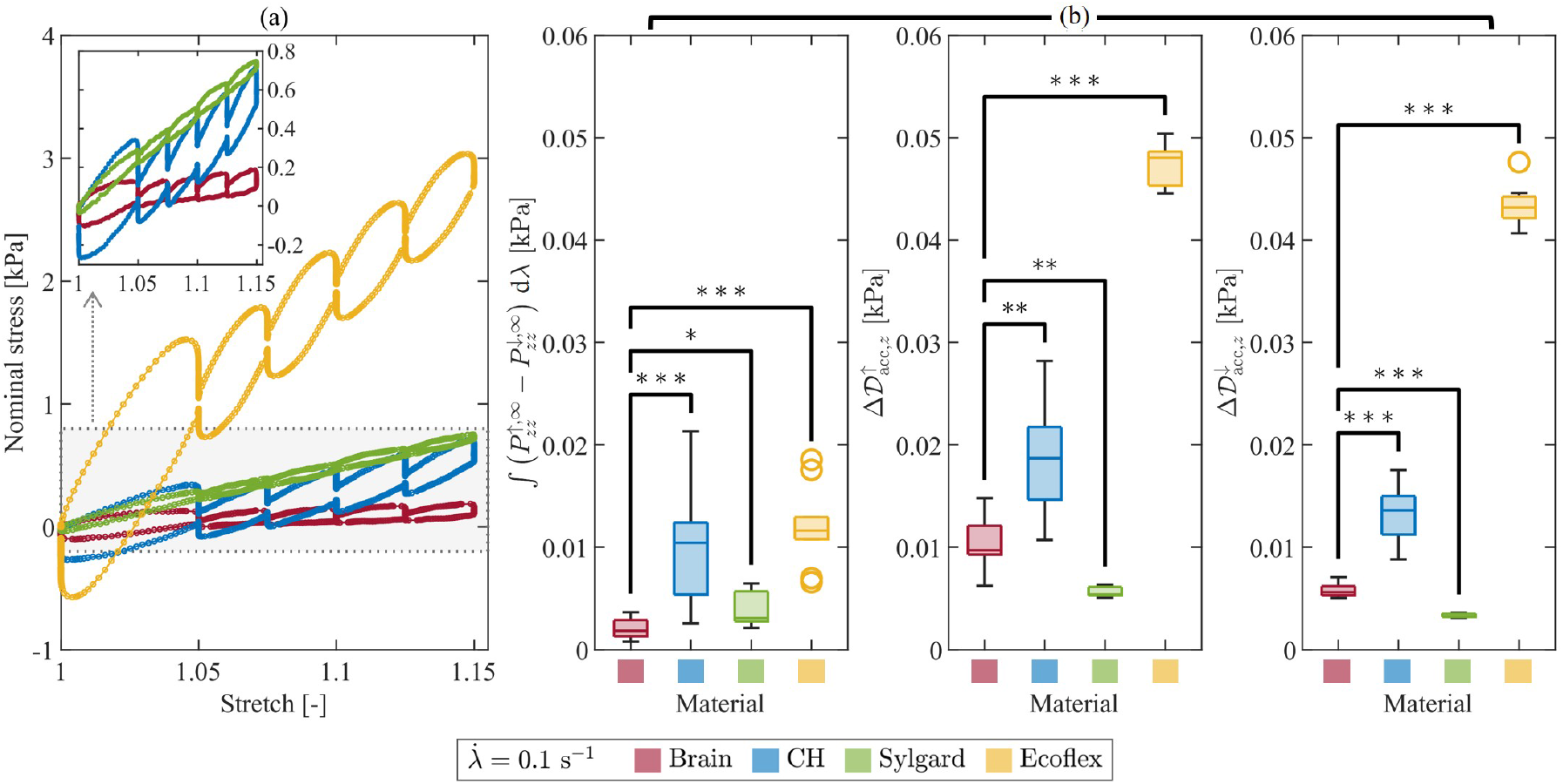
Comparison of inelastic effects in tension across all materials at the high stretch rate. (a) Median nominal stress–stretch response. The inset enlarges the stress range relevant to brain tissue, CH, and Sylgard to facilitate comparison; (b) comparison of the energy-based metrics between brain tissue and each simulant material (*p<0.05; **p<0.01; ***p<0.001). Corresponding results at the low stretch rate are provided in Fig. S22 of the Supplementary Material.

In contrast to its response in compression, CH exhibited greater values than brain tissue for all energy-based metrics in tension. Specifically, CH showed greater equilibrium stress work (high rate: p = 4.3964 × 10^−4^; low rate: p = 1.8267 × 10^−4^), accumulated dissipation during loading (high rate: p = 0.0017; low rate: p = 3.2984 × 10^−4^), and accumulated dissipation during unloading (both stretch rates: p = 1.8267 × 10^−4^). Furthermore, CH reproduced the asymmetric inelastic behavior observed in brain tissue in tension, with significantly greater accumulated dissipation during loading than unloading at both stretch rates (WSR test, p = 9.7656 · 10^−4^; Supplementary Material, Fig. S19c).

Sylgard showed closer correspondence with brain tissue in tension than in compression. At the low stretch rate, no statistically significant differences from brain tissue were observed for equilibrium stress work or accumulated dissipation during loading, whereas the accumulated dissipation during unloading was significantly lower (*p* = 6.6600 · 10^−4^; Supplementary Material, Fig. S22b). At the high stretch rate, Sylgard exhibited significantly greater equilibrium stress work than brain tissue (p=0.0400), while accumulated dissipation during loading and unloading is significantly lower (p=0.0013 and p=6.6600 · 10^−4^, respectively). These results indicate that the agreement between Sylgard and brain tissue depends on the stretch rate. Nevertheless, similar to brain tissue, Sylgard exhibited greater accumulated dissipation during loading than during unloading at both stretch rates (Supplementary Material, Fig. S20c; WSR test, p = 0.0313).

Consistent with the compression results, Ecoflex showed greater values than brain tissue for the energy-based metrics in tension. Statistically significant differences were observed for equilibrium stress work and accumulated dissipation during both loading and unloading at both stretch rates (p = 1.8267 · 10^−4^). Unlike in compression, Ecoflex exhibited significantly greater accumulated dissipation during loading than during unloading at both stretch rates (Supplementary Material, Fig. S21c; WSR test, p = 9.7565 · 10^−4^).

Finally, no statistically significant rate dependence in the equilibrium stress work was observed for any of the simulant materials (Supplementary Material, Figs. S19b–S21b). In contrast, the accumulated dissipation during both loading and unloading was significantly greater at the high stretch rate for Sylgard (p = 0.0079) and Ecoflex (p = 1.8267 · 10^−4^). CH did not show statistically significant rate dependence for any energy-based metric. Therefore, among the simulant materials, only Sylgard and Ecoflex reproduced the rate dependence of the inelastic response observed for brain tissue in tension.

A final comparison was performed for normalized stress relaxation in tension. Figure 17a shows the normalized nominal stress during the holding phase for brain tissue and the simulant materials at two representative loading steps (Step 1: *λ* = 1.05; Step 5: *λ* = 1.15), at the high stretch rate.

**Figure 17.**
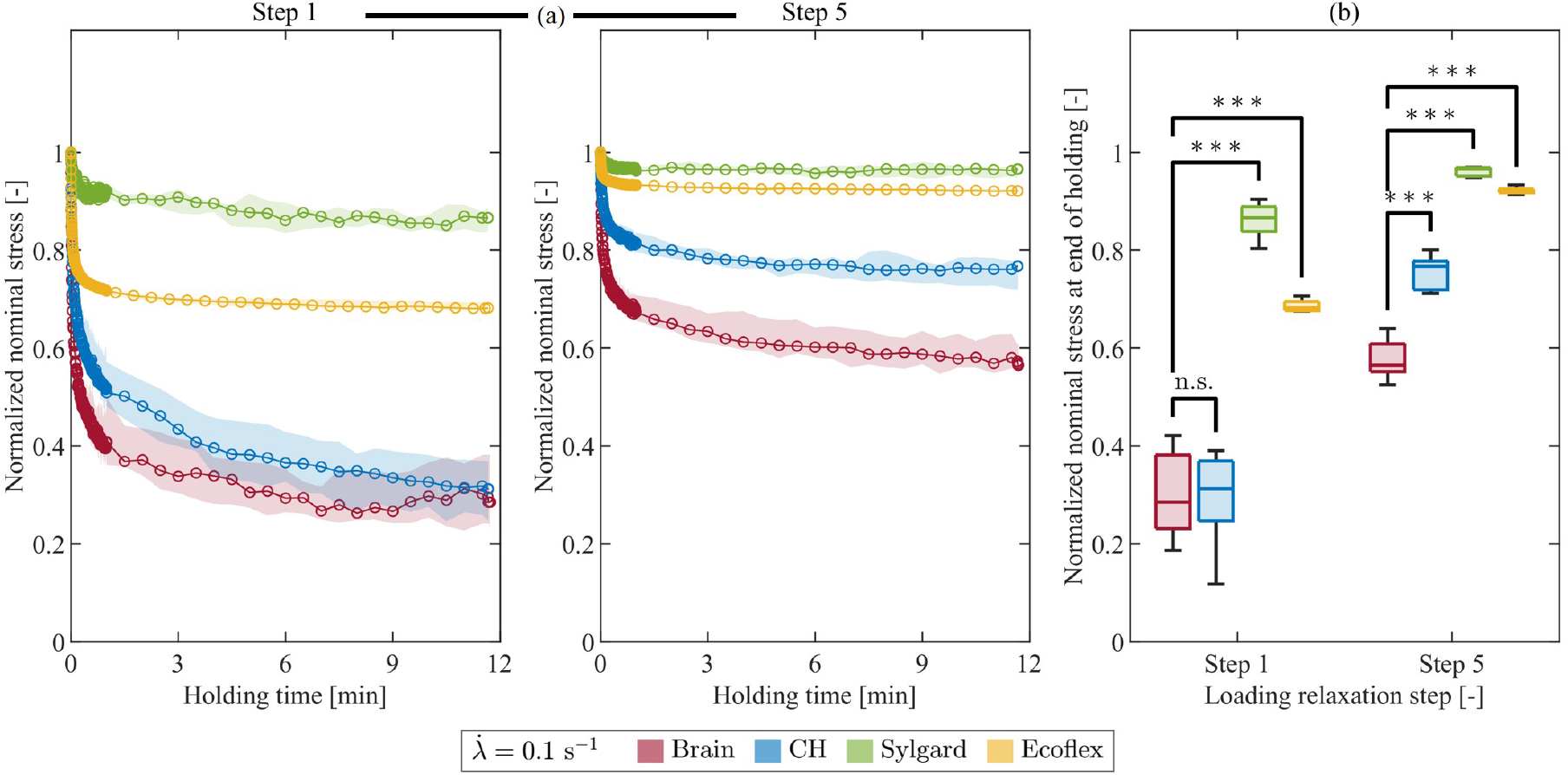
Comparison of normalized stress relaxation in tension across all materials at the high stretch rate for two representative loading steps (Step 1: *λ* = 1.05; Step 5: *λ* = 1.15). (a) Median normalized nominal stress during the holding phase with IQR (shaded areas); (b) comparison of normalized nominal stress at the end of the holding phase between brain tissue and each simulant material (***p<0.001). Corresponding results at the low stretch rate are provided in Fig. S26 of the Supplementary Material.

Similarly to the compression behavior, brain tissue exhibited rapid stress decay after the peak, with normalized stress decreasing to approximately approximately 40% of its peak value within the first minute of relaxation. At the high stretch rate, the normalized stress decreases from 0.77 to 0.29 during Step 1 and from 0.88 to 0.57 during Step 5. A comparable trend is observed at the low stretch rate, with normalized stress decreasing from 0.90 to 0.38 during Step 1 and from 0.93 to 0.73 during Step 5 (Fig. 17, Supplementary Material). Among the simulant materials, Sylgard and Ecoflex exhibited relaxation curves that remained consistently distinct from those of brain tissue. Specifically, Sylgard exhibited the most limited relaxation, with normalized stress remaining above 0.87 under all investigated conditions. Ecoflex showed a similar tendency, particularly at the higher deformation level, where the normalized stress remained above 0.92 at the high stretch rate and above 0.94 at the low stretch rate. At the lower deformation level, Ecoflex relaxed more substantially, reaching normalized stress values of 0.68 and 0.80 at the end of the holding phase for the high and low stretch rates, respectively. However, these value remained consistently greater than those measured for brain tissue.

In contrast, CH showed remarkably closer agreement with brain tissue in tension over most of the holding phases. At the high stretch rate, its normalized stress decreased from 0.87 to 0.31 during Step 1, approaching the brain tissue response at the end of the holding phase. At the higher deformation level, Step 5, CH exhibited more limited relaxation, with normalized stress decreasing from 0.95 to 0.77. At the low stretch rate, CH closely followed the brain-tissue response at both deformation levels, decreasing from 0.93 to 0.30 during Step 1 and from 0.96 to 0.76 during Step 5.

These findings are confirmed by the statistical analysis, illustrated for the high stretch rate at the end of the holding phase in Fig. 17b. A detailed quantitative comparison of normalized stress values at all selected time points is provided in the Supplementary Material (Tables S3 and S4 for the high and low stretch rates, respectively). At the high stretch rate, statistically significant differences from brain tissue were observed for Sylgard and Ecoflex at all investigated time points during both Step 1 and Step 5 (p < 0.001). CH provided a closer approximation to the normalized relaxation response of brain tissue than the other simulant materials. Nevertheless, statistically significant differences were found at nearly all investigated time points, with the only exception at the end of the holding phase during Step 1 (p = 0.9698). This indicates a convergence towards the normalized stress attained by brain tissue. At the low stretch rate, Sylgard and Ecoflex remained significantly different from brain tissue throughout the holding phase at both deformation levels. In contrast, CH showed no statistically significant differences from brain tissue at nearly all investigated time points during either Step 1 or Step 5 (p > 0.05). The only exception was observed at the end of the holding phase during Step 1 (p = 0.0312). Thus, CH closely reproduced the normalized relaxation response of brain tissue in tension at the low stretch rate.

Finally, the rate dependence of normalized stress relaxation in tension differed between brain tissue and the simulant materials (Supplementary Material, Figs. S23,–S25). Brain tissue revealed statistically significant differences between the high and low stretch rates at all investigated deformation levels. Ecoflex showed the same pattern of statistical significance. In contrast, CH and Sylgard did not exhibit statistically significant differences between stretch rates at any deformation level. Further details are provided in the Supplementary Material.

#### 3.2.3. Summary

In general, the main findings from the tension experiments can be summarized as follows:

- *Stress magnitude and equilibrium response*. None of the investigated simulant materials fully reproduced the equilibrium stress of brain tissue at the stretch levels considered. However, all materials correctly reproduced the absence of statistically significant rate dependence observed for brain tissue. CH and Sylgard provided a closer approximation than Ecoflex due to their stress levels remaining comparable with brain tissue across both loading and unloading.
- *Inelastic effects*. All simulant materials reproduced the loading–unloading asymmetry in accumulated dissipation observed in brain tissue, with greater dissipation during loading than during unloading. However, only Sylgard and Ecoflex captured the rate-dependent dissipative behavior, with Sylgard also partially approximating the magnitude of the energy-based metrics measured in brain tissue.
- *Normalized stress relaxation*. CH providesd the closest approximation of the normalized stress relaxation behavior of brain tissue, particularly at the low stretch rate. In contrast, Sylgard and Ecoflex retained a greater fraction of peak stress and exhibited slower relative stress decay than brain tissue. Conversely, Ecoflex was the only material to reproduce the consistent rate dependence of normalized stress relaxation observed for brain tissue across all investigated deformation levels.

## 4. Conclusions

Three synthetic simulant materials for brain phantoms were thoroughly compared to porcine brain tissue introducing a mechanical characterization based on multi-step stress relaxation protocols under uniaxial extension and compression. The protocol was selected due to its capacity in separating the equilibrium response of the material from rate-dependent dissipative phenomena, associated with viscous effects. The chosen metrics, including equilibrium stress and stress work, dissipated energy during loading and unloading, and normalized stress decay, provided a rich landscape to compare the simulant materials. Notably, all comparisons extends beyond a qualitative assessment by resorting to rigorous statistical analysis. This enabled us to identify aspects of similarity and differences with respect to the mechanical behavior of the native tissue, ultimately resulting in a critical assessment of the ideal simulant for brain phantoms. This point, together with some limitations and insights in future integrations of this study, are provided below.

### 4.1. Summary of the biomechanical fidelity of the candidate materials

To provide an overall interpretation of the experimental findings, Table 2 summarizes the extent to which each investigated material reproduces the mechanical response of porcine brain tissue under compression and tension. The comparison first considers the overall stress magnitude during loading and unloading, for which CH and Sylgard exhibit metrics closer to those of brain tissue at comparable stretch levels. The remaining comparison is organized according to the response features investigated in this work, namely equilibrium response, inelastic effects, and normalized stress relaxation. For each metric, the table distinguishes between the agreement in terms of magnitude and the ability to reproduce the rate-dependent behavior observed in brain tissue. Symbols are assigned based on the statistical comparisons reported in the previous sections and indicate whether agreement with the brain response is observed consistently across the investigated conditions, only under selected test conditions, or not observed. This qualitative overview provides information on the fidelity of each candidate material, i.e., its mechanical match with native brain tissue, with respect to the specific mechanical features addressed in the present study.

**Table 2.**
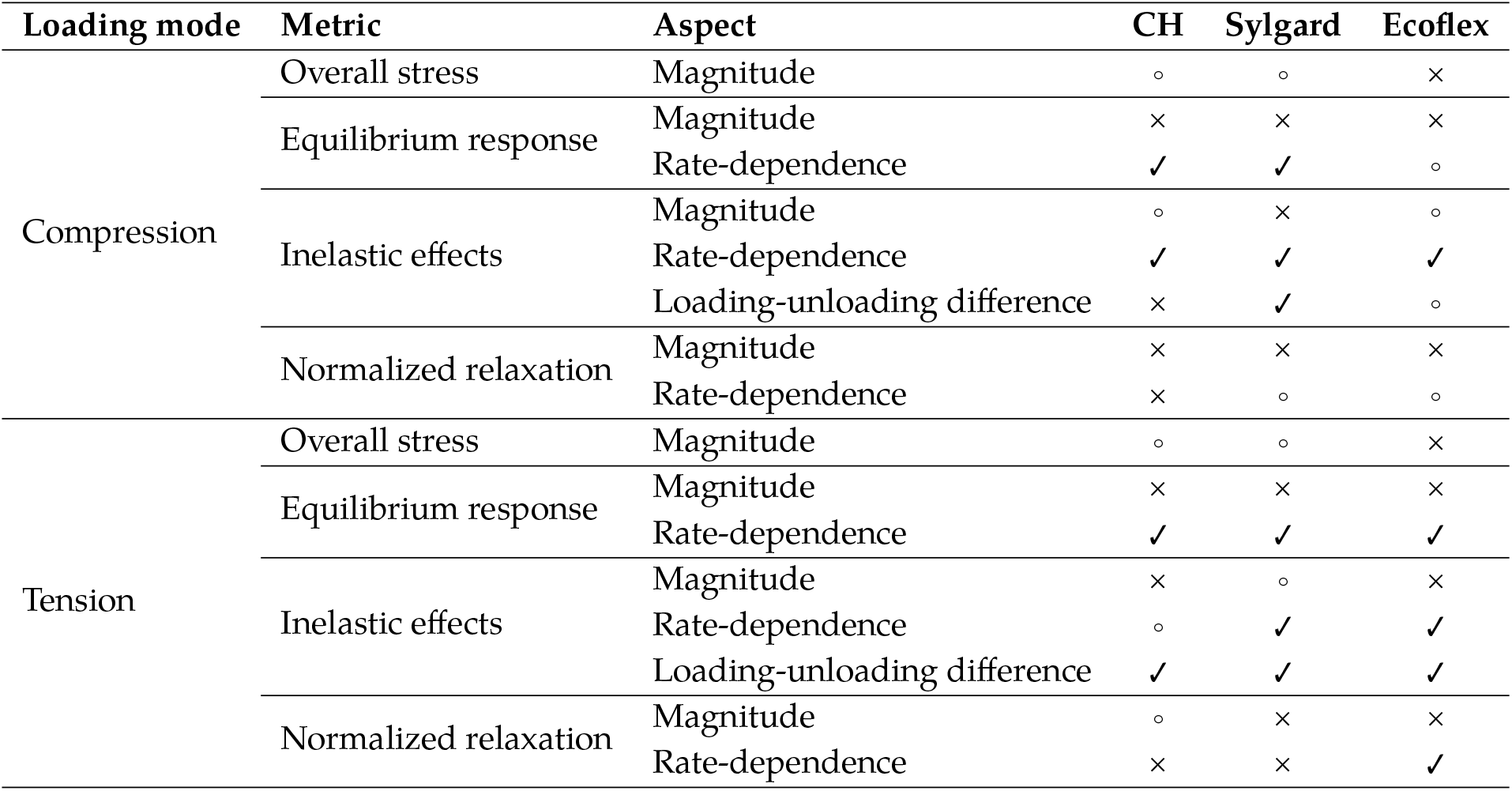
Qualitative summary of the fidelity of the investigated materials compared to porcine brain tissue in compression and tension. Symbols indicate the degree to which each material reproduces specific mechanical features of the brain: agreement (✓); agreement under selected conditions, such as deformation level, stretch rate, and energy-based quantity (°); no agreement (×). For rate-dependence and loading-unloading difference, the symbols reflect the consistency of the simulant behavior with that observed in native tissue, regardless of whether these effects are present or absent.

Overall, no single material fully reproduces the mechanical response of brain tissue across all the investigated metrics and loading conditions. Nevertheless, CH and Sylgard emerge as the most promising candidates among the investigated materials, although their degree of fidelity depends on the specific feature considered. These findings suggest that the selection of a brain tissue simulant should be guided by the mechanical requirements of the intended application rather than by a single measure of overall fidelity.

### 4.2. Research gaps and perspectives for brain tissue simulants

Sylgard was already established in the literature as a stable and reliable simulant material for brain tissue. The results obtained in this study further strengthen the claim, with a high level of agreement displayed for several of the mechanical metrics considered. However, it essentially remains a monophasic material, and as a result, unsuitable to replicate the biphasic nature of brain tissue arising from the interaction between solid and fluid. In fluid-saturated porous materials, mechanical deformation and interstitial fluid flow are coupled. This means that forces and deformations affect the pressure of the interstitial fluid, while pressure gradients result in fluid flow and subsequent mechanical deformation [83]. This is relevant by a mechanical point of view, since dissipative phenomena arising from this interaction add to the intrinsic viscoelasticity of brain tissue [84]. In addition, the biphasic nature of brain tissue needs to be replied in selected application of the simulant materials, related for instance, to the field of drug delivery models [85]. In this regard, the highly hydrated porous structure of CH, consisting of a deformable solid matrix and free-flowing interstitial fluid [74], appears as a physically more realistic simulant. Furthermore, it is worth mentioning that the CH composition can be tuned, for instance by changing the total volume fraction of polymer, the relative fractions of polymer constituents, or the freeze-casting protocol, in order to increase the mechano-fidelity to brain tissue for the intended application. This is reflected in changes affecting both the macroscopic mechanical behavior and the intrinsic porous microstructure of the material [86].

The characterization framework adopted here could be extended by including properties related to the porous structure and fluid mobility, such as the effective porosity and permeability coefficients. In biphasic models, the former identifies the free water fraction, i.e., the volume of fluid initially present in a saturated porous material that can be mechanically displaced, e.g. by consolidation. The latter describes the constitutive response of a porous material associated to the motion of fluid through the network. The effective porosity could be experimentally quantified through incremental consolidation under drained confined compression, as recently proposed for highly hydrated ultrasoft hydrogels [86]. Moreover, mechanical tests performed over different characteristic length and time scales could potentially offer insights in the distinction of fluid-related from viscous dissipation [74, 84]. Overall, these approaches should lead to the identification of additional criteria for designing and assessing future brain tissue simulants.

#### A. Unconfined uniaxial deformation of biphasic porous materials

The solution of the IBVP of unconfined compression is beyond the scope of this work. Nevertheless, some insights are here provided, since these help to: (i) appreciate why the deformation and stress fields are generally inhomogeneous during mechanical experiments on fluid-saturated solids; and (ii) demonstrate conditions under which these fields become homogeneous under the introduced kinematic ansatz. This is done by recalling the fundamental balance equations in quasi-static conditions, which are the balances of mass and linear momentum, complemented by boundary conditions. A minimal set of assumptions is introduced, in addition to those on kinematics already stated. Namely, we consider a fully saturated mixture consisting of intrinsically incompressible solid and fluid constituents, each free to move at different velocities [87]. Furthermore, we assume that: (i) the process is quasi-static and body forces are negligible; (ii) the solid is isotropic and homogeneous, and its contact with the platens is frictionless; (iii) the fluid flux is purely radial, ***w*** = *w*(*r, t*) ***e***_*r*_, consistently with the axial homogeneity of the kinematic ansatz and with impermeable platens. The ambient pressure *p*_0_ acts uniformly on all surfaces exposed to the bath (Fig. 1). It is therefore removed from the outset and the force *f*_*z*_ recorded by the load cell is understood as tared accordingly.

The total Cauchy stress tensor can then be expressed as

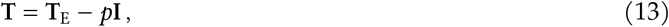

where **T**_E_ represents the extra stress component, and *p* is commonly referred to as the interstitial fluid pressure. Rigorously, *p* emerges as a Lagrange multiplier to enforce mixture saturation and phase incompressibility [88].

In the radial direction, the balance equations read [89]

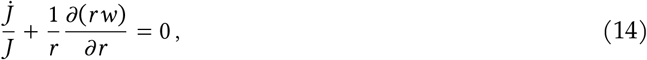

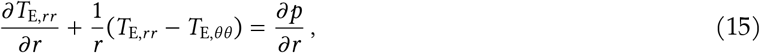

where *w* = *w*(*r, t*) is the radial component of the *discharge velocity*, and we have utilized the kinematic identity 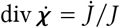 [90].

The following boundary conditions apply:

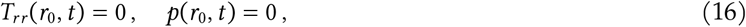

where the second condition expresses continuity of the interstitial pressure with the bath at ambient pressure.

Let us first consider the loading phase associated with a non-zero local volume change, 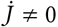, whose sign can in general be positive or negative throughout the protocols considered in this work. In this scenario, Eq. (14) demonstrates that non-zero radial filtration must exist over at least part of the cross-section. In a biphasic medium, this relative fluid motion is proportional to the pressure gradient *∂p*/*∂r*; the simplest case is the linear proportionality implied by Darcy’s law [88]. Hence, wherever *w* ≠ 0, a non-zero radial gradient emerges on the right-hand side of Eq. (15), dictating that the extra stress field cannot be spatially homogeneous. This does not rigorously prove that the total stress is inhomogeneous. A non-uniform pore-pressure field can, in principle, be balanced by a corresponding non-uniform extra stress field, leaving the total stress spatially homogeneous while still satisfying the traction boundary condition (16)_1_.

Now we proceed with the identification of relevant scenarios where deformation and stress fields are instead spatially homogeneous, i.e., independent of the radial coordinate. First, we consider the long-term thermodynamic equilibrium, *t* → ∞, for which *w*(*r, t*) → 0 and 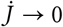 [86]. In the current experimental realization this condition is approximated after a sufficiently long holding time *t*^eq^. The mass balance equation (14) is trivially satisfied. Consistently with vanishing fluid flux, the pressure gradient *∂p*/*∂r* vanishes as well; using the boundary condition (16)_2_, this yields a uniform pressure field *p*(*r, t*^eq^) = 0. We cannot prove that this implies spatial homogeneity in the stress and deformation fields without additional constitutive assumptions. However, it can be shown that this condition admits a homogeneous solution. Taking

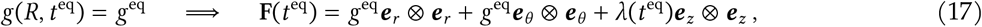

yields an equilibrium Jacobian *J* ^eq^ = *λ*(*g*^eq^)^2^. This is consistent with mass balance without requiring *J* ^eq^ = 1, showing that a homogeneous volume change is compatible with thermodynamic equilibrium. Under isotropy, the deformation gradient in Eq. (17) implies 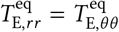. Because *p* = 0, the traction condition (16)_1_ requires 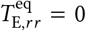 at *r* = *r*_0_, and hence everywhere by homogeneity. Therefore 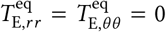. The radial momentum balance (15) is then trivially satisfied, and the total stress tensor is spatially homogeneous and uniaxial.

Next, we consider an undrained limit in which the fluid discharge is zero everywhere throughout the loading process, i.e., *w*(*r, t*) = 0 for all *r* and *t*. In the current work this limit is relevant for hydrophobic polymeric materials, for which *p* is simply the Lagrange multiplier enforcing the incompressibility constraint. In this scenario, 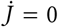, and Eq. (14) can be integrated with the initial condition *J* (*R*, 0) = 1 to yield *J* = 1 throughout the whole deformation process. Using the explicit Jacobian of the deformation gradient given by Eq. (2), one obtains

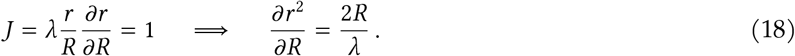

Integrating with respect to the radial coordinate under the symmetry condition *r*(0, *t*) = 0 yields

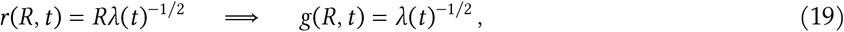

which reduces the deformation gradient in Eq. (2) to the spatially homogeneous form

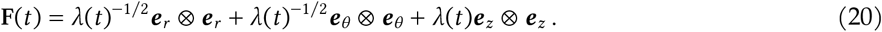

Since the material is homogeneous and isotropic, this produces a homogeneous extra stress field with *T*_E,*rr*_ = *T*_E,*θθ*_. The radial momentum balance (15) then reduces to *∂p*/*∂r* = 0, so that the pressure is spatially uniform. The value of *p* is fixed by the traction condition (16)_1_, which gives *p* = *T*_E,*rr*_. The total stress tensor is therefore spatially homogeneous and uniaxial, with *T*_*rr*_ = *T*_*θθ*_ = 0 and *T*_*zz*_ = *T*_E,*zz*_ − *T*_E,*rr*_.

In summary, we have shown that stress and deformation fields are generally inhomogeneous, and identified two limiting regimes where homogeneity is restored. In these special cases, Eq. (5) simplifies to

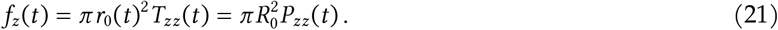

#### B. Dissipation in fluid-saturated materials

The dissipation inequality (7) for a biphasic fluid-saturated material takes the following explicit form [79, Eq. (110)]

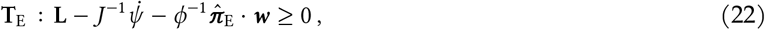

where *ϕ* is the porosity, 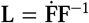 is the spatial velocity gradient of the solid, and *ψ* is the strain energy per unit reference volume of the solid. The stress power introduced in (7) is defined as *p*_int_ = **T**_E_ : **L**. The effective term 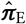 is related to the momentum exchange and can be interpreted as the local interaction force between interstitial fluid and solid skeleton [88]. Under the assumption of an inviscid fluid, the last term in (22), proportional to the discharge velocity ***w***, is the only transport-related dissipative term of the model. A standard way to enforce its non-negativity is to define

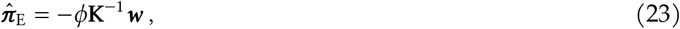

where **K**^−1^ represents a positive definite hydraulic resistivity or inverse permeability tensor [88]. This way, the transport-related dissipation appearing in the inequality (7) is

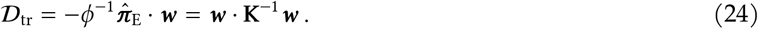

As stated in Sect. 2.1.2, only the volume-averaged nominal stress power

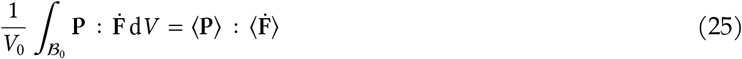

can be computed experimentally. To establish a relation with inequality (22), we define the effective stress power per unit reference volume of the solid as 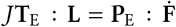. Recalling the stress decomposition introduced in Eq. (13) and the kinematic identity 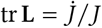[90], we can write

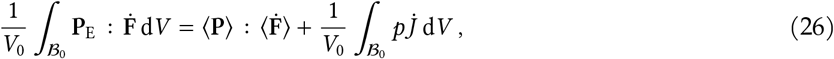

where the effective nominal stress is **P**_E_ = **P** + *pJ* **F**^−⊤^.

For the unconfined uniaxial deformation considered here, the effective nominal stress power can be expressed as

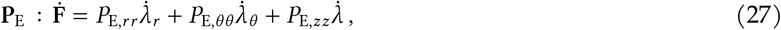

where the principal stretches *λ*_*r*_, *λ*_*θ*_, *λ* appear explicitly in Eq. (2) in front of the respective dyadic products. The volume-averaged nominal stress power reads

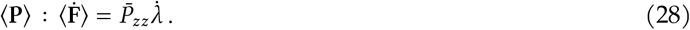

The isotropic transport-related dissipation for radial flow reduces to

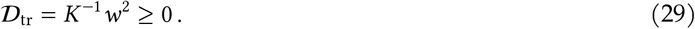

Multiplying inequality (22) by *J*, integrating over *B*_0_, dividing by *V*_0_, and collecting the terms depending on fluid-related variables, the resulting volume-averaged inequality is

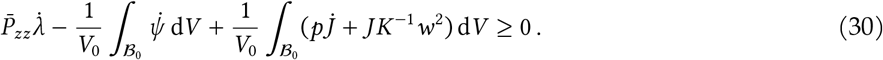

The last integral in inequality (30) is not accessible experimentally. However, we can prove that it cancels out in the current experiment, if the additional assumption of Darcy’s flow is taken. Accordingly, the transport-related dissipation in (29) can be written as [88]

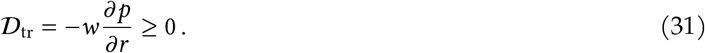

Using the mass balance (14) and (31), the second integrand in (30) satisfies

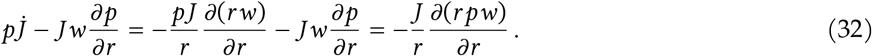

Through the relationship d*v* = *J* d*V* = 2*πrh*_0_ d*r*, we integrate the term above over the current configuration:

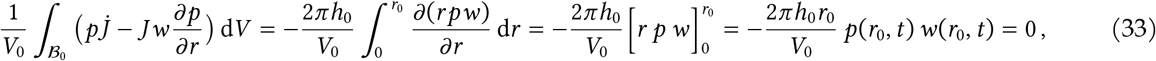

where the contribution at *r* = 0 vanishes by symmetry and the last equality follows from the boundary condition (16)_2_. The volume-averaged dissipation therefore reads

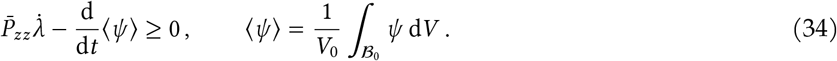

For the hydrophobic polymers considered in this work, this is immediate without any boundary condition, since 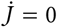 and *w* = 0 make both terms appearing in the second integrand of inequality (30) zero.

#### C. Stress relaxation criterion

The assessment of stress relaxation is fundamental for defining equilibrium stresses, yet the procedure to identify an equilibrium-like state is rarely described in the literature. In the present work, we introduced both quantitative and qualitative metrics to identify the onset of an equilibrium-like state during the holding phase following each prescribed stretch increment in the multi-step protocols. These criteria were applied to preliminary experiments on brain tissue, described in detail in Sect. S1.1 of the Supplementary Material available online. Figure 18 shows an example of the application of the relaxation criteria in compression; the final unloading step is considered here for illustrative purposes (Fig. 18a).

**Figure 18.**
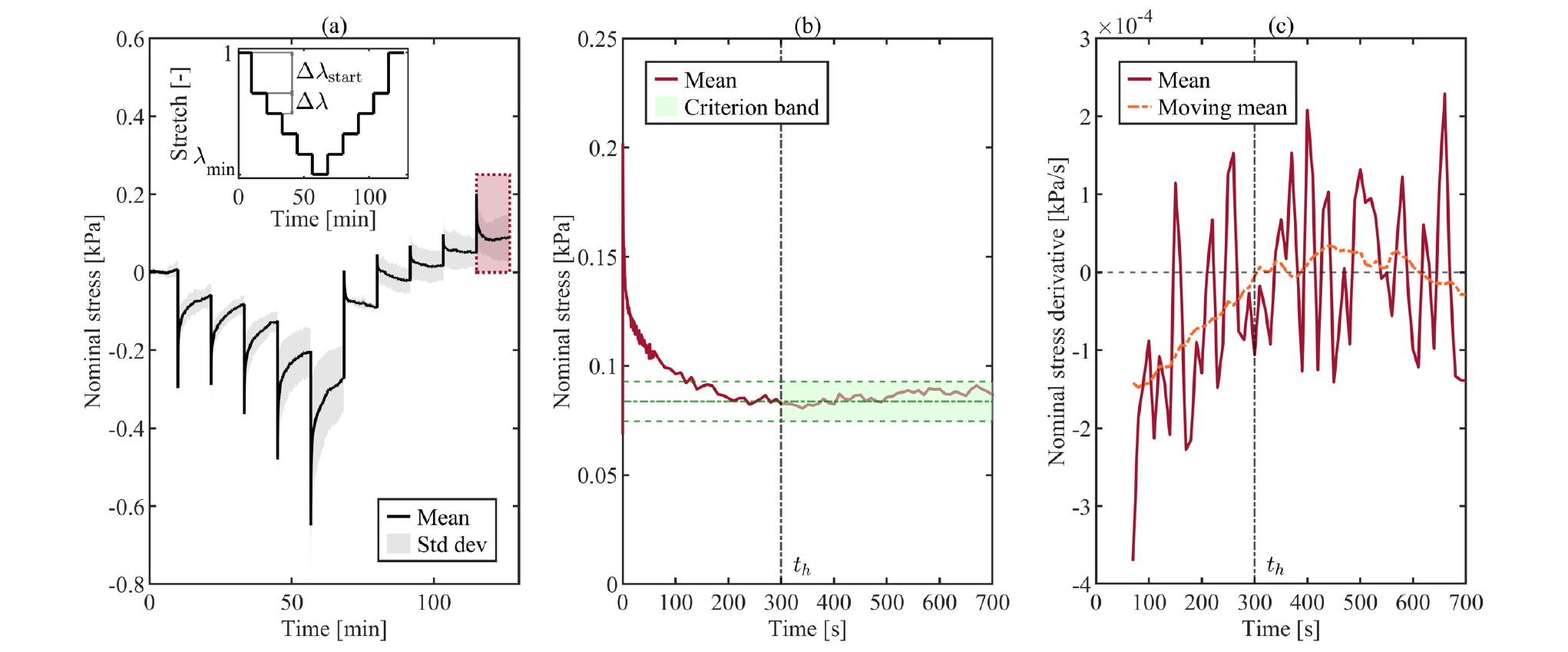
Illustration of the quantitative and qualitative stress relaxation criteria. (a) Evolution of the mean nominal stress over time and preliminary stretch-controlled protocol (inset), with the selected step for relaxation assessment highlighted in red; (b) evolution of the mean nominal stress over time during the selected step, with the application of the quantitative metric based on the threshold band highlighted in green; (c) evolution of the mean first derivative of the nominal stress over time during the selected step, alongside the moving mean for improved interpretation of the qualitative metric. The holding time *t*_h_ is highlighted in both figures (b) and (c). The data is taken from a compression experiment on brain tissue performed with a preliminary protocol with constant holding times of 700 s. Several holding phases during loading do not satisfy the relaxation criteria and therefore had to be extended in the final protocol.

From a quantitative point of view, relaxation was considered to be achieved at the earliest time after which the mean nominal stress remained within a threshold range, defined by the load-cell resolution normalized by the average initial cross-sectional area of the specimens, for the remainder of the holding phase (Fig. 18b). The corresponding stress threshold was defined as

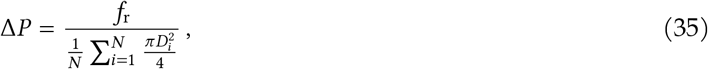

where *f*_r_ is the load-cell resolution (1 mN), *N* is the number of specimens, and *D*_*i*_ is the diameter of the *i*-th specimen. The criterion was formulated in terms of nominal stress rather than force to account for differences in specimen geometry among the tested materials.

From a qualitative perspective, the mean first derivative of the nominal stress was required to approach and remain close to zero until the end of the holding phase (Fig. 18c). For clarity, the derivative is shown from 60 s after the stress peak onward, in order to focus on the portion of the response relevant to the assessment of the relaxation criterion. A smoothed trend is also shown to improve the readability of the derivative signal.

The holding time *t*_h_ was defined as the first time at which both the quantitative and qualitative conditions were fulfilled. The resulting values of *t*_h_ at each prescribed stretch levels were used to establish the holding durations in the final multi-step protocols described in Sect. 2.2.3.

## Acknowledgements

E.B. acknowledges that this research was made possible by a grant from the Fondazione Ing. Aldo Gini. M.T. is grateful to Dr. Maximilian P. Wollner for the fruitful discussions on the theoretical aspects of this paper.

## Supplementary Material

This Supplementary Material provides additional methodological details, figures, and tables supporting the results presented in the main manuscript. Specifically:

- Section S1 provides further information on the experimental methodology. This includes the preliminary analyses performed using the relaxation criterion defined in the main manuscript, which guided the definition of the final testing protocols.
- Sections S2 and S3 present additional figures and tables for the compression and tension experiments, respectively.

As in the main manuscript, the supplementary results are organized by loading mode. Within each loading mode, the results are further organized according to the quantitative metrics used to assess simulant fidelity. Results for each simulant material (CH, Sylgard and Ecoflex) are presented following the same structure adopted for brain tissue in the main text. In addition, the supplementary results include comparisons between brain tissue and simulant materials at the low stretch rate, which complement the high-rate comparisons presented in the main manuscript.

For the equilibrium response, the supplementary figures report, for each simulant material, the median nominal stress–time response with the interquartile range (IQR), the equilibrium nominal stress–stretch curves with the IQR, and a quantitative comparison of equilibrium stresses between the two stretch rates at representative stretch levels. These levels correspond to *λ*_1_ = 0.95 and *λ*_5_ = 0.85 in compression, and *λ*_1_ = 1.05 and *λ*_5_ = 1.15 in tension. Statistical significance was assessed using the Mann–Whitney U (MWU) test.

For the inelastic response, the supplementary figures show, for each simulant material, the median nominal stress–stretch response. Shaded areas indicate the equilibrium stress work and the dissipation accumulated during loading and unloading. The figures also include a quantitative comparison of the energy-based metrics between the two stretch rates. Statistical significance was assessed using the MWU test. In addition, the accumulated dissipation during loading and unloading was compared at each stretch rate using the one-tailed Wilcoxon signed-rank (WSR) test.

For normalized stress relaxation, the supplementary figures present the normalized nominal stress evolution during the holding phases for each simulant material, together with quantitative comparisons between the two stretch rates at two representative loading steps. these correspond to loading step 1, at *λ* = 0.95 in compression and *λ* = 1.05 in tension, and loading step 5, at *λ* = 0.85 in compression and *λ* = 1.15 in tension. Statistical significance was assessed using the MWU test. In addition, the supplementary tables provide the complete comparison of normalized stress values at six selected time points (1, 10, 30, 60, and 300 s after the onset of the holding phase, as well as at the end of relaxation) for both representative loading steps at both stretch rates.

## S1. Experimental methodology

### S1.1. Preliminary analyses for protocol definition

To define the final testing protocols used in compression and tension, preliminary multi-step stress relaxation tests were conducted on porcine brain tissue.

The protocol design was informed by a review of the literature on single-step stress relaxation experiments performed on human, porcine, and bovine brain tissue under both compression and tension. To minimize the risk of damage to the tissue microstructure, the maximum engineering strain in tension was limited to 15%. This limit was selected on the base of recent evidence indicating that axonal damage in white matter may occur at tensile strains of approximately 18% [S1]. On the other hand, according to Reiter et al. [S2], porcine brain tissue can withstand compressive strains of up to 30% without permanent damage. However, Eskandari et al. [S3] suggested that brain tissue integrity may be impaired at compressive strains exceeding the 18% threshold associated with brain injury. Therefore, a maximum engineering strain of 15% was also selected for the compression protocol.

Reported holding times for single-step relaxation tests conducted at comparable maximum strain levels (up to 30% in compression and 15% in tension) [S1, S2, S4–S18] were found to be as high as 500–600 s in compression [S2, S9, S15] and 300 s in tension [S1, S17]. The 3500-s holding time reported by Cheng and Bilston [S5] was not considered because the authors reported limitations related to tissue damage and dehydration. The available studies differ from the present work in terms of factors such as preconditioning, the application of multiple loading modes to the same specimen, and testing in air. Consequently, preliminary testing was required to assess the relaxation behavior of the samples under the present experimental conditions.

A conservative holding time of 700 s was chosen for each relaxation step, exceeding the durations reported in the literature. The preliminary protocol consisted of five stepwise loading ramps applied at a constant stretch rate and separated by holding phases. After immersion in the fluid bath, specimens were held for 10 min at *λ* = 1 to allow mechanical stabilization before the first loading ramp. The deformation was then applied through successive stretch increments of Δ*λ* = 0.025, except for the first increment, for which Δ*λ*_start_ = 0.05 was used. Accordingly, the stretch progressively decreased to a minimum value *λ*_min_ = 0.85 in compression, and increased to a maximum value *λ*_max_ = 1.15 in tension. The larger first increment was introduced to reduce the likelihood that equilibrium forces at the lowest deformation level would approach the load-cell resolution, given the softness of brain tissue. After reaching *λ*_min_ in compression or *λ*_max_ in tension, a symmetric stepwise unloading sequence was applied. Two constant stretch rates were considered: 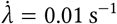 and 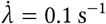, hereafter denoted to as the ‘low’ and ‘high’ stretch rates, respectively. The high stretch rate was selected based on the limitations of the testing device, which permitted a maximum displacement rate 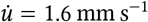. For the specimen geometry considered, this corresponded to a maximum attainable stretch rate of approximately 0.32 s^−1^. The preliminary protocol was applied in both compression and tension only at the high stretch rate, as this was deemed to represent the most critical condition for relaxation. Indeed, higher loading rates are expected to produce longer relaxation times; thus, this condition was used to calibrate the holding durations for the final protocols described in the main manuscript.

The preliminary relaxation data were evaluated using the criterion defined in the main manuscript. Based on this, a holding time of 700 s proved sufficient to achieve relaxation during all unloading steps in compression, as well as during both loading and unloading under tension. In contrast, 700 s was insufficient to ensure complete relaxation during the compression loading steps. Therefore, an additional preliminary compression protocol was conducted to refine the loading-phase holding times while limiting the overall experimental effort. This additional protocol included only the loading path, using the same sequence of stretch levels down to the minimum stretch *λ*_min_ = 0.85. Each loading increment was followed by an extended holding phase of 20 min, without performing any unloading. The results from this additional assessment were combined with those from the initial preliminary protocol to establish the final multi-step compression protocol presented in the main manuscript.

### S1.2. Supplementary figures on the experimental setup

**Figure S1.**
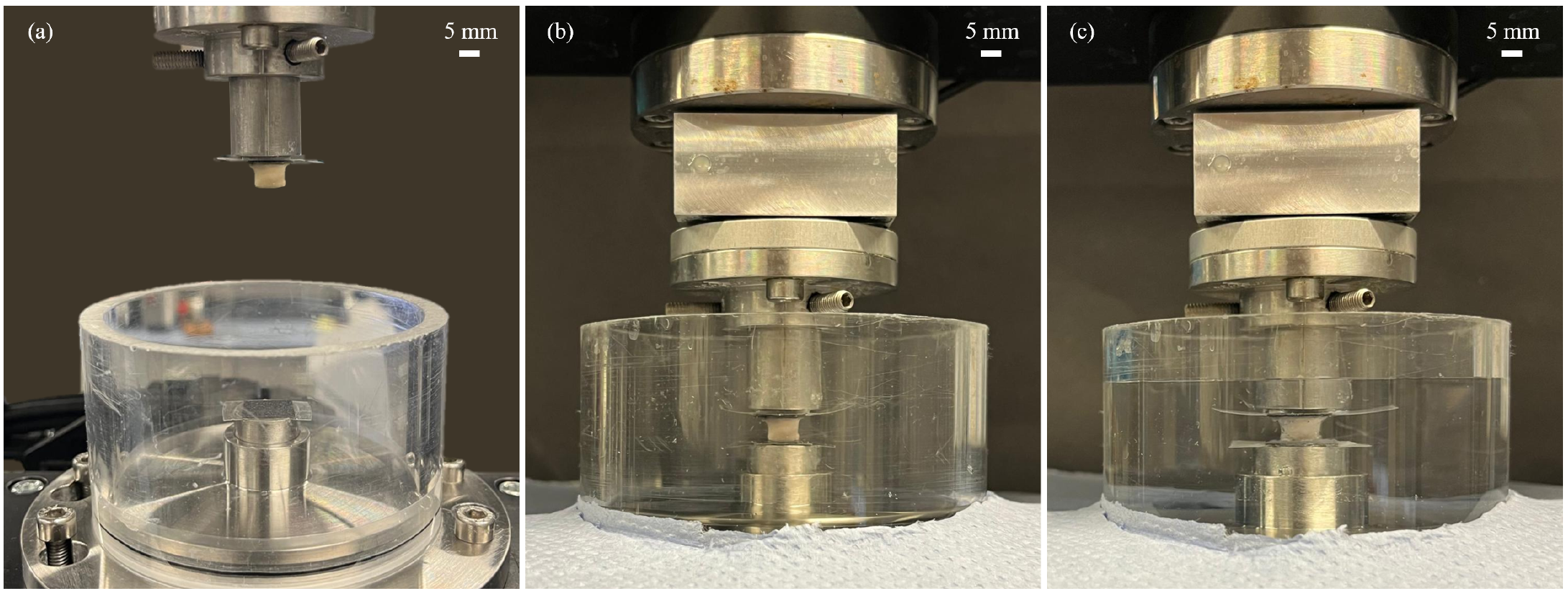
Experimental setup for mechanical testing. (a) Configuration at the beginning of the approaching procedure; (b) configuration at the end of the approaching procedure, with the sample brought into contact with the lower adapter; (c) configuration after filling the bath with testing fluid, with the sample fully submerged.

**Figure S2.**
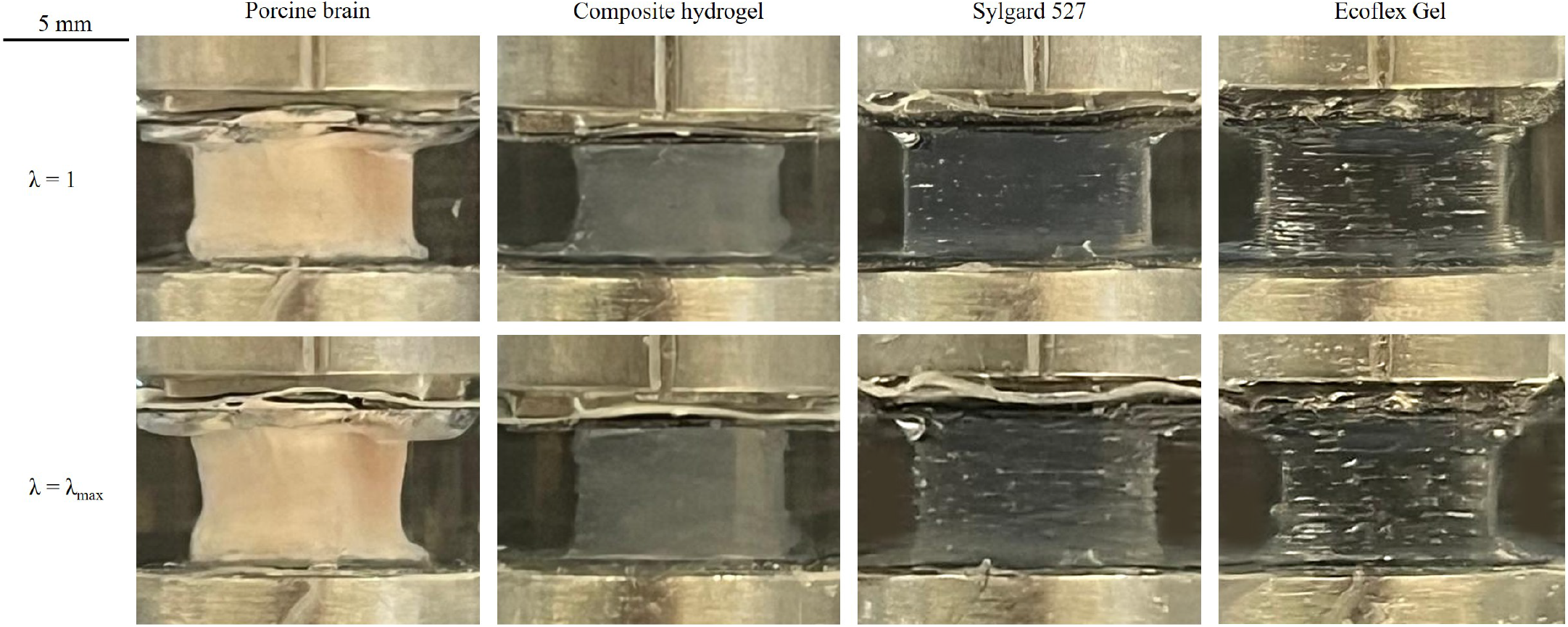
Representative configuration of the four tested materials (porcine brain tissue, composite hydrogel (CH), Sylgard, and Ecoflex during tension experiments. The first row refers to the undeformed configuration (*λ* = 1), while the second one refers to the configuration under maximum tension (*λ* = *λ*_max_ = 1.15).

## S2. Compression experiments: additional figures and tables

### S2.1. Equilibrium response

**Figure S3.**
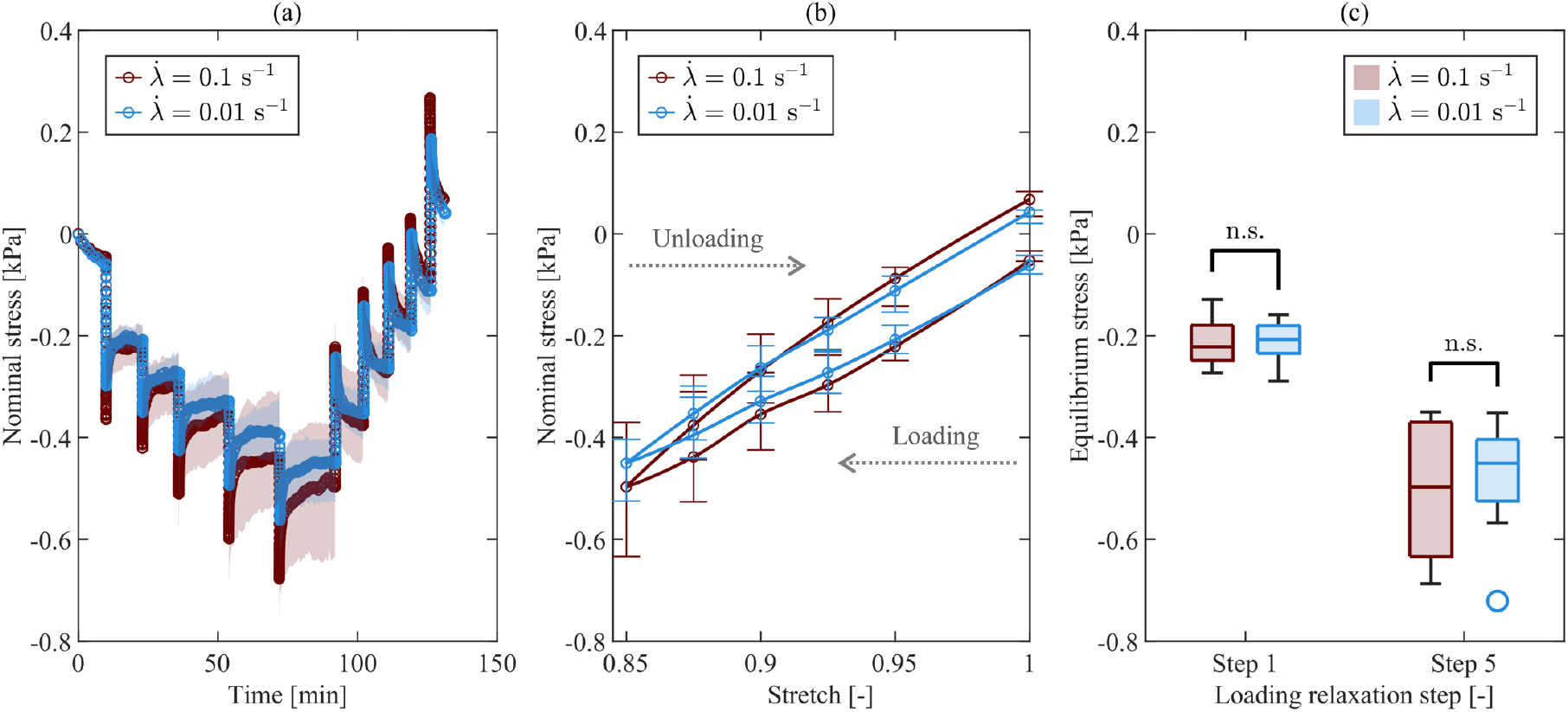
CH equilibrium response in compression at high (red) and low (blue) stretch rates. (a) Median nominal stress–time with IQR (shaded areas); (b) equilibrium stress–stretch relation with IQR (bars); (c) comparison of equilibrium stress at two representative loading steps (Step 1: *λ* = 0.95; Step 5: *λ* = 0.85) between stretch rates, showing no significant differences.

**Figure S4.**
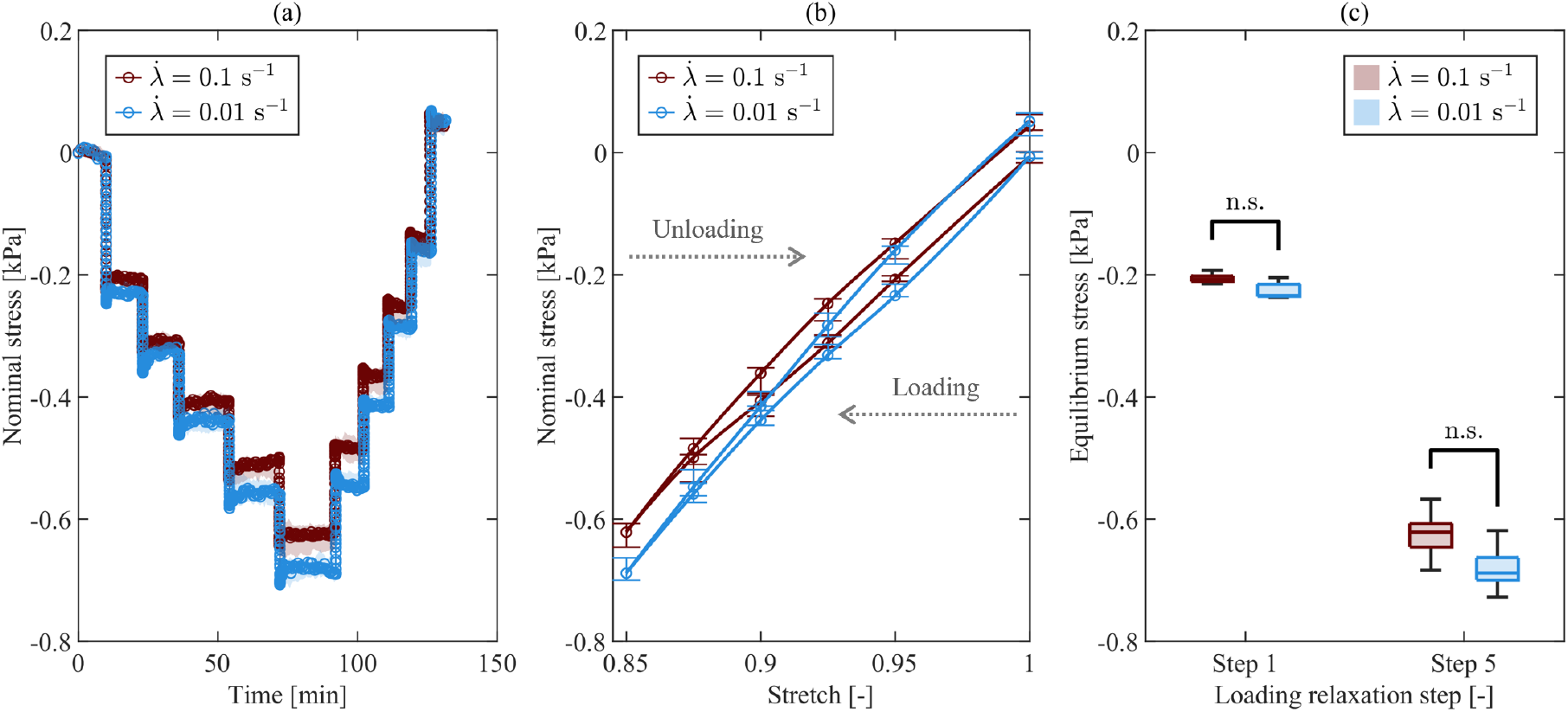
Sylgard equilibrium response in compression at high (red) and low (blue) stretch rates. (a) Median nominal stress–time with IQR (shaded areas); (b) equilibrium stress–stretch relation with IQR (bars); (c) comparison of equilibrium stress at two representative loading steps (Step 1: *λ* = 0.95; Step 5: *λ* = 0.85) between stretch rates, showing no significant differences.

**Figure S5.**
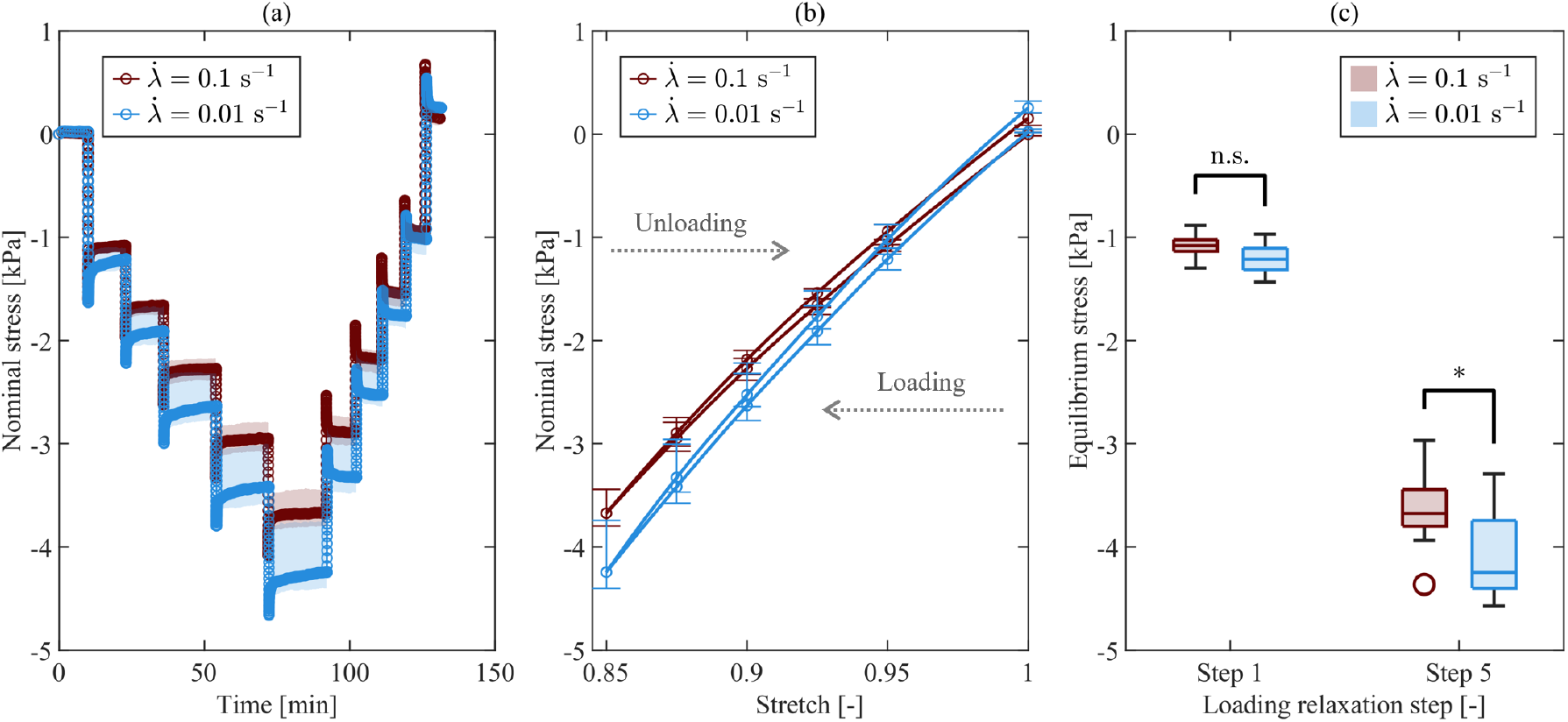
Ecoflex equilibrium response in compression at high (red) and low (blue) stretch rates. (a) Median nominal stress–time with IQR (shaded areas); (b) equilibrium stress–stretch relation with IQR (bars); (c) comparison of equilibrium stress at two representative loading steps (Step 1: *λ* = 0.95; Step 5: *λ* = 0.85) between stretch rates, showing significantly higher equilibrium stresses at higher rates in Step 5 (*p<0.05).

**Figure S6.**
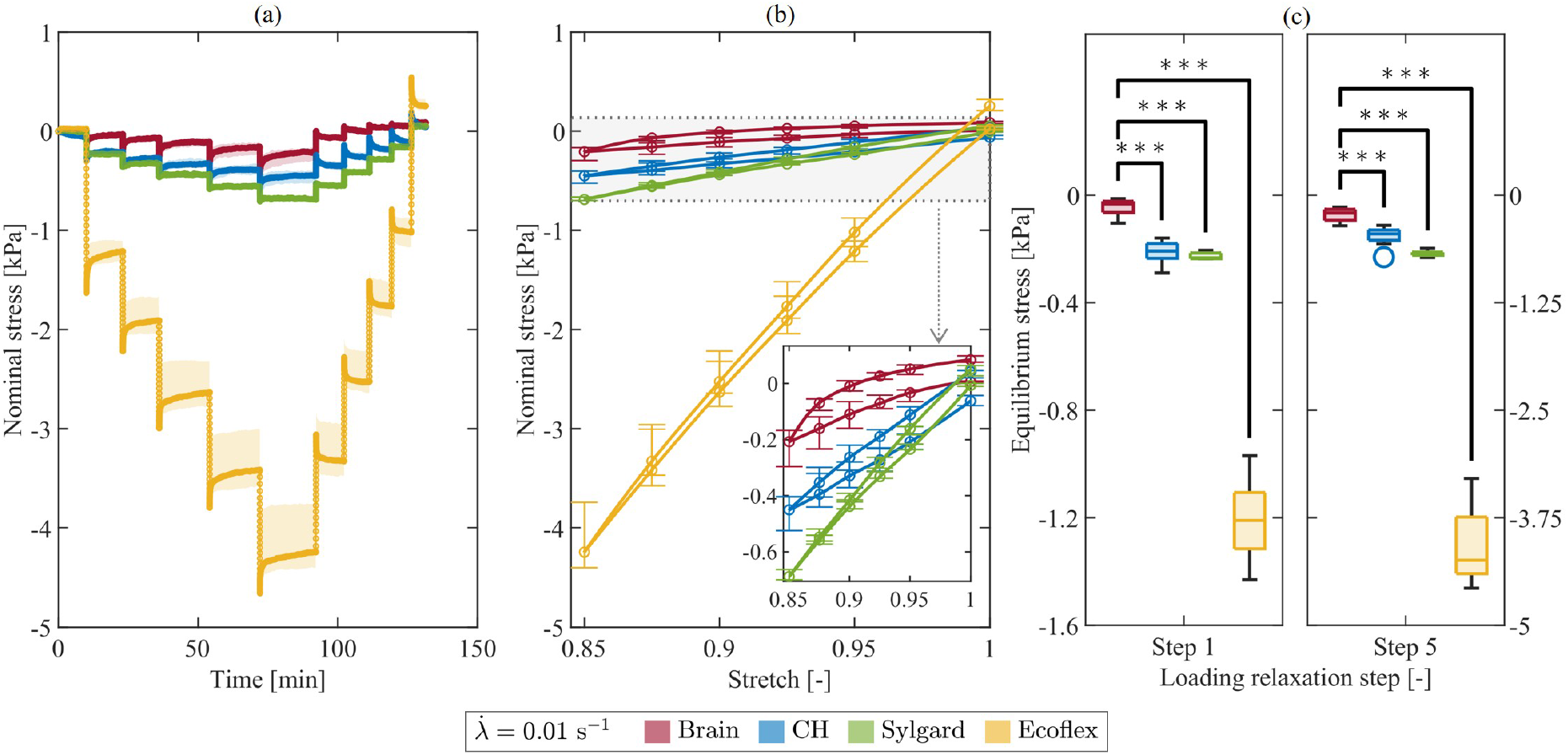
Comparison of the equilibrium response in compression across all materials at the low stretch rate. (a) Median nominal stress–time with IQR (shaded areas); (b) equilibrium stress–stretch relation with IQR (bars). The inset enlarges the stress range relevant to brain tissue, CH, and Sylgard to facilitate comparison; (c) comparison of equilibrium stress at two representative loading steps (1: *λ* = 0.95; 5: *λ* = 0.85) between brain tissue and each simulant material (***p<0.001).

### S2.2. Inelastic effects

**Figure S7.**
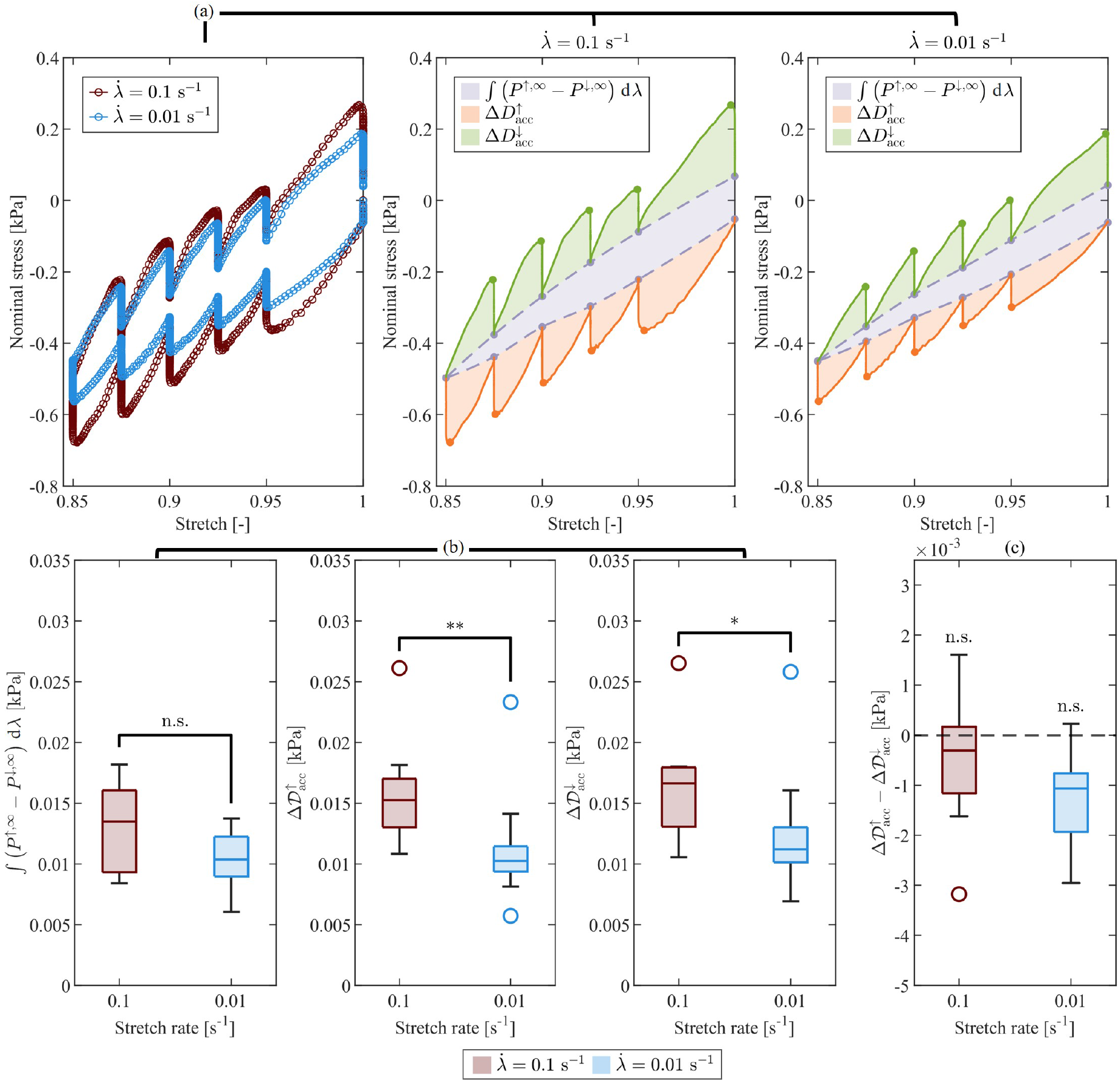
CH inelastic effects in compression at high (red) and low (blue) stretch rates. (a) Median nominal stress–stretch response with shaded areas representing the stress work associated with the equilibrium relation (gray), and accumulated dissipation during loading (orange) and unloading (green), at the two stretch rates; (b) comparison of the energy-based metrics between stretch rates (MWU test), showing rate-dependent accumulated dissipation (*p<0.05; **p<0.01) but no evidence of significant rate dependence in the equilibrium stress work; (c) comparison between loading and unloading accumulated dissipation (one-tailed WSR test), indicating no evidence of significantly higher dissipation during loading at both stretch rates.

**Figure S8.**
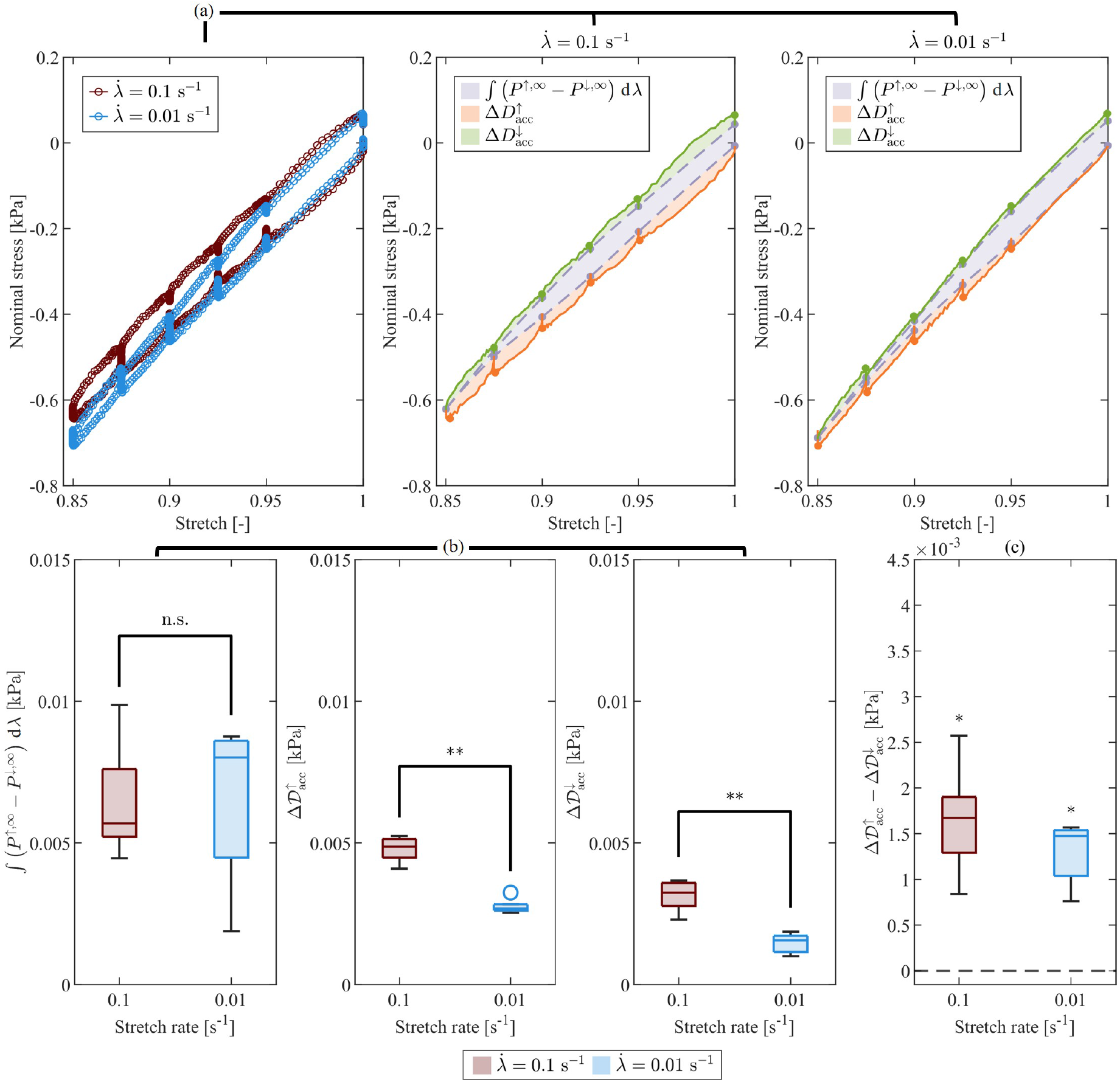
Sylgard inelastic effects in compression at high (red) and low (blue) stretch rates. (a) Median nominal stress–stretch response with shaded areas representing the stress work associated with the equilibrium relation (gray), and accumulated dissipation during loading (orange) and unloading (green), at the two stretch rates; (b) comparison of the energy-based metrics between stretch rates (MWU test), showing rate-dependent accumulated dissipation (**p<0.01) and no evidence of significant rate dependence in the equilibrium stress work; (c) comparison between loading and unloading accumulated dissipation (one-tailed WSR test), indicating significantly higher dissipation during loading at both stretch rates (*p<0.05).

**Figure S9.**
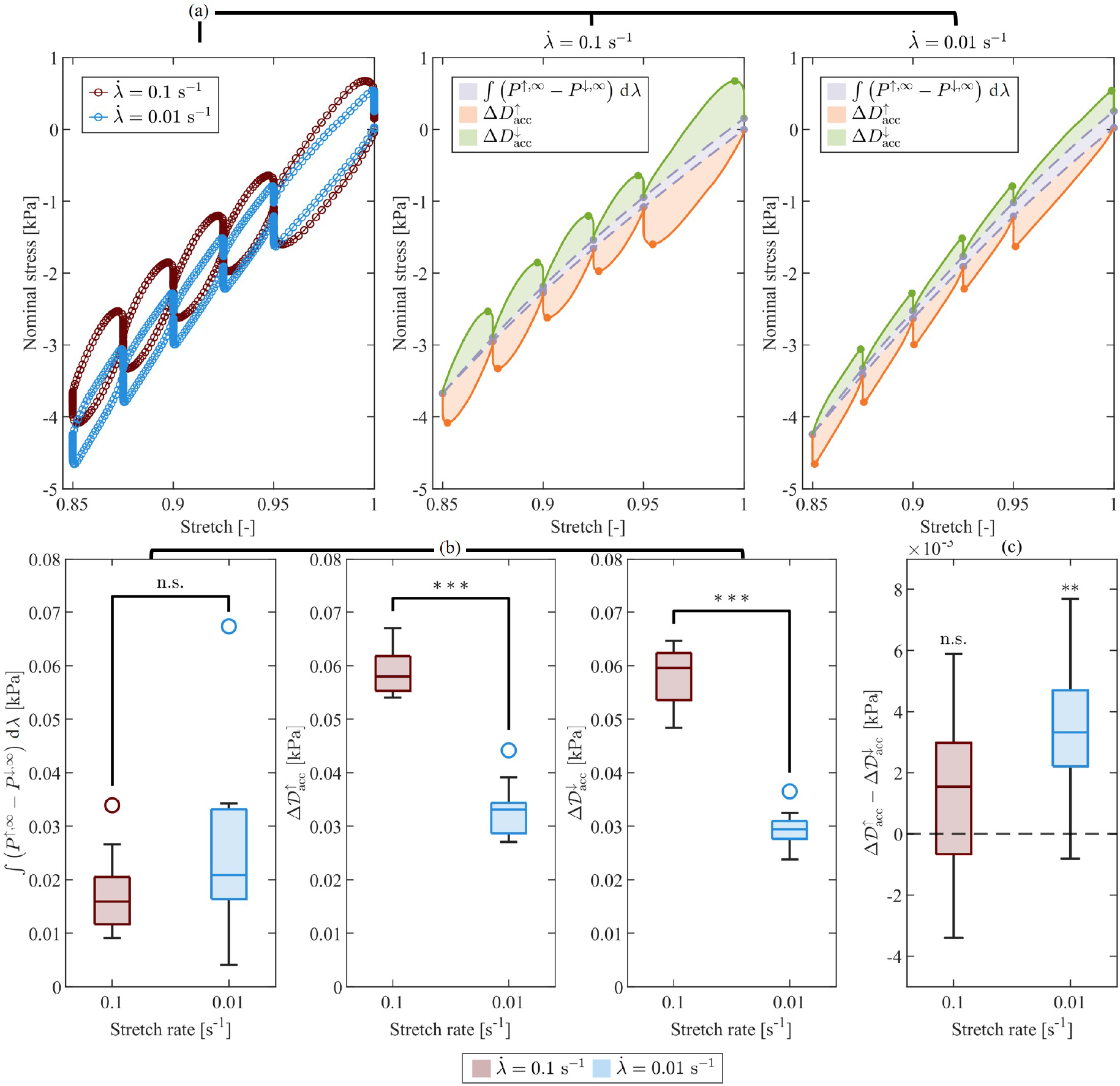
Ecoflex inelastic effects in compression at high (red) and low (blue) stretch rates. (a) Median nominal stress–stretch response with shaded areas representing the stress work associated with the equilibrium relation (gray), and accumulated dissipation during loading (orange) and unloading (green), at the two stretch rates; (b) comparison of the energy-based metrics between stretch rates (MWU test), showing rate-dependent accumulated dissipation (***p<0.001) and no evidence of significant rate dependence in the equilibrium stress work; (c) comparison between loading and unloading accumulated dissipation (one-tailed WSR test), indicating significantly higher dissipation during loading at the low stretch rate (**p<0.01).

**Figure S10.**
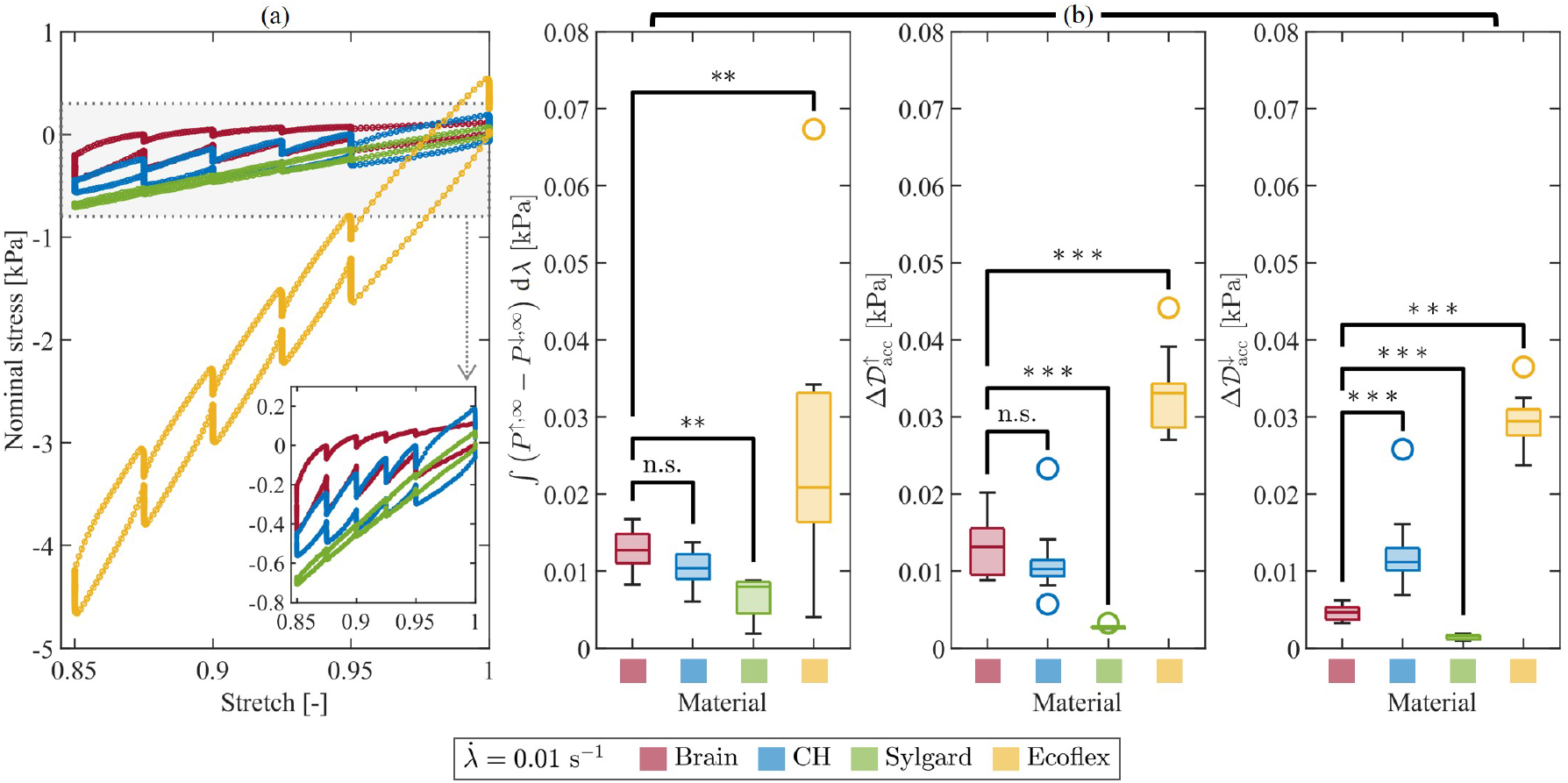
Comparison of inelastic effects in compression across all materials at the low stretch rate. (a) Median nominal stress–stretch response. The inset enlarges the stress range relevant to brain tissue, CH, and Sylgard to facilitate comparison; (b) comparison of the energy-based metrics between brain tissue and each simulant material (**p<0.01; ***p<0.001).

### S2.3. Normalized stress relaxation

**Table S1.** Normalized stress relaxation at the high stretch rate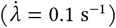 in compression for two representative loading steps (Step 1: *λ* = 0.95; Step 5: *λ* = 0.85) and six time points after the peak stresses. Values are reported as median [Q25–Q75].

| Step<br>[-] | Time after<br>peak [s] | Normalized stress [-] |  |  |  |
| --- | --- | --- | --- | --- | --- |
|  |  | Brain | CH | Sylgard | Ecoflex |
| 1 | 1 | 0.813 [0.793–0.822] | 0.904 [0.892–0.914] | 0.970 [0.960–0.971] | 0.874 [0.868–0.878] |
|  | 10 | 0.584 [0.541–0.612] | 0.774 [0.741–0.794] | 0.920 [0.917–0.932] | 0.759 [0.751–0.764] |
|  | 30 | 0.486 [0.474–0.499] | 0.717 [0.679–0.735] | 0.903 [0.889–0.930] | 0.724 [0.719–0.732] |
|  | 60 | 0.409 [0.391–0.480] | 0.684 [0.638–0.709] | 0.890 [0.884–0.903] | 0.709 [0.700–0.719] |
|  | 300 | 0.280 [0.216–0.337] | 0.622 [0.545–0.636] | 0.887 [0.878–0.897] | 0.690 [0.676–0.698] |
|  | 1200 | 0.218 [0.160–0.285] | 0.595 [0.525–0.629] | 0.886 [0.840–0.921] | 0.677 [0.666–0.685] |
| 5 | 1 | 0.891 [0.872–0.907] | 0.960 [0.952–0.966] | 0.989 [0.988–0.992] | 0.960 [0.960–0.963] |
|  | 10 | 0.747 [0.724–0.761] | 0.894 [0.865–0.910] | 0.978 [0.974–0.979] | 0.927 [0.925–0.931] |
|  | 30 | 0.668 [0.654–0.699] | 0.859 [0.825–0.878] | 0.967 [0.966–0.969] | 0.917 [0.915–0.923] |
|  | 60 | 0.625 [0.607–0.655] | 0.833 [0.797–0.855] | 0.968 [0.960–0.969] | 0.914 [0.911–0.919] |
|  | 300 | 0.514 [0.488–0.545] | 0.771 [0.721–0.797] | 0.965 [0.951–0.973] | 0.905 [0.902–0.912] |
|  | 1200 | 0.417 [0.387–0.456] | 0.733 [0.652–0.769] | 0.942 [0.932–0.970] | 0.901 [0.896–0.905] |

**Table S2.** Normalized stress relaxation at the low stretch rate 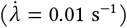 in compression for two representative loading steps (Step 1: *λ* = 0.95; Step 5: *λ* = 0.85) and six time points after the peak stresses. Values are reported as median [Q25–Q75].

| Step [-] | Time after peak [s] | Normalized stress [-] |  |  |  |
| --- | --- | --- | --- | --- | --- |
|  |  | Brain | CH | Sylgard | Ecoflex |
| 1 | 1 | 0.898 [0.884–0.911] | 0.955 [0.947–0.961] | 0.980 [0.974–0.985] | 0.960 [0.957–0.961] |
|  | 10 | 0.700 [0.678–0.744] | 0.847 [0.834–0.856] | 0.957 [0.951–0.963] | 0.883 [0.878–0.887] |
|  | 30 | 0.565 [0.550–0.593] | 0.785 [0.737–0.810] | 0.926 [0.910–0.934] | 0.850 [0.846–0.858] |
|  | 60 | 0.517 [0.462–0.553] | 0.760 [0.708–0.770] | 0.924 [0.919–0.938] | 0.831 [0.824–0.839] |
|  | 300 | 0.361 [0.320–0.377] | 0.662 [0.635–0.685] | 0.937 [0.931–0.953] | 0.800 [0.786–0.809] |
|  | 1200 | 0.214 [0.158–0.337] | 0.654 [0.589–0.720] | 0.944 [0.927–0.955] | 0.791 [0.774–0.806] |
| 5 | 1 | 0.926 [0.919–0.936] | 0.972 [0.965–0.977] | 0.991 [0.988–0.992] | 0.983 [0.982–0.983] |
|  | 10 | 0.800 [0.798–0.803] | 0.908 [0.897–0.925] | 0.982 [0.978–0.982] | 0.960 [0.958–0.960] |
|  | 30 | 0.726 [0.724–0.737] | 0.871 [0.848–0.912] | 0.974 [0.968–0.975] | 0.951 [0.949–0.952] |
|  | 60 | 0.684 [0.673–0.695] | 0.846 [0.823–0.889] | 0.971 [0.969–0.971] | 0.946 [0.944–0.948] |
|  | 300 | 0.574 [0.550–0.587] | 0.789 [0.754–0.817] | 0.965 [0.959–0.967] | 0.937 [0.935–0.939] |
|  | 1200 | 0.477 [0.448–0.496] | 0.764 [0.724–0.800] | 0.959 [0.951–0.966] | 0.929 [0.925–0.933] |

**Figure S11.**
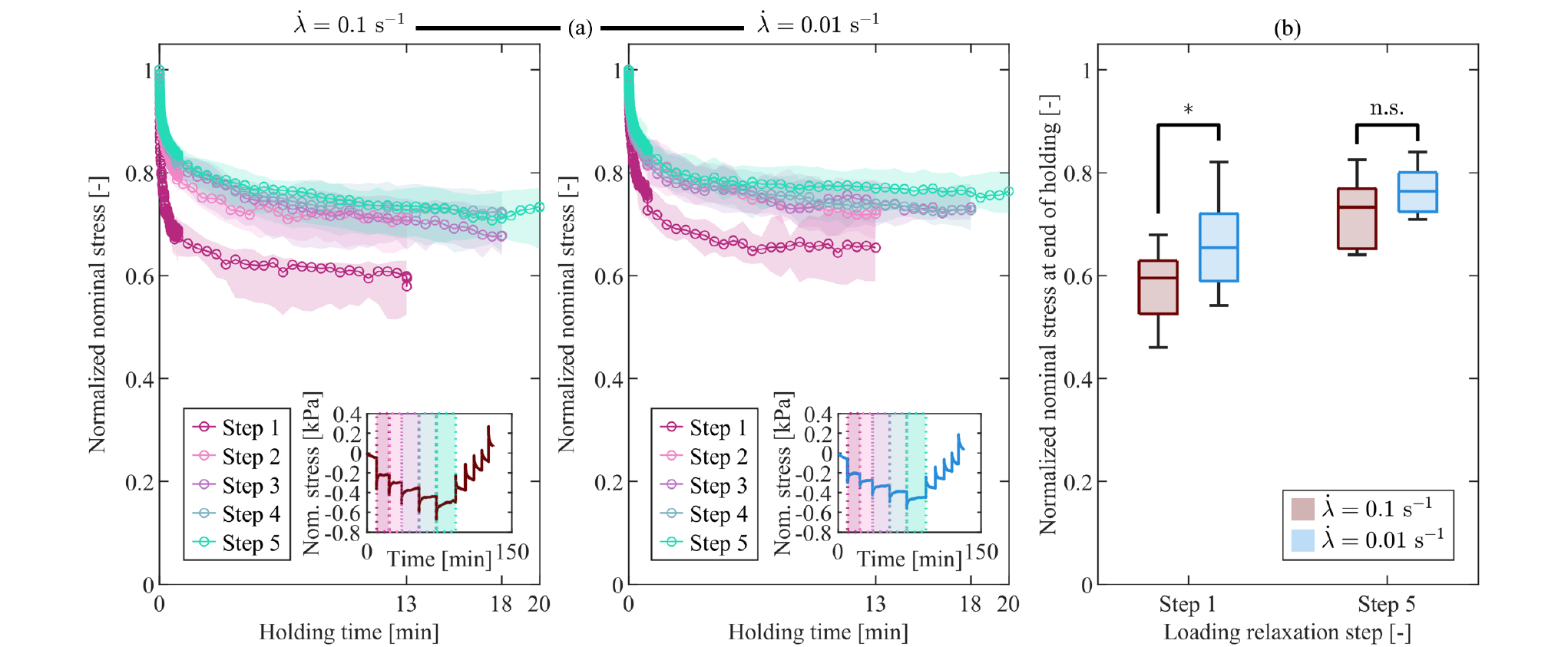
CH normalized stress relaxation response in compression at high and low stretch rates. (a) Median normalized nominal stress during the holding phases for all loading steps and both stretch rates. Shaded regions indicate the interquartile range (IQR). The insets show the corresponding median nominal stress–time curves and identify the individual loading steps; (b) comparison of normalized stress between the high (red) and low (blue) stretch rates for two representative loading steps (Step 1: *λ* = 0.95; Step 5: *λ* = 0.85). Significant difference was found at Step 1 (*p<0.05) but not at Step 5.

**Figure S12.**
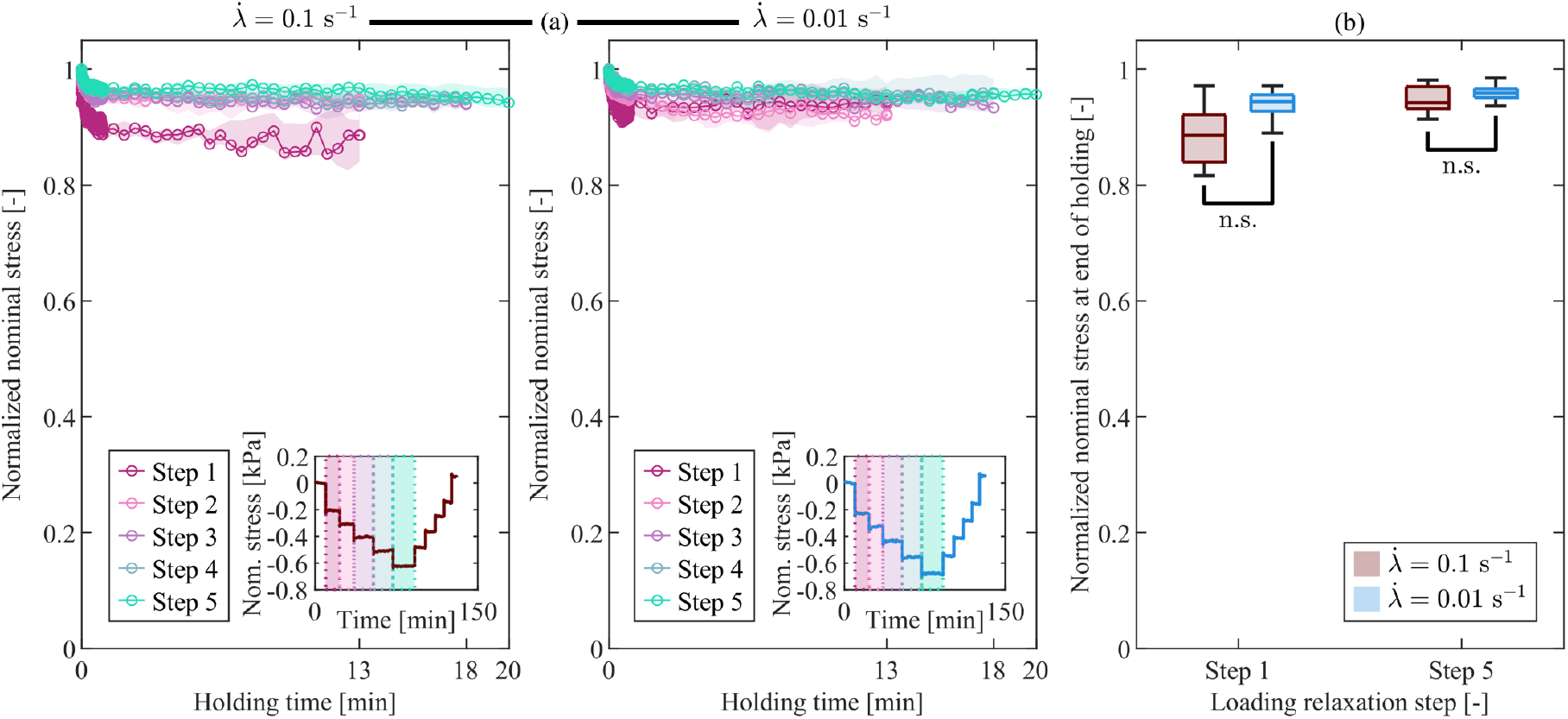
Sylgard normalized stress relaxation response in compression at high and low stretch rates. (a) Median normalized nominal stress during the holding phases for all loading steps and both stretch rates. Shaded regions indicate the interquartile range (IQR). The insets show the corresponding median nominal stress–time curves and identify the individual loading steps; (b) comparison of normalized stress between the high (red) and low (blue) stretch rates for two representative loading steps (Step 1: *λ* = 0.95; Step 5: *λ* = 0.85). No significant differences were found.

**Figure S13.**
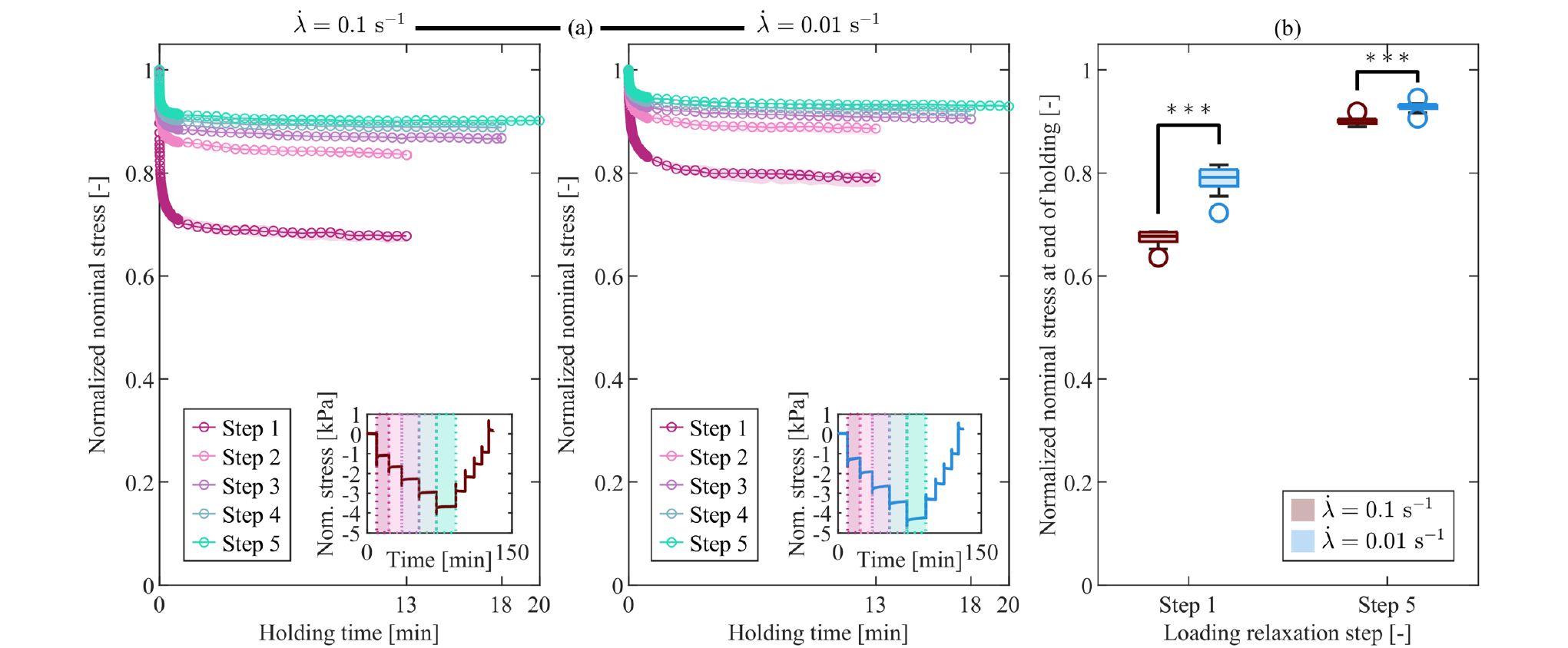
Ecoflex normalized stress relaxation response in compression at high and low stretch rates. (a) Median normalized nominal stress during the holding phases for all loading steps and both stretch rates. Shaded regions indicate the interquartile range (IQR). The insets show the corresponding median nominal stress–time curves and identify the individual loading steps; (b) comparison of normalized stress between the high (red) and low (blue) stretch rates for two representative loading steps (Step 1: *λ* = 0.95; Step 5: *λ* = 0.85). Significant differences were found at both loading steps (***p<0.001).

As mentioned in the main manuscript, the simulant materials exhibited different stretch-rate dependence of normalized stress relaxation compared to brain tissue (Figs. S11–S13). In brain tissue, statistically significant differences in normalized nominal stress at the end of the holding phase between stretch rates are observed only at the higher deformation levels, corresponding to Steps 4 and 5 (p = 0.0376 and p = 0.0452, respectively). In contrast, CH showed a statistically significant dependence on the stretch rate at the lowest deformation level (Step 1: p = 0.0312), while no consistent trend was observed across the remaining steps. Similarly, Sylgard exhibited a rate dependence that is limited to a low deformation level (Step 2: p = 0.0317), without a clear pattern throughout the loading history. Ecoflex, on the other hand, displayed a pronounced rate dependence at all deformation levels, with statistically significant differences between stretch rates observed at every step (Step 1: p = 1.8267 · 10^−4^; Step 2: p = 3.2984 · 10^−4^; Step 3: p = 5.8284 · 10^−4^; Step 4: p = 7.6854 · 10^−4^; Step 5: p = 3.2984 · 10^−4^).

**Figure S14.**
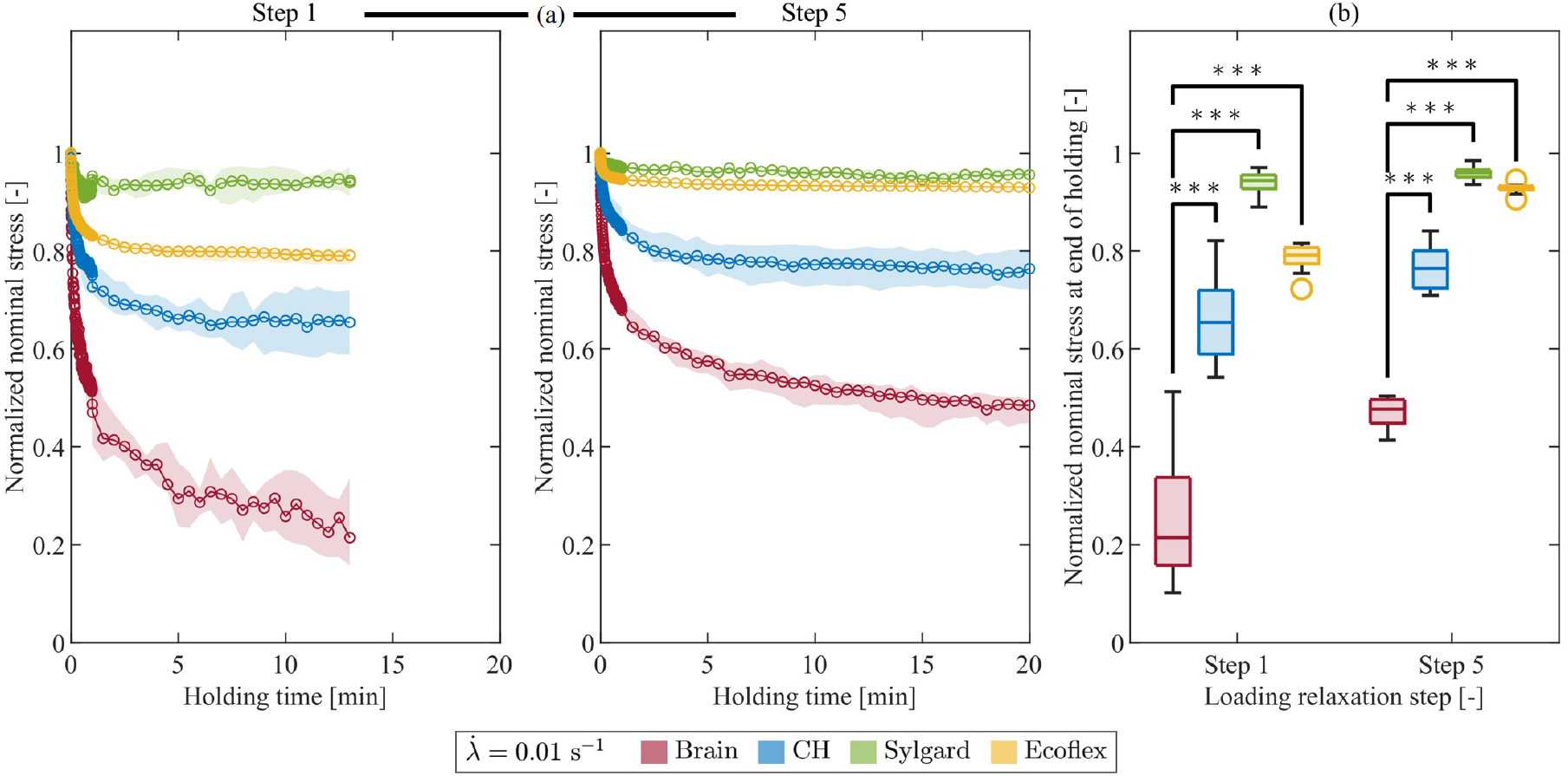
Comparison of normalized stress relaxation in compression across all materials at the low stretch rate for two representative loading steps (Step 1: *λ* = 0.95; Step 5: *λ* = 0.85). (a) Median normalized nominal stress during the holding phase with IQR (shaded areas); (b) comparison of normalized nominal stress at the end of the holding phase between brain tissue and each simulant material (***p<0.001).

## S3. Tension experiments: additional figures and tables

### S3.1. Equilibrium response

**Figure S15.**
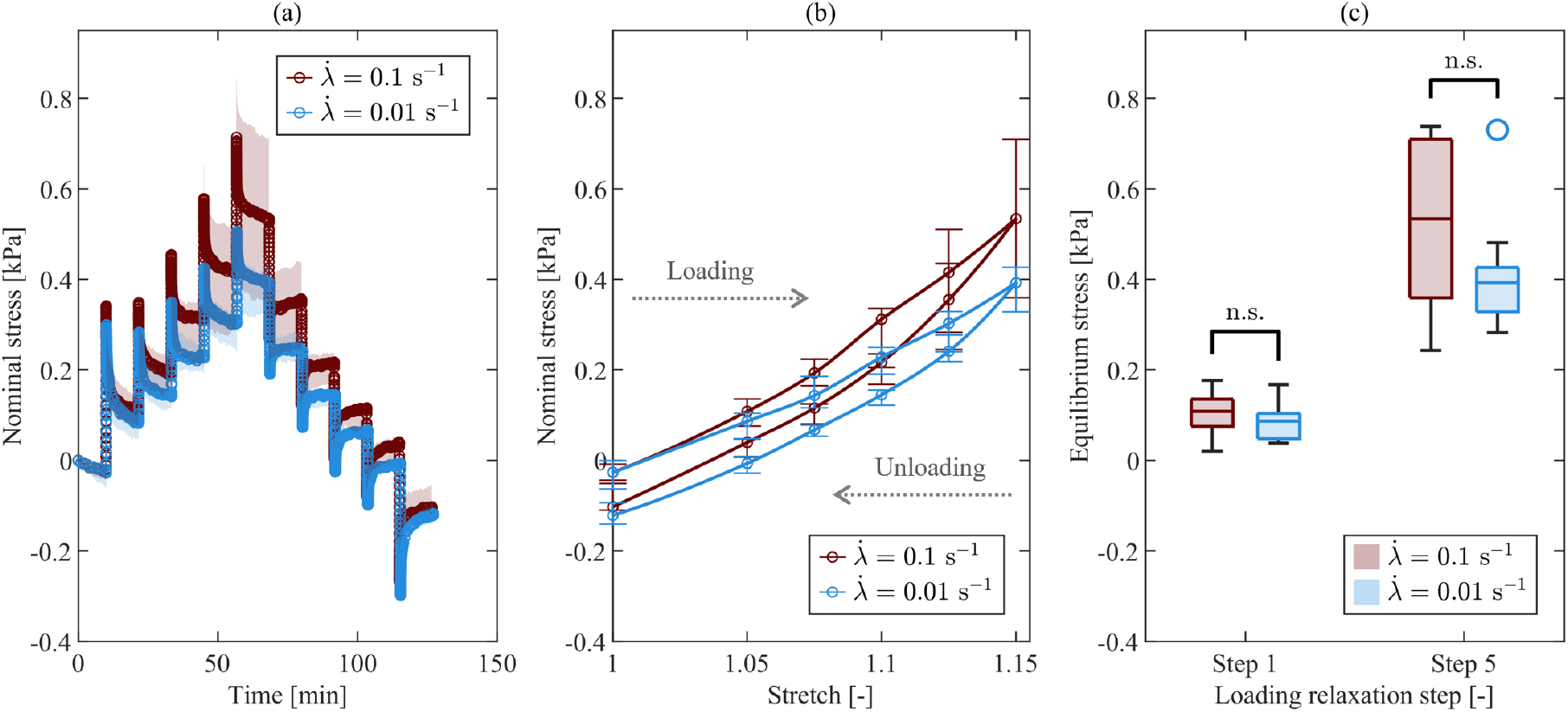
CH equilibrium response in tension at high (red) and low (blue) stretch rates. (a) Median nominal stress–time with IQR (shaded areas); (b) equilibrium stress–stretch relation with IQR (bars); (c) comparison of equilibrium stress at two representative loading steps (Step 1: *λ* = 1.05; Step 5: *λ* = 1.15) between stretch rates, showing no significant differences.

**Figure S16.**
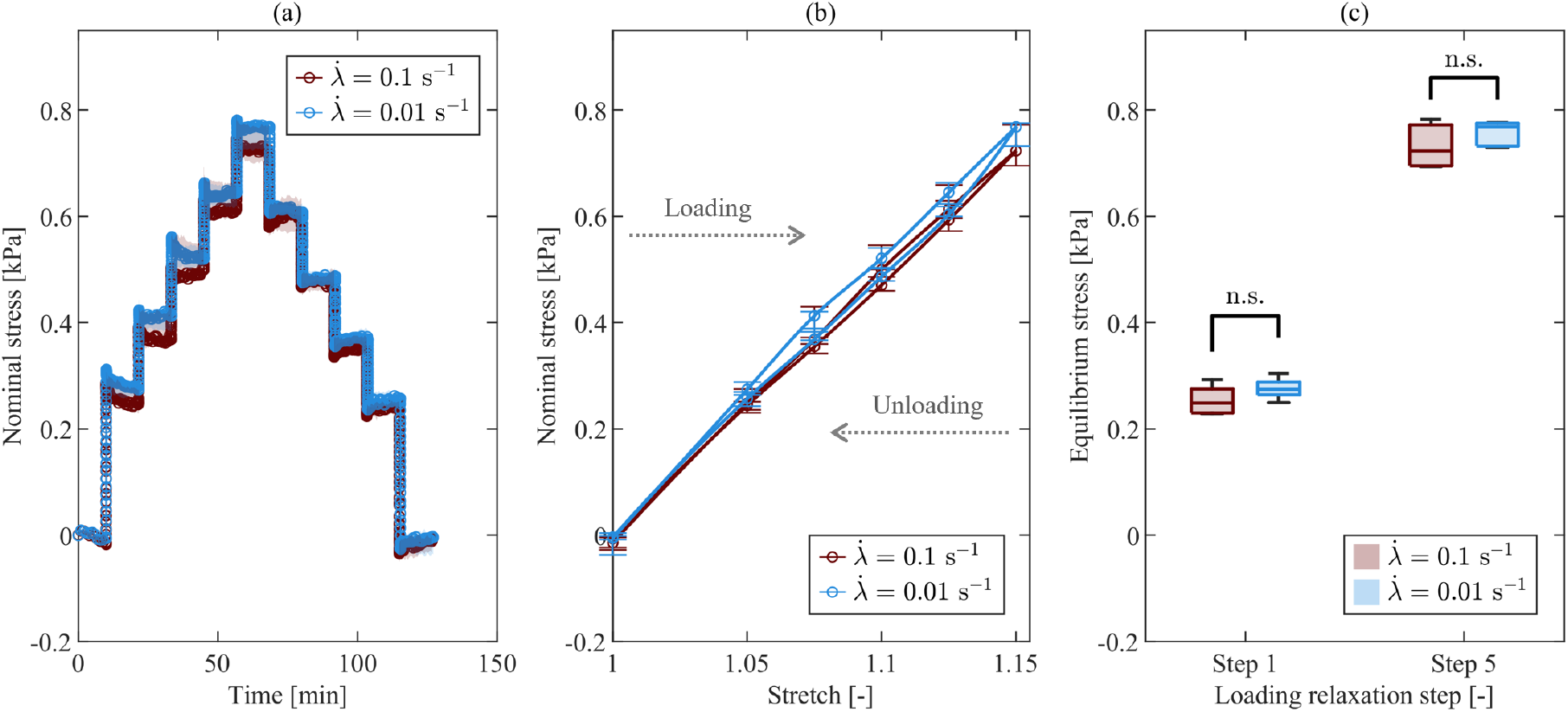
Sylgard equilibrium response in tension at high (red) and low (blue) stretch rates. (a) Median nominal stress–time with IQR (shaded areas); (b) equilibrium stress–stretch relation with IQR (bars); (c) comparison of equilibrium stress at two representative loading steps (Step 1: *λ* = 1.05; Step 5: *λ* = 1.15) between stretch rates, showing no significant differences.

**Figure S17.**
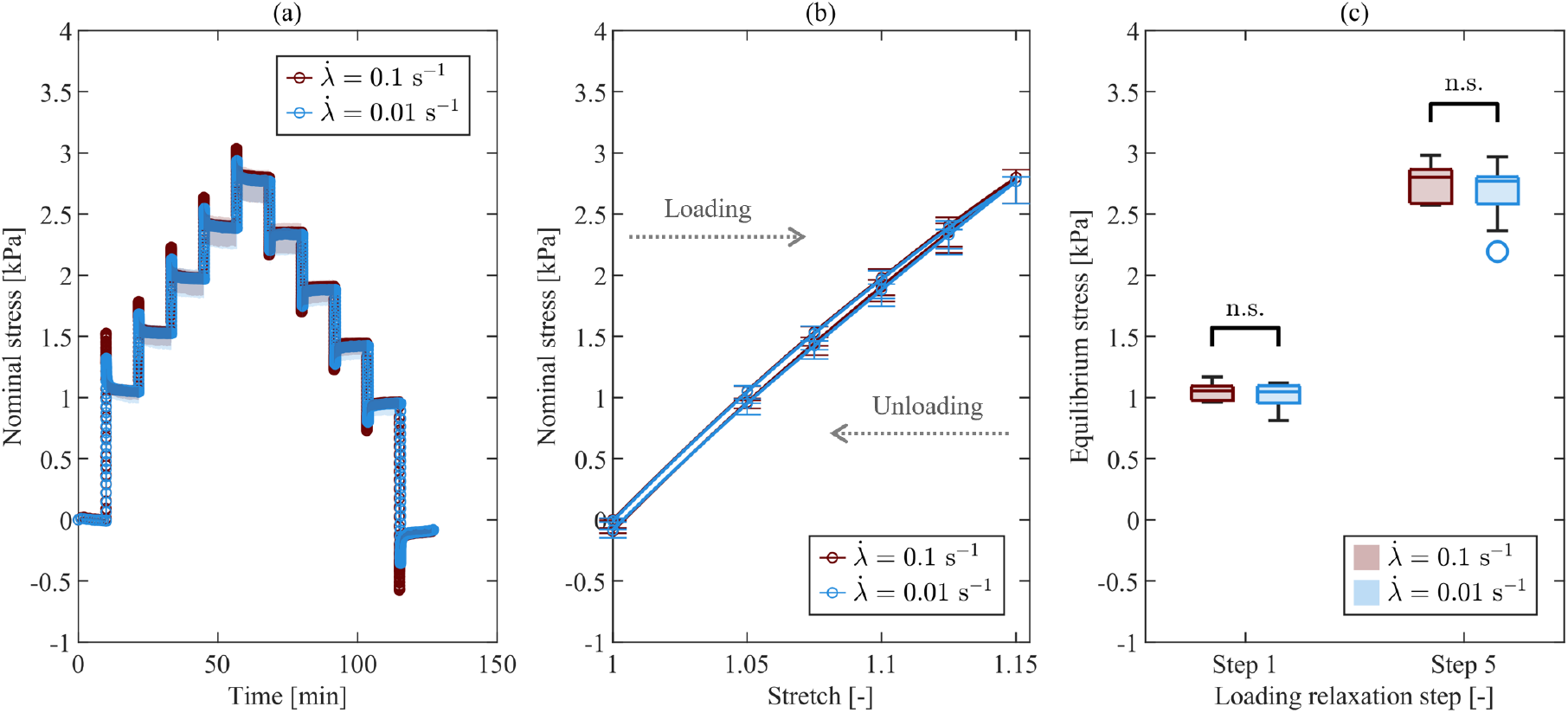
Ecoflex equilibrium response in tension at high (red) and low (blue) stretch rates. (a) Median nominal stress–time with IQR (shaded areas); (b) equilibrium stress–stretch relation with IQR (bars); (c) comparison of equilibrium stress at two representative loading steps (Step 1: *λ* = 1.05; Step 5: *λ* = 1.15) between stretch rates, showing no significant differences.

**Figure S18.**
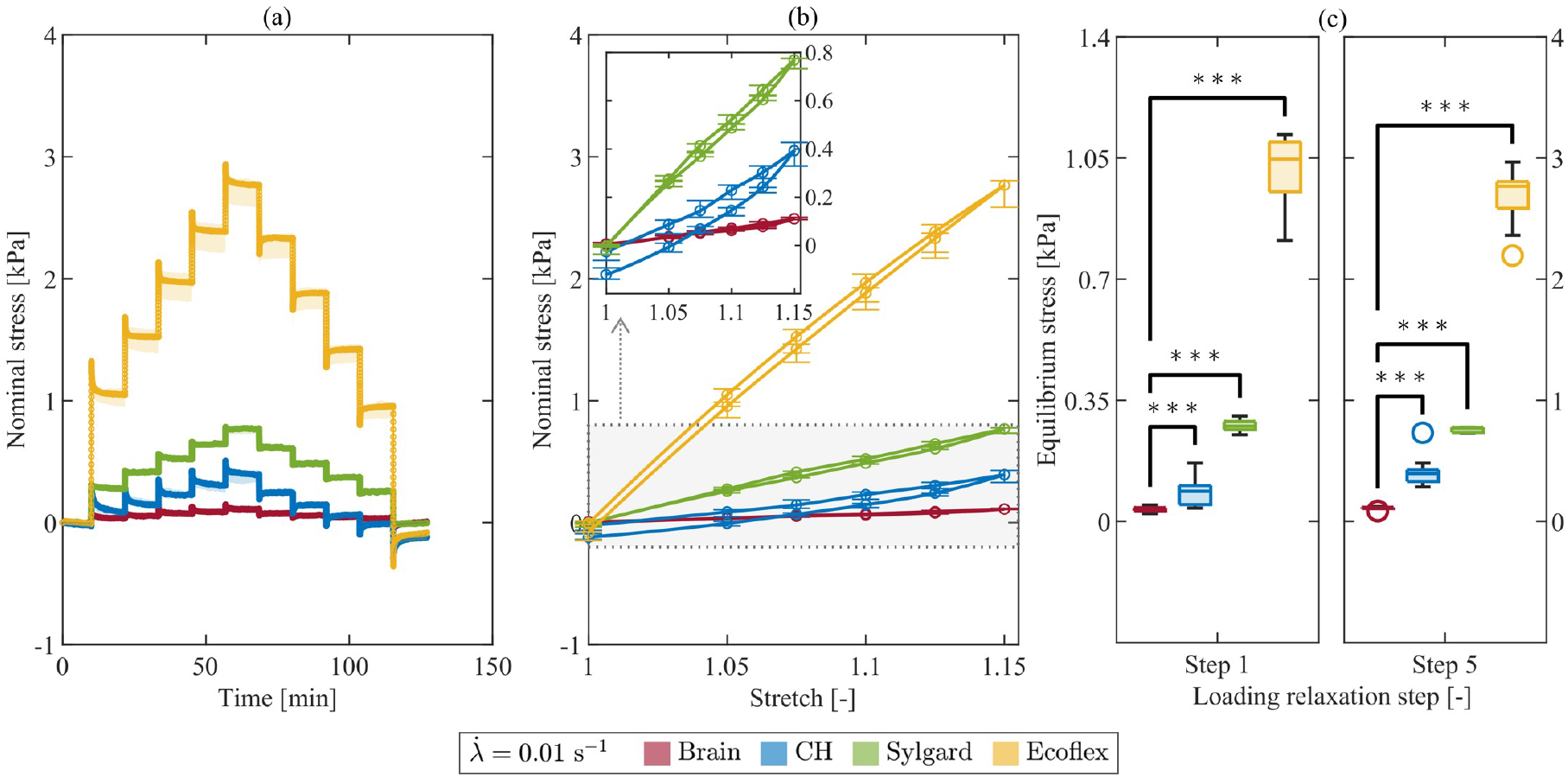
Comparison of equilibrium response in tension across all materials at the low stretch rate. (a) Median nominal stress–time with IQR (shaded areas); (b) equilibrium stress–stretch relation with IQR (bars). The inset enlarges the stress range relevant to brain tissue, CH, and Sylgard to facilitate comparison; (c) comparison of equilibrium stress at two representative loading steps (Step 1: *λ* = 1.05; Step 5: *λ* = 1.15) between brain tissue and each simulant material (***p<0.001).

### S3.2. Inelastic effects

**Figure S19.**
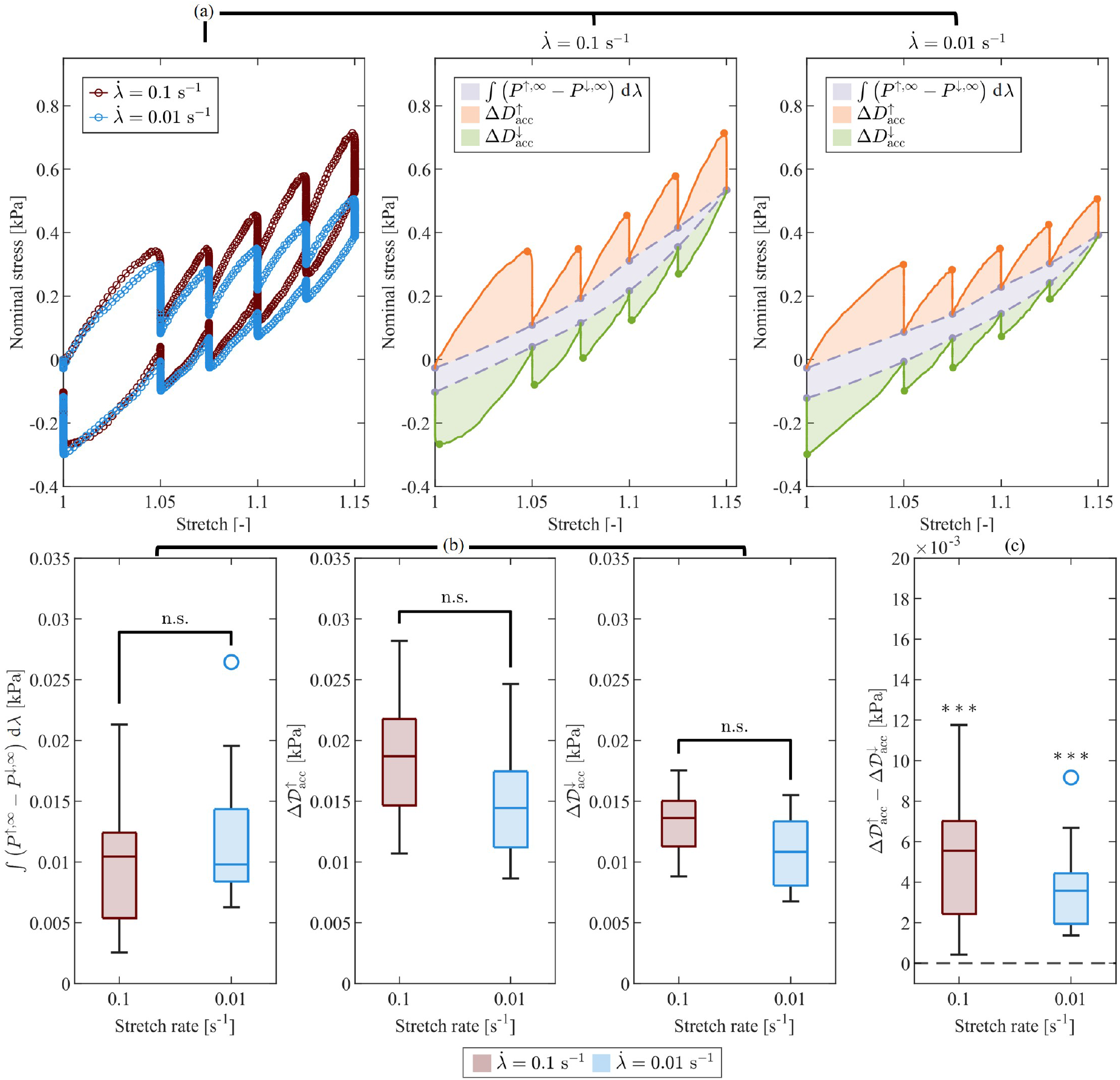
CH inelastic effects in tension at high (red) and low (blue) stretch rates. (a) Median nominal stress–stretch response with shaded areas representing the stress work associated with the equilibrium relation (gray), and accumulated dissipation during loading (orange) and unloading (green), at the two stretch rates; (b) comparison of the energy-based metrics between the stretch rates (MWU test), showing no evidence of significant rate dependence; (c) comparison between loading and unloading accumulated dissipation (one-tailed WSR test), indicating significantly higher dissipation during loading at both stretch rates (***p<0.001).

**Figure S20.**
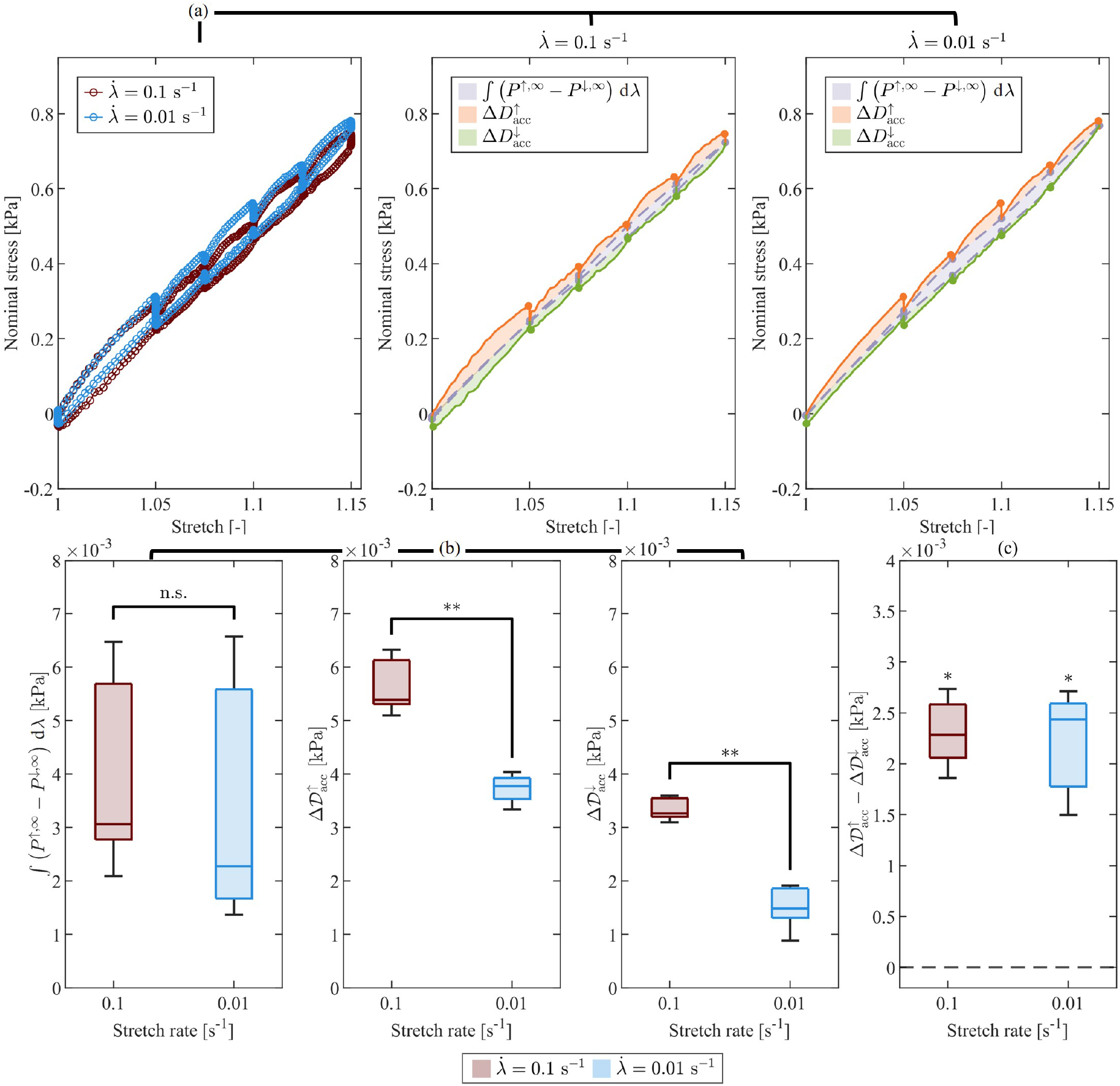
Sylgard inelastic effects in tension at high (red) and low (blue) stretch rates. (a) Median nominal stress–stretch response with shaded areas representing the stress work associated with the equilibrium relation (gray), and accumulated dissipation during loading (orange) and unloading (green), at the two stretch rates; (b) comparison of the energy-based metrics between the stretch rates (MWU test), showing rate-dependent accumulated dissipation (**p<0.01) but no evidence of significant rate dependence in the equilibrium stress work; (c) comparison between loading and unloading accumulated dissipation (one-tailed WSR test), indicating significantly higher dissipation during loading at both stretch rates (*p<0.05).

**Figure S21.**
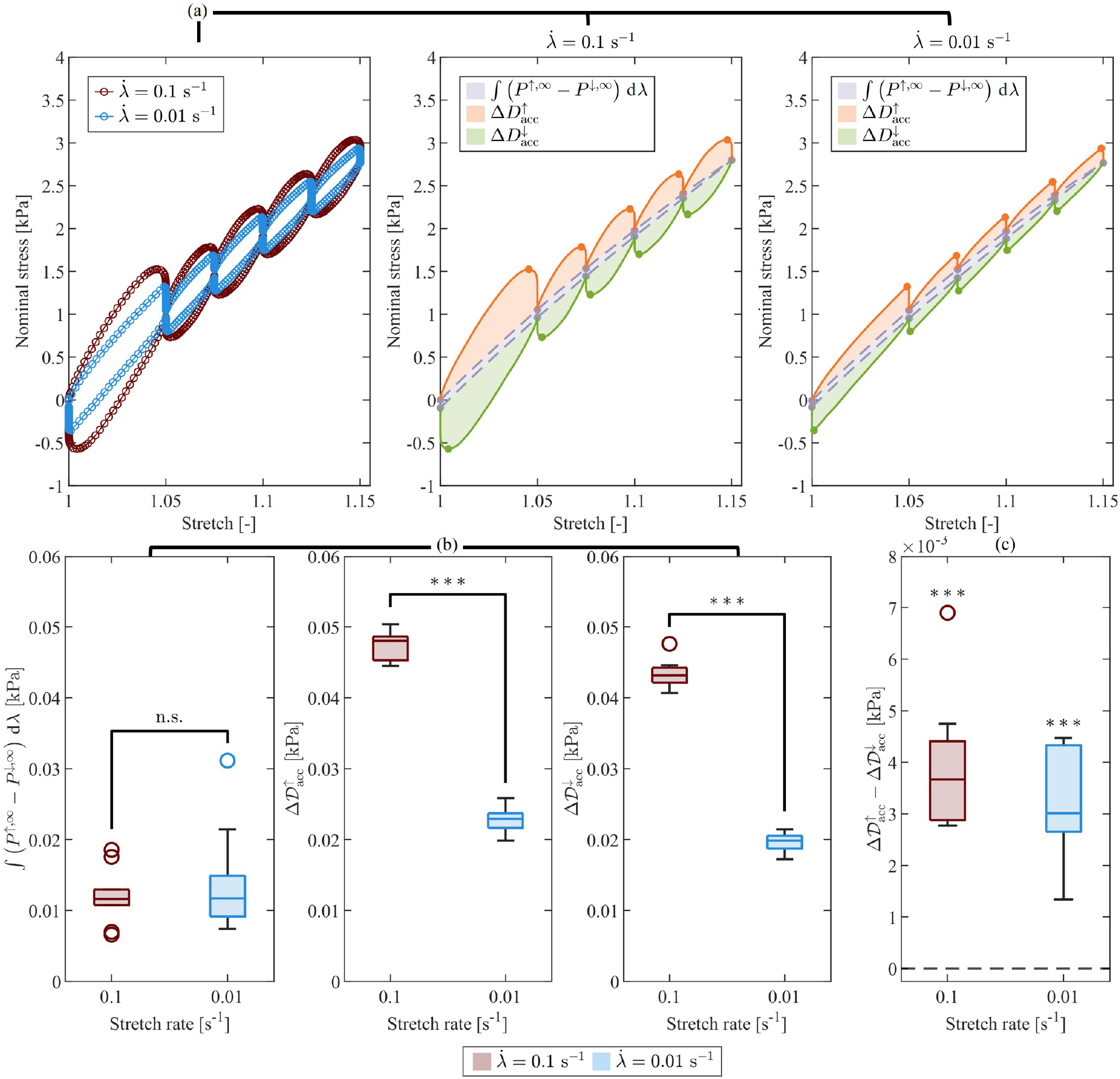
Ecoflex inelastic effects in tension at high (red) and low (blue) stretch rates. (a) Median nominal stress–stretch response with shaded areas representing the stress work associated with the equilibrium relation (gray), and accumulated dissipation during loading (orange) and unloading (green), at the two stretch rates; (b) comparison of the energy-based metrics between the stretch rates (MWU test), showing rate-dependent accumulated dissipation (***p<0.001) but no evidence of significant rate dependence in the equilibrium stress work; (c) comparison between loading and unloading accumulated dissipation (one-tailed WSR test), indicating significantly higher dissipation during loading at both stretch rates (***p<0.001).

**Figure S22.**
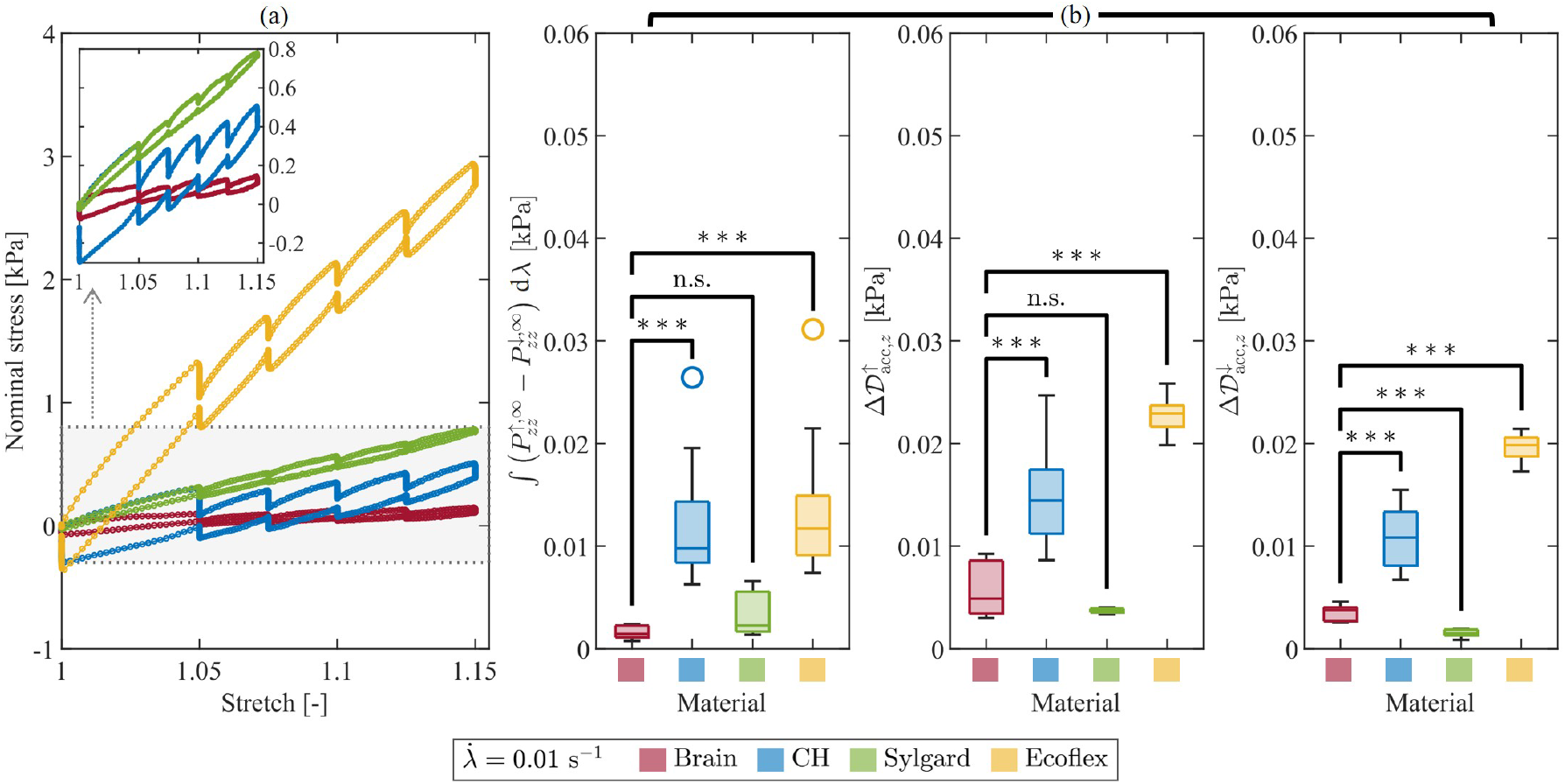
Comparison of inelastic effects in tension across all materials at the low stretch rate. (a) Median nominal stress–stretch response. The inset enlarges the stress range relevant to brain tissue, CH, and Sylgard to facilitate comparison; (b) comparison of the energy-based metrics between brain tissue and each simulant material (***p<0.001).

### S3.3. Normalized stress relaxation

**Table S3.** Normalized stress relaxation at the high stretch rate 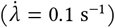 in tension for two representative loading steps (Step 1: *λ* = 1.05; Step 5: *λ* = 1.15) and six time points after the peak stresses. Values are reported as median [Q25–Q75].

| Step<br>[-] | Time after<br>peak [s] | Normalized stress [-] |  |  |  |
| --- | --- | --- | --- | --- | --- |
|  |  | Brain | CH | Sylgard | Ecoflex |
| 1 | 1 | 0.767 [0.751–0.793] | 0.867 [0.845–0.886] | 0.979 [0.973–0.982] | 0.875 [0.873–0.878] |
|  | 10 | 0.539 [0.488–0.575] | 0.663 [0.641–0.706] | 0.940 [0.929–0.945] | 0.763 [0.759–0.765] |
|  | 30 | 0.462 [0.433–0.509] | 0.572 [0.550–0.653] | 0.915 [0.899–0.931] | 0.732 [0.729–0.736] |
|  | 60 | 0.406 [0.369–0.498] | 0.517 [0.484–0.584] | 0.922 [0.913–0.939] | 0.717 [0.714–0.721] |
|  | 300 | 0.282 [0.247–0.366] | 0.384 [0.327–0.445] | 0.905 [0.849–0.913] | 0.693 [0.685–0.701] |
|  | 700 | 0.285 [0.231–0.382] | 0.312 [0.246–0.370] | 0.866 [0.838–0.889] | 0.682 [0.677–0.695] |
| 5 | 1 | 0.876 [0.867–0.889] | 0.945 [0.943–0.947] | 0.989 [0.988–0.993] | 0.970 [0.969–0.971] |
|  | 10 | 0.747 [0.741–0.767] | 0.864 [0.852–0.877] | 0.976 [0.969–0.979] | 0.944 [0.942–0.945] |
|  | 30 | 0.707 [0.693–0.744] | 0.837 [0.813–0.851] | 0.969 [0.965–0.970] | 0.937 [0.934–0.938] |
|  | 60 | 0.674 [0.657–0.713] | 0.815 [0.796–0.836] | 0.966 [0.962–0.967] | 0.933 [0.931–0.934] |
|  | 300 | 0.607 [0.579–0.658] | 0.769 [0.745–0.780] | 0.966 [0.954–0.971] | 0.926 [0.921–0.928] |
|  | 700 | 0.565 [0.552–0.609] | 0.767 [0.719–0.778] | 0.966 [0.952–0.969] | 0.921 [0.918–0.925] |

**Table S4.**
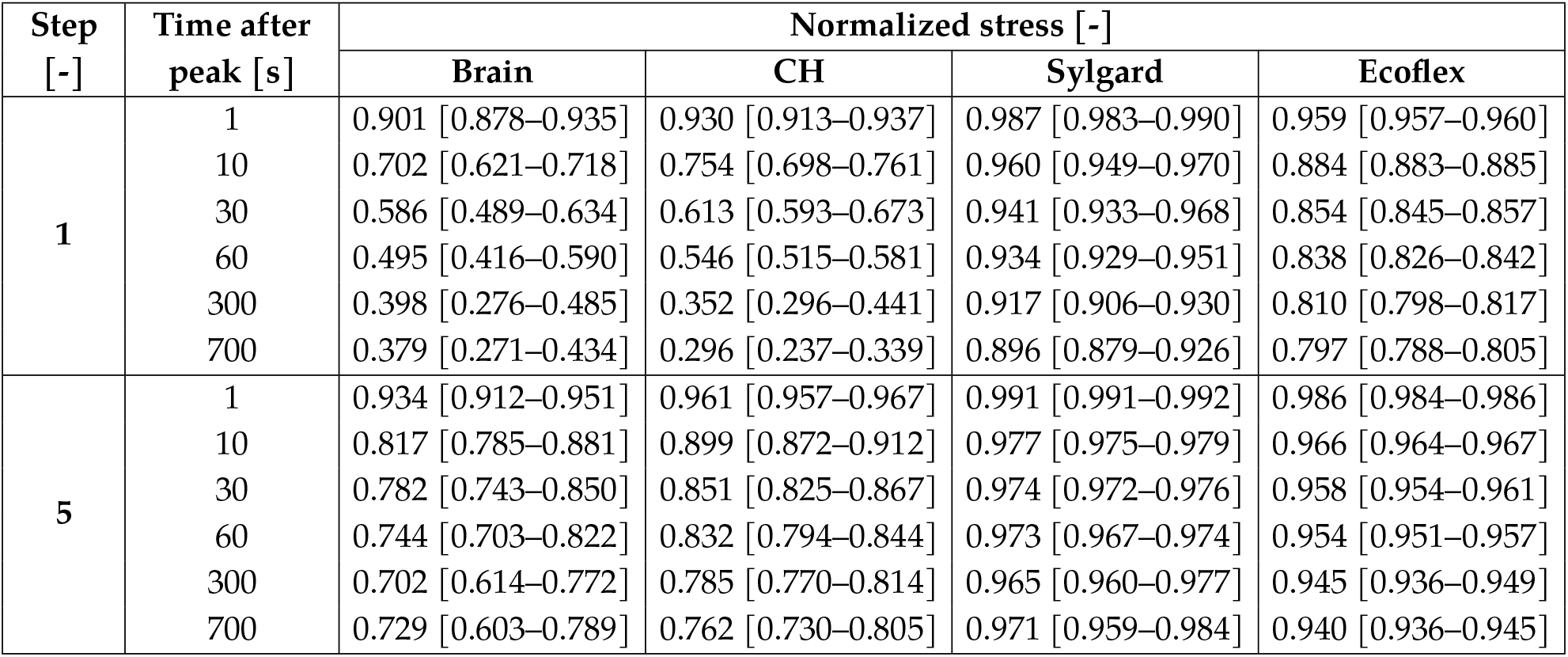
Normalized stress relaxation at the low stretch rate 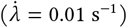 in tension for two representative loading steps (Step 1: *λ* = 1.05; Step 5: *λ* = 1.15) and six time points after the peak stresses. Values are reported as median [Q25–Q75].

| Step<br>[-] | Time after<br>peak [s] | Normalized stress [-] |  |  |  |
| --- | --- | --- | --- | --- | --- |
|  |  | Brain | CH | Sylgard | Ecoflex |
| 1 | 1 | 0.901 [0.878–0.935] | 0.930 [0.913–0.937] | 0.987 [0.983–0.990] | 0.959 [0.957–0.960] |
|  | 10 | 0.702 [0.621–0.718] | 0.754 [0.698–0.761] | 0.960 [0.949–0.970] | 0.884 [0.883–0.885] |
|  | 30 | 0.586 [0.489–0.634] | 0.613 [0.593–0.673] | 0.941 [0.933–0.968] | 0.854 [0.845–0.857] |
|  | 60 | 0.495 [0.416–0.590] | 0.546 [0.515–0.581] | 0.934 [0.929–0.951] | 0.838 [0.826–0.842] |
|  | 300 | 0.398 [0.276–0.485] | 0.352 [0.296–0.441] | 0.917 [0.906–0.930] | 0.810 [0.798–0.817] |
|  | 700 | 0.379 [0.271–0.434] | 0.296 [0.237–0.339] | 0.896 [0.879–0.926] | 0.797 [0.788–0.805] |
| 5 | 1 | 0.934 [0.912–0.951] | 0.961 [0.957–0.967] | 0.991 [0.991–0.992] | 0.986 [0.984–0.986] |
|  | 10 | 0.817 [0.785–0.881] | 0.899 [0.872–0.912] | 0.977 [0.975–0.979] | 0.966 [0.964–0.967] |
|  | 30 | 0.782 [0.743–0.850] | 0.851 [0.825–0.867] | 0.974 [0.972–0.976] | 0.958 [0.954–0.961] |
|  | 60 | 0.744 [0.703–0.822] | 0.832 [0.794–0.844] | 0.973 [0.967–0.974] | 0.954 [0.951–0.957] |
|  | 300 | 0.702 [0.614–0.772] | 0.785 [0.770–0.814] | 0.965 [0.960–0.977] | 0.945 [0.936–0.949] |
|  | 700 | 0.729 [0.603–0.789] | 0.762 [0.730–0.805] | 0.971 [0.959–0.984] | 0.940 [0.936–0.945] |

**Figure S23.**
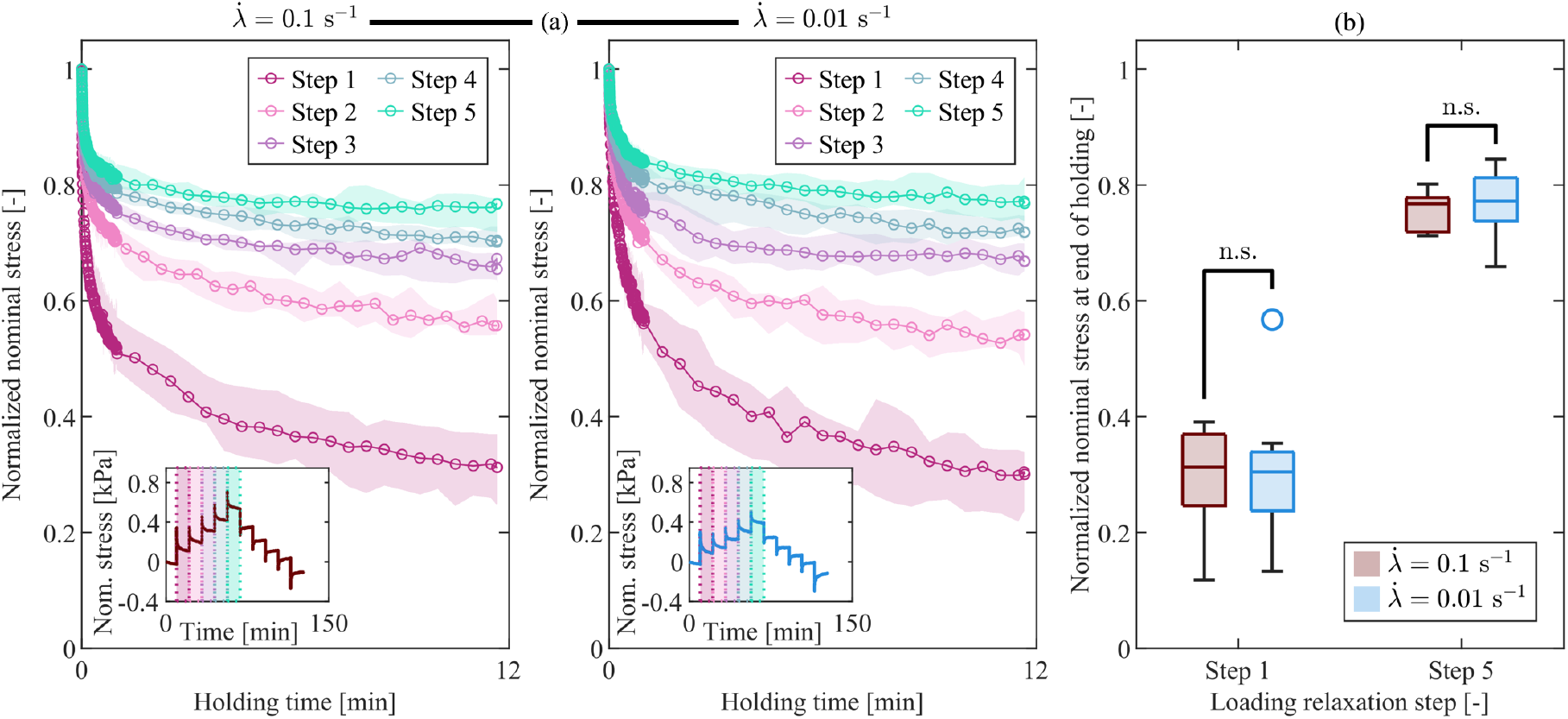
CH normalized stress relaxation response in tension at high andlow stretch rates. (a) Median normalized nominal stress during the holding phases for all loading steps and both stretch rates. Shaded regions indicate the interquartile range (IQR). The insets show the corresponding median nominal stress–time curves and identify the individual loading steps; (b) comparison of normalized stress between the high (red) and low (blue) stretch rates for two representative loading steps (Step 1: *λ* = 1.05; Step 5: *λ* = 1.15). No significant differences were observed.

**Figure S24.**
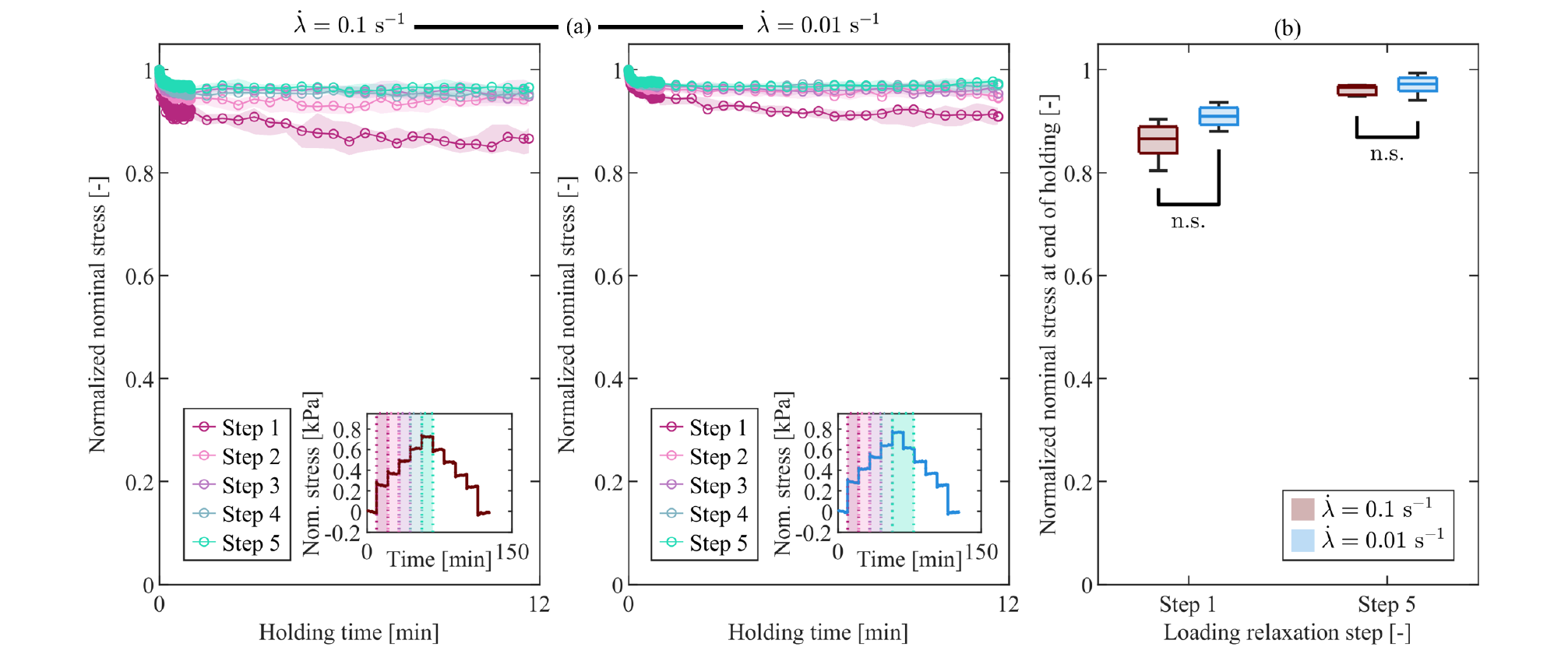
Sylgard normalized stress relaxation response in tension at high and low stretch rates. (a) Median normalized nominal stress during the holding phases for all loading steps and both stretch rates. Shaded regions indicate the interquartile range (IQR). The insets show the corresponding median nominal stress–time curves and identify the individual loading steps; (b) comparison of normalized stress between the high (red) and low (blue) stretch rates for two representative loading steps (Step 1: *λ* = 1.05; Step 5: *λ* = 1.15). No significant differences were observed.

**Figure S25.**
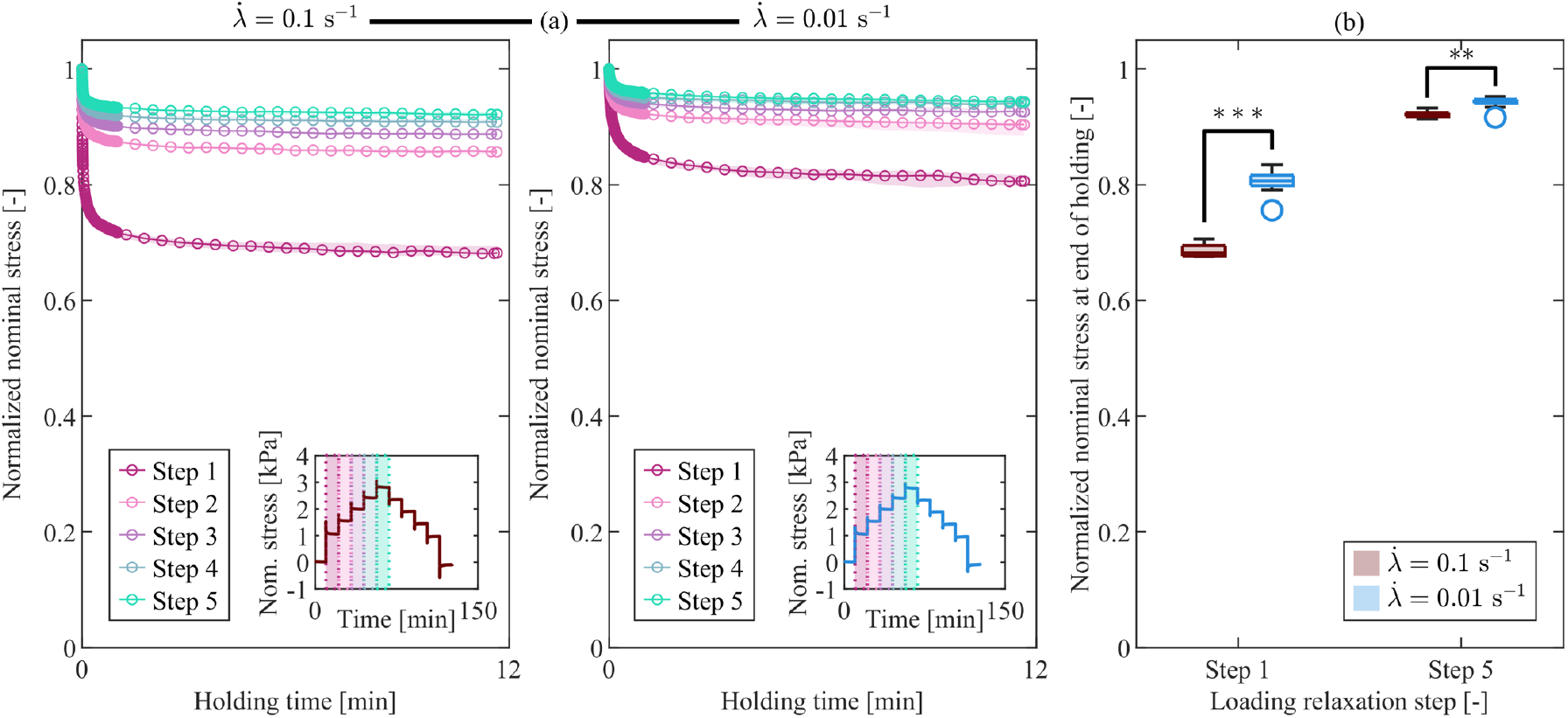
Ecoflex normalized stress relaxation response in tension at high and low stretch rates. (a) Median normalized nominal stress during the holding phases for all loading steps and both stretch rates. Shaded regions indicate the interquartile range (IQR). The insets show the corresponding median nominal stress–time curves and identify the individual loading steps; (b) comparison of normalized stress between the high (red) and low (blue) stretch rates for two representative loading steps (Step 1: *λ* = 1.05; Step 5: *λ* = 1.15). Significant differences were found in both loading steps (**p<0.01; ***p<0.001).

As mentioned in the main manuscript, the rate-dependent normalized stress relaxation of CH and Sylgard in tension diverged consistently from that of brain tissue (Figs. S23–S25). Brain tissue revealed statistically significant differences between the stretch rates at all investigated deformation levels (Step 1: p = 0.0312; Step 2: p = 0.0091; Step 3: p = 0.0028; Step 4: p = 0.0046; Step 5: p = 0.0008), indicating that the rate influences the normalized stress relaxation response throughout the entire loading history. In contrast, CH and Sylgard showed no evidence of statistically significant differences between stretch rates at any investigated deformation level (Steps 1–5, *p* > 0.05). Ecoflex, on the other hand, displayed a pronounced rate dependence, with statistically significant differences between high and low stretch rates observed at all deformation levels (Step 1: p=2.0 · 10^−4^; Step 2: p=2.0 · 10^−4^; Step 3: p=0.0022; Step 4: p=6.0 · 10^−4^; Step 5: p=0.0017).

**Figure S26.**
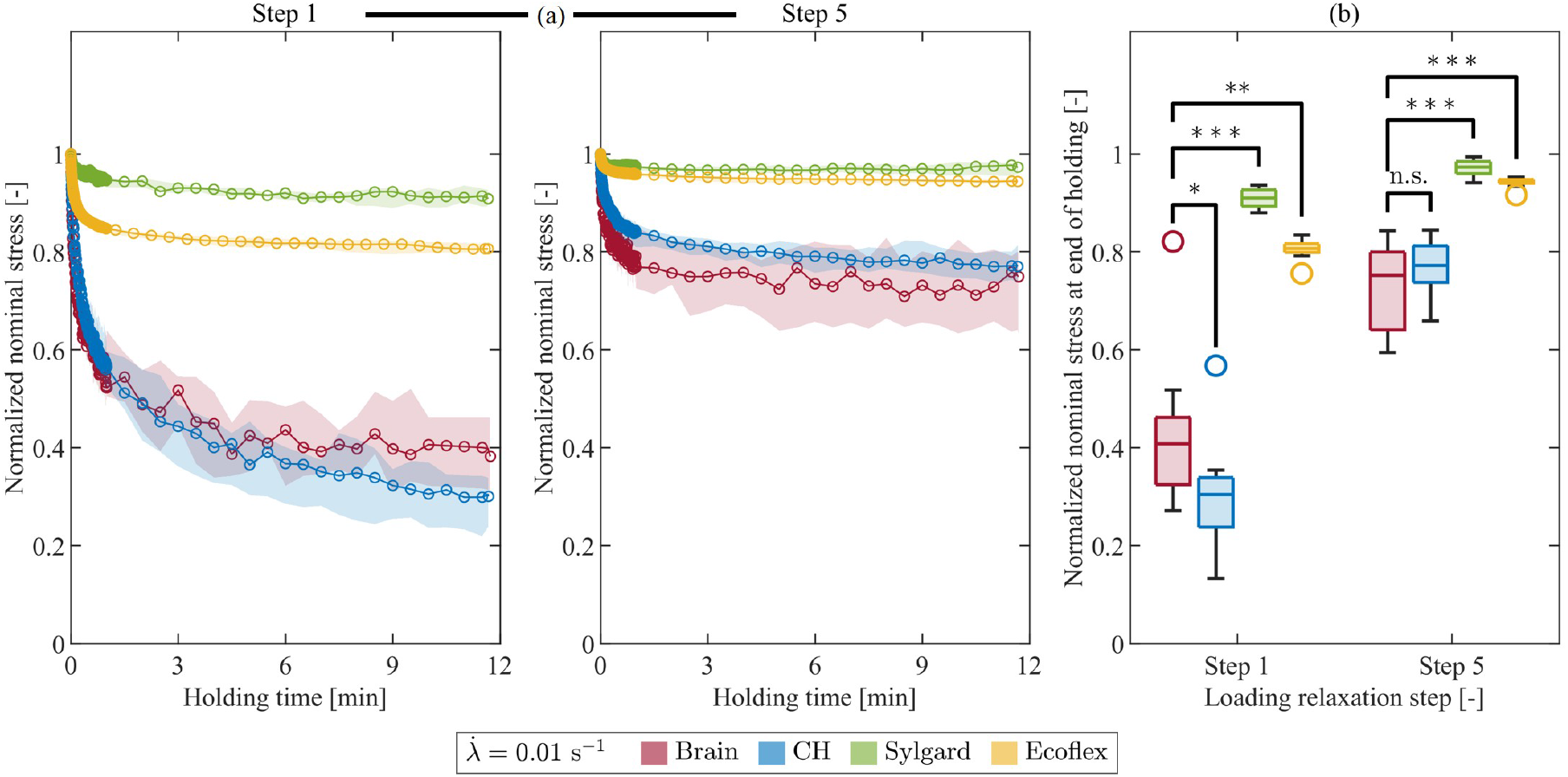
Comparison of normalized stress relaxation in tension across all materials at the low stretch rate for two representative loading steps (Step 1: *λ* = 1.05; Step 5: *λ* = 1.15). (a) Median normalized nominal stress during the holding phase with IQR (shaded areas); (b) comparison of normalized nominal stress at the end of the holding phase between brain tissue and each simulant material (*p<0.05; **p<0.01; ***p<0.001).

## Footnotes

1 Kinematic quantities in a biphasic mixture are customarily referred to the solid constituent. Accordingly, the motion, deformation gradient, and Jacobian introduced here are referred to the solid phase. A superposed dot denotes the material time derivative following the solid motion. Constituent-specific indices, which are common in mixture theories, are omitted in order to improve readability.

2 The dissipation inequality at the mixture level is formulated at a material point in the current configuration, because, in general, the different constituents do not share a common reference configuration.

## References

[1] A. Leibinger, A. E. Forte, Z. Tan, M. J. Oldfield, F. Beyrau, D. Dini, F. Rodriguez y Baena, Soft tissue phantoms for realistic needle insertion: A comparative study, Annals Biomed. Eng. 44 (2016) 2442–2452.

[2] A. E. Forte, S. Galvan, F. Manieri, F. Rodriguez y Baena, D. Dini, A composite hydrogel for brain tissue phantoms, Mater. Des. 112 (2016) 227–238.

[3] A. A. Alshareef, J. S. Giudice, D. F. Shedd, Y.-C. Lu, C. L. Johnson, D. L. Pham, M. B. Panzer, Multimodal characterization of intracranial biomechanics in a 3D biofidelic head surrogate, J. Mech. Behav. Biomed. Mater. 173 (2026) 107250.

[4] J. F. Cuello, A. Saenz, J. M. Liñares, P. Martinez, C. Ruiz, R. Argañaraz, M. M. Bailez, B. Mantese, Low-cost stereotactic brain biopsy simulation model, World Neurosurg. 138 (2020) 285–290.

[5] M. Lai, S. Skyrman, F. Kor, R. Homan, D. Babic, E. Edström, O. Persson, G. Burström, A. Elmi-Terander, B. H. W. Hendriks, P. H. N. De With, Development of a CT-compatible anthropomorphic skull phantom for surgical planning, training, and simulation, in: B. J. Park, T. M. Deserno (Eds.), Medical Imaging 2021: Imaging Informatics for Healthcare, Research, and Applications, SPIE, Online Only, United States, 2021, p. 9.

[6] E. Baldoin, G. Zullo, P. Carraretto, A. Koptyug, N. Petrone, Experimental evaluation of anti-rotational helmet technologies using a biofidelic human head replica embedding sensorised csf-meninges-brain simulants, 2024 IRCOBI Conf. Proc. (2024).

[7] R. Mantecón, B. Valverde-Marcos, I. Rubio, G. Youssef, J. A. Loya, J. Díaz-Álvarez, M. H. Miguélez, Additive Manufacturing of Head Surrogates for Evaluation of Protection in Sports, Polymers 16 (2024) 1753.

[8] F. Abayazid, K. Ding, K. Zimmerman, H. Stigson, M. Ghajari, A new assessment of bicycle helmets: The brain injury mitigation effects of new technologies in oblique impacts, Annals Biomed. Eng. 49 (2021) 2716–2733.

[9] S. Ouellet, M. Philippens, Correction to: The multi-modal responses of a physical head model subjected to various blast exposure conditions, Shock. Waves 28 (2018) 437–437.

[10] N. Petrone, G. Candiotto, E. Marzella, F. Uriati, G. Carraro, M. Bäckström, A. Koptyug, Feasibility of using a novel instrumented human head surrogate to measure helmet, head and brain kinematics and intracranial pressure during multidirectional impact tests, J. Sci. Medicine Sport 22 (2019) S78–S84.

[11] E. Baldoin, G. Zullo, A. Koptyug, N. Petrone, The role of additive manufacturing in the development of a biofidelic instrumented human head surrogate for impact tests, Transactions on Addit. Manuf. Meets Medicine Trans. AMMM 5 (2023) 10.

[12] B. Stone, S. Mitchell, Y. Miyazaki, N. Peirce, A. Harland, A destructible headform for the assessment of sports impacts, Proc. Inst. Mech. Eng. Part P: J. Sports Eng. Technol. 237 (2023) 7–18.

[13] E. Baldoin, G. Zullo, P. Carraretto, A. Koptyug, P. G. Pavan, N. Petrone, Development of a coupled experimental–numerical methodology using an instrumented human head replica for impact testing, Sports Eng. 28 (2025) 38.

[14] S. B. Hladky, M. A. Barrand, Mechanisms of fluid movement into, through and out of the brain: Evaluation of the evidence, Fluids Barriers CNS 11 (2014) 26.

[15] A. Goriely, M. G. Geers, G. A. Holzapfel, J. Jayamohan, A. Jérusalem, S. Sivaloganathan, W. Squier, J. A. van Dommelen, S. Waters, E. Kuhl, Mechanics of the brain: perspectives, challenges, and opportunities, Biomech. Model. Mechanobiol. 14 (2015) 931–965.

[16] G. Franceschini, D. Bigoni, P. Regitnig, G. A. Holzapfel, Brain tissue deforms similarly to filled elastomers and follows consolidation theory, J. Mech. Phys. Solids 54 (2006) 2592–2620.

[17] S. Budday, G. Sommer, C. Birkl, C. Langkammer, J. Haybaeck, J. Kohnert, M. Bauer, F. Paulsen, P. Steinmann, E. Kuhl, G. A. Holzapfel, Mechanical characterization of human brain tissue, Acta Biomater. 48 (2017) 319–340.

[18] A. E. Forte, S. Galvan, D. Dini, Models and tissue mimics for brain shift simulations, Biomech. Model. Mechanobiol. 17 (2018) 249–261.

[19] R. J. Waxweiler, D. Thurman, J. Sniezek, D. Sosin, J. O’Neil, Monitoring the impact of traumatic brain injury: A review and update, J. Neurotrauma 12 (1995) 509–516.

[20] S. Budday, T. C. Ovaert, G. A. Holzapfel, P. Steinmann, E. Kuhl, Fifty Shades of Brain: A Review on the Mechanical Testing and Modeling of Brain Tissue, Arch. Comput. Methods Eng. 27 (2020) 1187–1230.

[21] J. E. Galford, J. H. McElhaney, A viscoelastic study of scalp, brain, and dura, J. Biomech. 3 (1970) 211–221.

[22] M. S. Estes, J. H. McElhaney, Response of brain tissue to compressive loading, in: ASME Paper No. 70-WA/BHF-13, American Society of Mechanical Engineers, 1970.

[23] K. Miller, K. Chinzei, Constitutive modelling of brain tissue: Experiment and theory, J. Biomech. 30 (1997) 1115–1121.

[24] F. Shen, T. E. Tay, J. Z. Li, S. Nigen, P. V. S. Lee, H. K. Chan, Modified Bilston nonlinear viscoelastic model for finite element head injury studies, J. Biomech. Eng. 128 (2006) 797–801.

[25] S. Cheng, L. E. Bilston, Unconfined compression of white matter, J. Biomech. 40 (2007) 117–124.

[26] A. Tamura, S. Hayashi, I. Watanabe, K. Nagayama, T. Matsumoto, Mechanical characterization of brain tissue in high-rate compression, J. Biomech. Sci. Eng. 2 (2007) 115–126.

[27] B. Rashid, M. Destrade, M. D. Gilchrist, Mechanical characterization of brain tissue in compression at dynamic strain rates, J. Mech. Behav. Biomed. Mater. 10 (2012) 23–38.

[28] A. Shafiee, M. T. Ahmadian, M. Hoviattalab, Mechanical Characterization of Brain Tissue in Compression, in: Volume 3: 18th International Conference on Advanced Vehicle Technologies; 13th International Conference on Design Education; 9th Frontiers in Biomedical Devices, American Society of Mechanical Engineers, Charlotte, North Carolina, USA, 2016, p. V003T11A001.

[29] A. E. Forte, S. M. Gentleman, D. Dini, On the characterization of the heterogeneous mechanical response of human brain tissue, Biomech. Model. Mechanobiol. 16 (2017) 907–920.

[30] M. Hosseini-Farid, A. Rezaei, A. Eslaminejad, M. Ramzanpour, M. Ziejewski, G. Karami, Instantaneous and equilibrium responses of the brain tissue by stress relaxation and quasi-linear viscoelasticity theory, Sci. Iran. 26 (2019) 2047–2056.

[31] M. Navarro-Lozoya, M. S. Kennedy, D. Dean, J. I. Rodriguez-Devora, Development of phantom material that resembles compression properties of human brain tissue for training models, Materialia 8 (2019) 100438.

[32] K. Miller, K. Chinzei, Mechanical properties of brain tissue in tension, J. Biomech. 35 (2002) 483–490.

[33] F. Velardi, F. Fraternali, M. Angelillo, Anisotropic constitutive equations and experimental tensile behavior of brain tissue, Biomech. Model. Mechanobiol. 5 (2006) 53–61.

[34] A. Tamura, S. Hayashi, K. Nagayama,, T. Matsumoto, Mechanical characterization of brain tissue in high-rate extension, J. Biomech. Sci. Eng. 3 (2008) 263–274.

[35] B. Rashid, M. Destrade, M. D. Gilchrist, A high rate tension device for characterizing brain tissue, Proc. Inst. Mech. Eng. Part P: J. Sports Eng. Technol. 226 (2012) 170–176.

[36] W. Zhao, S. Ji, Brain strain uncertainty due to shape variation in and simplification of head angular velocity profiles, Biomech. Model. Mechanobiol. 16 (2017) 449–461.

[37] G. Fallenstein, V. Hulce, J. Melvin, Dynamic mechanical properties of human brain tissue, J. Biomech. 2 (1969) 217–226.

[38] B. R. Donnelly, J. Medige, Shear properties of human brain tissue, J. Biomech. Eng. 119 (1997) 423–432.

[39] L. E. Bilston, Z. Liu, N. Phan-Thien, Large strain behaviour of brain tissue in shear: some experimental data and differential constitutive model, Biorheology 38 (2001) 335–345.

[40] E. G. Takhounts, J. R. Crandall, K. Darvish, On the importance of nonlinearity of brain tissue under large deformations, in: 47th Stapp Car Crash Conference (2003), 2003, pp. 2003–22–0005.

[41] M. Hrapko, J. A. Van Dommelen, G. W. Peters, J. S. Wismans, The mechanical behaviour of brain tissue: Large strain response and constitutive modelling, Biorheology 43 (2006) 59–69.

[42] M. Destrade, M. D. Gilchrist, J. G. Murphy, B. Rashid, G. Saccomandi, Extreme softness of brain matter in simple shear, Int. J. Non-Linear Mech. 75 (2015) 54–58.

[43] S. Chatelin, A. Constantinesco, R. Willinger, Fifty years of brain tissue mechanical testing: From in vitro to in vivo investigations, Biorheol. The Off. J. Int. Soc. Biorheol. 47 (2010) 255–276.

[44] R. Moran, J. H. Smith, J. J. García, Fitted hyperelastic parameters for human brain tissue from reported tension, compression, and shear tests, J. Biomech. 47 (2014) 3762–3766.

[45] L. Mihai, L. Chin, P. A. Janmey, A. Goriely, A comparison of hyperelastic constitutive models applicable to brain and fat tissues, J. Royal Soc. Interface 12 (2015) 1–12.

[46] G. Z. Voyiadjis, A. Samadi-Dooki, Hyperelastic modeling of the human brain tissue: Effects of no-slip boundary condition and compressibility on the uniaxial deformation, J. Mech. Behav. Biomed. Mater. 83 (2018) 63–78.

[47] M. J. Lohr, G. P. Sugerman, S. Kakaletsis, E. Lejeune, M. K. Rausch, An introduction to the Ogden model in biomechanics: Benefits, implementation tools and limitations, Philos. Transactions Royal Soc. A: Math. Phys. Eng. Sci. 380 (2022) 20210365.

[48] A. Anssari-Benam, M. Destrade, G. Saccomandi, Modelling brain tissue elasticity with the Ogden model and an alternative family of constitutive models, Philos. Transactions Royal Soc. A: Math. Phys. Eng. Sci. 380 (2022) 20210325.

[49] J. Hinrichsen, N. Reiter, L. Bräuer, F. Paulsen, S. Kaessmair, S. Budday, Inverse identification of regionspecific hyperelastic material parameters for human brain tissue, Biomech. Model. Mechanobiol. 22 (2023) 1729–1749.

[50] S. R. St. Pierre, K. Linka, E. Kuhl, Principal-stretch-based constitutive neural networks autonomously discover a subclass of Ogden models for human brain tissue, Brain Multiphysics 4 (2023) 100066.

[51] N. Filla, J. Hou, T. Liu, S. Budday, X. Wang, Accuracy meets simplicity: A constitutive model for heterogenous brain tissue, J. Mech. Behav. Biomed. Mater. 150 (2024) 106271.

[52] F. Bogoni, M. P. Wollner, G. A. Holzapfel, On the experimental identification of equilibrium relations and the separation of inelastic effects in soft biological tissues, J. Mech. Phys. Solids 193 (2024) 105868.

[53] M. P. Wollner, G. A. Holzapfel, The Mullins effect, mechanical preconditioning, and predictive power, J. Mech. Phys. Solids 215 (2026) 106698.

[54] A. Lion, A constitutive model for carbon black filled rubber: Experimental investigations and mathematical representation, Continuum Mech. Thermodyn. 8 (1996) 153–169.

[55] J. Bergström, Constitutive modeling of the large strain time-dependent behavior of elastomers, J. Mech. Phys. Solids 46 (1998) 931–954.

[56] J. Bergström, M. Boyce, Large strain time-dependent behavior of filled elastomers, Mech. Mater. 32 (2000) 627–644.

[57] C. Miehe, J. Keck, Superimposed finite elastic–viscoelastic–plastoelastic stress response with damage in filled rubbery polymers. Experiments, modelling and algorithmic implementation, J. Mech. Phys. Solids 48 (2000) 323–365.

[58] P. Haupt, K. Sedlan, Viscoplasticity of elastomeric materials: Experimental facts and constitutive modelling, Arch. Appl. Mech. (Ingenieur Arch. 71 (2001) 89–109.

[59] A. S. Khan, O. Lopez-Pamies, Time and temperature dependent response and relaxation of a soft polymer, Int. J. Plast. 18 (2002) 1359–1372.

[60] A. Amin, M. Alam, Y. Okui, An improved hyperelasticity relation in modeling viscoelasticity response of natural and high damping rubbers in compression: Experiments, parameter identification and numerical verification, Mech. Mater. 34 (2002) 75–95.

[61] J. Diani, M. Brieu, J. M. Vacherand, A damage directional constitutive model for mullins effect with permanent set and induced anisotropy, Eur. J. Mech. A/Solids 25 (2006) 483–496.

[62] A. Vandenbroucke, H. Laurent, N. A. Hocine, G. Rio, A hyperelasto-visco-hysteresis model for an elastomeric behaviour: Experimental and numerical investigations, Comput. Mater. Sci. 48 (2010) 495–503.

[63] H. Laurent, A. Vandenbroucke, G. Rio, N. A. Hocine, A simplified methodology to identify material parameters of a hyperelasto-visco-hysteresis model: application to a fluoro-elastomer, Model. Simul. Mater. Sci. Eng. 19 (2011) 085004.

[64] M. Hossain, D. K. Vu, P. Steinmann, Experimental study and numerical modelling of VHB 4910 polymer, Comput. Mater. Sci. 59 (2012) 65–74.

[65] M. Kästner, M. Obst, J. Brummund, K. Thielsch, V. Ulbricht, Inelastic material behavior of polymers experimental characterization, formulation and implementation of a material model, Mech. Mater. 52 (2012) 40–57.

[66] Y. Mao, S. Lin, X. Zhao, L. Anand, A large deformation viscoelastic model for double-network hydrogels, J. Mech. Phys. Solids 100 (2017) 103–130.

[67] S. Wang, S. A. Chester, Experimental characterization and continuum modeling of inelasticity in filled rubber-like materials, Int. J. Solids Struct. 136-137 (2018) 125–136.

[68] M. Hossain, R. Navaratne, D. Perić, 3D printed elastomeric polyurethane: Viscoelastic experimental characterizations and constitutive modelling with nonlinear viscosity functions, Int. J. Non-Linear Mech. 126 (2020) 103546.

[69] K. Alkhoury, R. Ivko, M. Hossain, S. Nadimpalli, S. A. Chester, Experiments and modeling of the coupled viscoelasticity and Mullins effect in filled rubber materials, J. Mech. Phys. Solids 188 (2024) 105650.

[70] F. Bogoni, M. P. Wollner, G. A. Holzapfel, Experimental investigation on the rate dependence of the equilibrium relations in aortic tissue, J. Mech. Phys. Solids 211 (2026) 106547.

[71] S. Pettenuzzo, M. Terzano, M. Todesco, M. Casarin, E. Belluzzi, P. Ruggieri, E. L. Carniel, G. A. Holzapfel, C. G. Fontanella, Histological analysis, viscoelastic characterization, and modeling of human plantar fascia, Acta Biomater. 217 (2026) 488–502.

[72] D. Singh, S. Boakye-Yiadom, D. S. Cronin, Comparison of porcine brain mechanical properties to potential tissue simulant materials in quasi-static and sinusoidal compression, J. Biomech. 92 (2019) 84–91.

[73] T. Distler, E. Schaller, P. Steinmann, A. Boccaccini, S. Budday, Alginate-based hydrogels show the same complex mechanical behavior as brain tissue, J. Mech. Behav. Biomed. Mater. 111 (2020) 103979.

[74] M. P. Kainz, A. Greiner, J. Hinrichsen, D. Kolb, E. Comellas, P. Steinmann, S. Budday, M. Terzano, G. A. Holzapfel, Poro-viscoelastic material parameter identification of brain tissue-mimicking hydrogels, Front. Bioeng. Biotechnol. 11 (2023) 1143304.

[75] P. Haupt, Continuum Mechanics and Theory of Materials, Springer Berlin Heidelberg, Heidelberg, 2002.

[76] C. Truesdell, Rational Thermodynamics, Springer New York, New York, NY, 1984.

[77] C. G. Armstrong, W. M. Lai, V. C. Mow, An Analysis of the Unconfined Compression of Articular Cartilage, J. Biomech. Eng. 106 (1984) 165–173.

[78] R. W. Ogden, Non-Linear Elastic Deformations, Dover Publications, New York, 1997.

[79] W. Ehlers, J. Bluhm (Eds.), Porous Media, pnSpringer Berlin Heidelberg, Berlin, Heidelberg, 2002.

[80] R. Hill, On constitutive macro-variables for heterogeneous solids at finite strain, Proc. Royal Soc. London. A. Math. Phys. Sci. 326 (1972) 131–147.

[81] G. Sommer, T. C. Gasser, P. Regitnig, M. Auer, G. A. Holzapfel, Dissection properties of the human aortic media: An experimental study, J. Biomech. Eng. 130 (2008) 021007.

[82] G. Sommer, M. Eder, L. Kovacs, H. Pathak, L. Bonitz, C. Mueller, P. Regitnig, G. A. Holzapfel, Multiaxial mechanical properties and constitutive modeling of human adipose tissue: A basis for preoperative simulations in plastic and reconstructive surgery, Acta Biomater. 9 (2013) 9036–9048.

[83] M. Terzano, A. Spagnoli, D. Dini, A. E. Forte, Fluid–solid interaction in the rate-dependent failure of brain tissue and biomimicking gels, J. Mech. Behav. Biomed. Mater. 119 (2021) 104530.

[84] L. Su, M. Wang, J. Yin, F. Ti, J. Yang, C. Ma, S. Liu, T. J. Lu, Distinguishing poroelasticity and viscoelasticity of brain tissue with time scale, Acta Biomater. 155 (2023) 423–435.

[85] T. Yuan, W. Zhan, M. Terzano, G. A. Holzapfel, D. Dini, A comprehensive review on modeling aspects of infusion-based drug delivery in the brain, Acta Biomater. 185 (2024) 1–23.

[86] M. P. Kainz, M. Terzano, D. Kolb, G. A. Holzapfel, Effective porosity and fluid flow in macroporous ultrasoft hydrogels: An experimental characterization, Acta Biomater. 217 (2026) 458–474.

[87] R. M. Bowen, Theory of Mixtures, in: Continuum Physics, Elsevier, 1976, pp. 1–127.

[88] B. Markert, A constitutive approach to 3-d nonlinear fluid flow through finite deformable porous continua, Transp. Porous Media 70 (2007) 427–450.

[89] W. Ehlers, G. Eipper, Finite Elastic Deformations in Liquid-Saturated and Empty Porous Solids, in: Porous Media: Theory and Experiments, volume 34, Springer Netherlands, Dordrecht, 1999, pp. 179–191.

[90] G. A. Holzapfel, Nonlinear Solid Mechanics: A Continuum Approach for Engineering Science, Wiley, Chichester, 2000.

## Supplementary references

[S1] J. Faber, J. Hinrichsen, A. Greiner, N. Reiter, S. Budday, Tissue-scale biomechanical testing of brain tissue for the calibration of nonlinear material models, Curr. Protoc. 2 (2022).

[S2] N. Reiter, B. Roy, F. Paulsen, S. Budday, Insights into the microstructural origin of brain viscoelasticity: Prospects for microstructure-informed constitutive modeling, J. Elast. 145 (2021) 99–116.

[S3] F. Eskandari, M. Shafieian, M. M. Aghdam, K. Laksari, Tension strain-softening and compression strain-stiffening behavior of brain white matter, Annals Biomed. Eng. 49 (2021) 276–286.

[S4] M. Prange, D. F. Meaney, Defining Brain Mechanical Properties: Effects of Region, Direction, and Species, in: 44th Stapp Car Crash Conference (2000), 2000, pp. 2000–01–SC15.

[S5] S. Cheng, L. E. Bilston, Unconfined compression of white matter, J. Biomech. 40 (2007) 117–124.

[S6] M. Hrapko, J. A. V. Dommelen, G. W. Peters, J. S. Wismans, Characterisation of the mechanical behaviour of brain tissue in compression and shear, Biorheology 45 (2008) 663–676.

[S7] A. Tamura, S. Hayashi, K. Nagayama,, T. Matsumoto, Mechanical characterization of brain tissue in high-rate extension, J. Biomech. Sci. Eng. 3 (2008) 263–274.

[S8] K. Laksari, M. Shafieian, K. Darvish, Constitutive model for brain tissue under finite compression, J. Biomech. 45 (2012) 642–646.

[S9] B. Rashid, M. Destrade, M. D. Gilchrist, Mechanical characterization of brain tissue in compression at dynamic strain rates, J. Mech. Behav. Biomed. Mater. 10 (2012) 23–38.

[S10] B. Rashid, M. Destrade, M. D. Gilchrist, A high rate tension device for characterizing brain tissue, Proc. Inst. Mech. Eng. Part P: J. Sports Eng. Technol. 226 (2012) 170–176.

[S11] M. Laksari, K. Darvish, K. Crandall, J. Shafieian, Development of a constitutive model for brain tissue under multiaxial loading, 2012 IRCOBI Conf. Proc. - Int. Res. Counc. on Biomech. Inj. (2012).

[S12] S. Javid, A. Rezaei, G. Karami, A micromechanical procedure for viscoelastic characterization of the axons and ecm of the brainstem, J. Mech. Behav. Biomed. Mater. 30 (2014) 290–299.

[S13] A. Shafiee, M. T. Ahmadian, M. Hoviattalab, Mechanical Characterization of Brain Tissue in Compression, in: Volume 3: 18th International Conference on Advanced Vehicle Technologies; 13th International Conference on Design Education; 9th Frontiers in Biomedical Devices, American Society of Mechanical Engineers, Charlotte, North Carolina, USA, 2016, p. V003T11A001.

[S14] S. Budday, G. Sommer, C. Birkl, C. Langkammer, J. Haybaeck, J. Kohnert, M. Bauer, F. Paulsen, P. Steinmann, E. Kuhl, G. A. Holzapfel, Mechanical characterization of human brain tissue, Acta Biomater. 48 (2017) 319–340.

[S15] A. E. Forte, S. M. Gentleman, D. Dini, On the characterization of the heterogeneous mechanical response of human brain tissue, Biomech. Model. Mechanobiol. 16 (2017) 907–920.

[S16] M. Hosseini-Farid, A. Rezaei, A. Eslaminejad, M. Ramzanpour, M. Ziejewski, G. Karami, Instantaneous and equilibrium responses of the brain tissue by stress relaxation and quasi-linear viscoelasticity theory, Sci. Iran. 26 (2019) 2047–2056.

[S17] T. Distler, E. Schaller, P. Steinmann, A. Boccaccini, S. Budday, Alginate-based hydrogels show the same complex mechanical behavior as brain tissue, J. Mech. Behav. Biomed. Mater. 111 (2020) 103979.

[S18] A. Greiner, N. Reiter, F. Paulsen, G. A. Holzapfel, P. Steinmann, E. Comellas, S. Budday, Poro-viscoelastic effects during biomechanical testing of human brain tissue, Front. Mech. Eng. 7 (2021).

